# Variation in colour signals among *Sarracenia* pitcher plants and the potential role of areoles in the attraction of flying Hymenoptera

**DOI:** 10.1101/2021.09.15.460199

**Authors:** Corentin Dupont, Claire Villemant, Tom Hatterman, Jeremie Pratviel, Laurence Gaume, Doris Gomez

**Author notes:** co-last authors (Laurence Gaume, Doris Gomez).

## Abstract

*Sarracenia* insectivorous plants show a diversity of visual features in their pitchers but their perception by insects and their role in attraction, have received little attention. They also vary in prey composition, with some species trapping more flying Hymenoptera, such as bees. To test the hypothesis of a link between visual signal variability and prey segregation ability, and to identify which signal could attract flying Hymenoptera, we characterised, the colour patterns of 32 pitchers belonging to four taxa, modelled their perception by flying Hymenoptera, and examined the prey they trapped. The pitchers of the four taxa differed in colour patterns, with notably two long-leaved taxa displaying clear areoles, which contrasted strongly in colour and brightness with the vegetative background and with other pitcher areas in the eyes of flying Hymenoptera. These taxa trapped high proportion of flying hymenoptera. This suggests that contrasting areoles may act as a visual lure for flying Hymenoptera, making plants particularly visible to these insects. Prey capture also differed according to pitcher stage, morphology, season and visual characteristics. Further studies on prey visitation are needed to better understand the link between prey capture and attraction feature.

## INTRODUCTION

Angiosperms, the most diverse group of plants, underwent explosive radiation during the Cretaceous in a coevolutionary process with insect pollinators (Grimaldi, 1999; Van der Niet & Johnson, 2012). These flowering plants have thus evolved different combinations of nectar rewards, and sensory signals, such as visual and olfactory cues, which attract a diversity of pollinators (Kevan & Baker, 1983; Dobson, 1994; Lunau & Maier, 1995; Chittka & Raine, 2006). Visual and olfactory signals often operate synergistically in attracting pollinators (Raguso, 2008). Some plants, such as orchid species, have even evolved ‘dishonest’ signals that deceive pollinators, by exploiting their sensory biases (Schaefer & Ruxton, 2009) and/or by using deceit mimicry of nectar or sexual mate (Dafni, 1984; Schiestl, 2010), without any reward. Carnivorous plants are particularly efficient at exploiting the sensory biases of generalist insect-pollinators because their leaves often resemble flowers in their visual and olfactory traits such as colour patterns (Moran *et al*., 1999; Schaefer & Ruxton, 2008; Moran *et al*., 2012b), even in the UV range (Joel *et al*., 1985; Moran, 1996; Kurup *et al*., 2013; Golos, 2020), nectar guides (Dress *et al*., 1997; Bennett & Ellison, 2009), and perfume (Jaffé *et al*., 1995; Jürgens *et al*., 2009; Di Giusto *et al*., 2010). The most adorned plants regarding these floral features attract a wide range of flower-visiting insects (Joel, 1988; Di Giusto *et al*., 2010). Part of the attracted insects are then trapped with an arsenal of leaf traits, as diverse as snap-traps (Forterre *et al*., 2005), viscoelastic mucilage (Gaume & Forterre, 2007; Adlassnig *et al*., 2010) and slippery surfaces (Gaume *et al*., 2004; Bauer *et al*., 2013). They are digested by endogenous and/or exogenous proteolytic enzymes, and ultimately supplement carnivorous plants with essential nutrients, which often lack in the soil where they grow (Ellison & Adamec, 2018).

One group of carnivorous plants, the so-called pitcher plants, includes the well-known Sarraceniaceae from the Americas and the Nepenthaceae from Southeast Asia. These plants exhibit several morphological and physiological adaptations, such as pitcher-like leaves of different sizes and shapes (Cresswell, 1993; Bhattarai & Horner, 2009; Gaume *et al*., 2016), different attractive visual or olfactory signals (Moran *et al*., 1999; Di Giusto *et al*., 2010) and capture devices (Bonhomme *et al*., 2011). However, it is difficult to disentangle the contribution of the different features since they are often combined to form whole trapping syndromes, which target specific guilds of animals or characterise specific diets (Pavlovič *et al*., 2007; Gaume *et al*., 2016).

Compared to olfactory cues (Jaffé *et al*., 1995; Jürgens *et al*., 2009; Di Giusto *et al*., 2010), the visual cues of pitcher plants as perceived by insects have received no attention to our knowledge and their role in attraction is the subject of controversy. For instance, pitcher red colouration has been proposed to play an important role in prey attraction by some authors (Newell & Nastase, 1998; Schaefer & Ruxton, 2008) but not by others (Green & Horner, 2007). In an experiment with *Sarracenia purpurea* (Linnaeus), extrafloral nectar, often associated to red colouration in carnivorous plants (Bennett & Ellison, 2009; Gaume *et al.*, 2016), was shown to account for prey attraction, rather than the red colouration itself (Bennett & Ellison, 2009). Yet, colour signals should not be overlooked since they operate at larger distances than nectar and are thus needed to attract flying insects. Moreover, many pitcher plants sport a large colour diversity with contrasting patterns and not just bicolor patterns with nuances of green and red (Moran *et al*., 1999), which suggests that visual signals are important components of the attractive system in these plants. For instance, clear white areoles present in some carnivorous plants (Schnell, 2002; McPherson, 2006) have been shown to function as light lures helping insect capture in *Sarracenia minor* (Walter) (McGregor *et al*., 2016), and in the Sumatran *Nepenthes aristolochioides* (Jebb & Cheek) (Moran *et al*., 2012a). Yet, according to Schaefer & Ruxton (2014), these areoles only play a role in attracting insects to pitcher plants, but have no role in prey capture. The role, if any, of clear areoles in attracting prey – a prerequisite for capture in carnivorous plants – remains to be clarified. As visual contrasts are important features of attraction in pollinating systems (Spaethe *et al*., 2001), we hypothesise that clear areoles are better perceived by flying prey since they are more conspicuous against the green vegetation background and the remaining parts of the plant. We test this hypothesis by exploring the variation in colouration in four *Sarracenia* pitcher plants of different pitcher morphology and visual characteristics, especially regarding the areoles, and by modelling how these colours are perceived by flying Hymenoptera. We secondly investigated whether clear areoles play a role in attracting this kind of insects to the pitcher by examining the relationship between the composition in prey trapped by pitchers and the visual signals they displayed, as seen by flying Hymenoptera.

## MATERIAL AND METHODS

### PLANT TAXA AND GROWING CONDITIONS

We considered four *Sarracenia* taxa differing in the visual aspect and leaf shape (Fig. 1). Two were natural species: *S. purpurea* subsp. *venosa* (Rafinesque) and *S.* × *mitchelliana* = *S. purpurea* × *S. leucophylla* hybrid. *S. purpurea,* which does not bear clear areoles, mostly traps ants and comparatively few flying hymenopterans (Cresswell, 1991; Heard, 1998; Newell & Nastase, 1998; Bhattarai & Horner, 2009). The two others were horticultural hybrids with clear areoles: *S.* × Juthatip soper = (*S. leucophylla* × *S. purpurea*) × *S. leucophylla* and *S.* × *leucophylla*=*S. leucophylla* × *S.* × Juthatip soper. The long-leaved *Sarracenia leucophylla* (Rafinesque) characterized by clear areoles has been observed to trap a large amount of bees in the field (Gibson, 1983a) and its horticultural hybrids have been observed to trap wasps and hornets in large amounts (Meurgey & Perrocheau, 2015). Plants were grown in polyculture in a 6 m^2^ container filled with a peat mixture consisting of 2/3 blond peat and 1/3 sand on a background of clay balls. Containers were placed outdoors in the experimental station of AMAP (Montpellier, France) in a sunny place. They were regularly watered with demineralized water to match their natural moist and mineral-poor environment.

**Figure 1.**
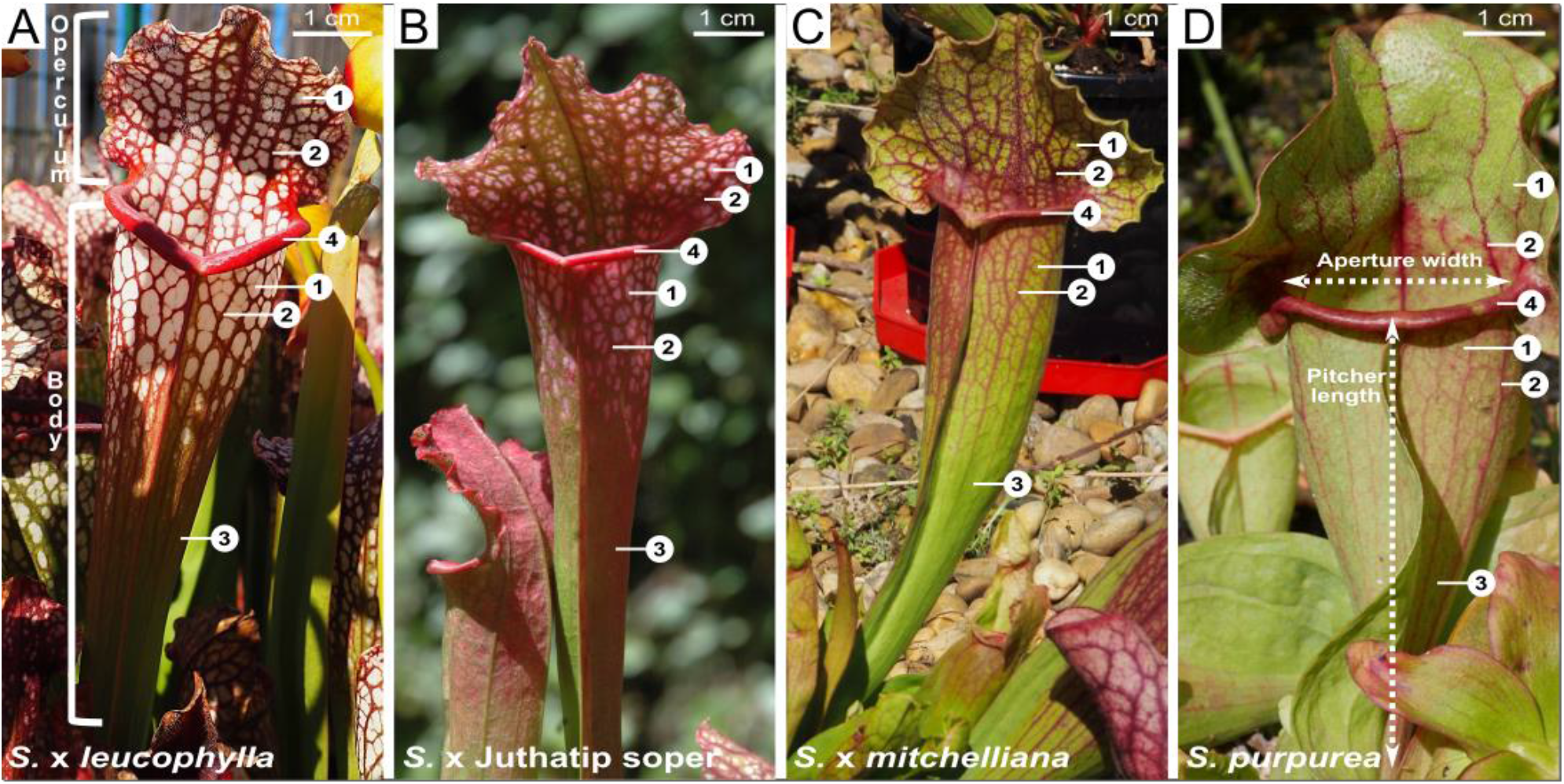
Pitchers of the four *Sarracenia* taxa: (A) *S.* × *leucophylla,* (B) *S.* × Juthatip soper, (C) *S*. × *mitchelliana,* (D) *S. purpurea* and their different areas: areoles (1) and veins (2) on the operculum, areoles (1), veins (2) and tube (3) on the pitcher body, and the peristome (4). For all species, areoles are defined as the areas between the veins, in the pitcher body or in the operculum. Areoles appear white (A), pink (B), or green (C, D) to a human eye. Aperture width and pitcher length, shown in (D) were measured in all pitchers.

### PITCHER MORPHOLOGICAL MEASUREMENTS AND PREY CAPTURE ANALYSIS

Pitchers were marked the day they opened and four pitcher stages were thus scored (1: young, 2: midle, 3: mature, 4: old) corresponding to 5±1, 15±3, 40±5, and 60±10 days after pitcher opening, respectively. We measured a total of 32 pitchers on 21 plants (8 pitchers per taxon). We took a first set of measurements for 24 pitchers of different stages belonging to 5-6 plants of each of the four taxa in mid-August 2016 (summer), and a second set of 8 pitchers of stage 2 belonging to two plants of each of the four taxa in October 2016 (autumn). The plants measured in autumn were different from those measured in summer. We measured pitcher length and aperture width (see Fig. 1D) before taking colour measurements (see below). These measurements were compared among taxa using one-way ANOVAs. We then collected pitcher content and preserved it in 70% ethanol. Claire Villemant identified prey individuals to the species level when possible, and at least always to the family or super-family level, depending on the stage of the digestive process. It was always possible to make the distinction between flying Hymenoptera (e. g. social or solitary bees and wasps, hornets, parasitoid wasps and sawflies) and crawling Hymenoptera (ants).

### PITCHER COLOUR MEASUREMENT AND ANALYSIS

To study the visual signals of the carnivorous plants, we measured the reflectance spectra of the 32 pitchers using spectrophotometry. The light produced by a Deuterium-Halogen light source (Avalight) was conducted to the sample through an optic probe (FCR-7UV200-2-45-ME, terminated by a quartz transparent window cut at 45° guaranteeing a constant distance between the sample and the light beam). The light reflected by the sample was conducted through the probe to a spectrometer (Ocean Optics USB 2000). Measurements were taken between 300 and 700 nm, which included the range of flying Hymenoptera sensitivity, considered to be between 300 nm and 650 nm, as in honeybees (De Ibarra *et al*., 2014). In each of the 32 pitchers, we took 4 to 5 repeated spectral measurements of the following six areas: areoles of the body, veins of the body, peristome, areoles of the operculum, veins of the operculum and tube (Fig. 1).

*Sarracenia* pitcher plants trap mainly bees, wasps and hornets as flying Hymenoptera (Gibson, 1983a; Farnsworth & Ellison, 2008; Meurgey & Perrocheau, 2015), and all have similar trichromatic vision (Peitsch *et al*., 1992) with similar spectral sensitivity (Peitsch *et al*., 1992; Briscoe & Chittka, 2001; De Ibarra *et al*., 2014). Hence, we chose the hornet *Vespa crabro* (Linnaeus) to model how *Sarracenia* colours are perceived by flying Hymenoptera, using Vorobyev & Osorio’s discriminability model (Vorobyev & Osorio, 1998) to compute the visual contrasts produced by a pitcher colour seen against a green grass background or against another pitcher colour. We took the D65 standard illuminant as the irradiance spectrum of an open habitat, the reflectance spectrum of a colour area of a *Sarracenia* pitcher plant, the mean reflectance spectrum of green grass as a visual background (Gomez, personal data) for whenever the contrast against the background was computed. We built flying Hymenoptera photoreceptor absorptance curves by using photoreceptor peaks measured for hornet by Peitsch *et al.* (1992), that is 336, 436 and 536 nm for the S, M, and L photoreceptors respectively. We assumed a neural noise and ωi-values of 0.13, 0.06 and 0.12 for S, M, and L receptors respectively, as in Vorobyev & Osorio (1998). We assumed that brightness was detected by L receptors (Lehrer, 1993).

We first computed the colour and brightness contrasts displayed by the various pitcher areas against the mean green grass background. These contrasts are used to quantify how visible the pitcher plant stands out against its natural background, and has been used to better understand pollinator attraction to plants, as in orchids for instance (Streinzer *et al*., 2009; Streinzer *et al*., 2010). These authors computed colour and brightness contrasts to understand attraction at short and long distances respectively, as Hymenoptera use colour signal at close range and brightness signal to detect small objects or objets at larger distances, respectively (Giurfa *et al.*, 1996; Dyer *et al.*, 2008). Second, we computed the colour and brightness contrasts between any two pitcher areas. Aguiar *et al.* (2020) considered an equivalent of colour contrast against the background of sepals, petals and labellum of tropical orchids pollinated by bees, but also the colour contrast between each floral piece and showed the importance of intrafloral colour patterns in pollinator perception, which could serve as a guide to the flower centre. Colour patterns between pitcher areas could similarly help attracting prey to the trap (Moran, 1996). Regarding pitcher areas, areoles of the pitcher body and areoles of the operculum had similar colouration and yielded similar contrasts, they were thus pooled into a common ‘areoles’ category. Likewise, the veins of the pitcher body and the veins of the operculum had similar colouration and yielded similar contrasts, and were pooled into a common ‘veins’ category. Hence the variable pitcher area had four levels: areoles, veins, peristome, and tube. All computations were performed with the R package pavo version 2 (Maia *et al.*, 2019).

### STATISTICAL ANALYSES

We analysed variation in plant colouration and variation in prey capture using a mixed model approach as it is well suited to repeated observations. All statistics were performed using the software R version 4.0.3 (R Core Team, 2020). Model assumptions were checked by plotting residuals versus fitted values.

First, we explored to which extent the colour contrast and the brightness contrast displayed by a pitcher area against the green background varied with plant taxon, pitcher stage and pitcher area. We took the colour contrast or the brightness contrast of all spectral measurements as the dependent variable, a normal error distribution, and Plant identity and Pitcher identity nested within Plant identity as random effects. We tested plant taxon, pitcher stage, season, pitcher area, and all relevant two-way and three-way interactions when possible (for season, only the simple effect could be included). Both dependent variables were square-root transformed to match residual normality.

Second, considering the contrasts between any two pitcher areas, we explored to which extent the colour contrast and the brightness contrast displayed by a pitcher area against another varied with plant taxon, pitcher stage and the pitcher area considered. We took the colour contrasts or the brightness contrasts between pitcher areas as the dependent variable, a normal error distribution, and Plant identity and Pitcher identity nested within Plant identity as random effects. We tested plant taxon, pitcher stage, season, pair of areas (each combination of two areas contrasted with each other), and all relevant two-way and three-way interactions when possible (for season, only the simple effect could be included). Both dependent variables were square-root transformed to match residual normality.

Finally, we tested whether the number of prey trapped was linked to plant taxon, pitcher stage, pitcher morphology and season. The total number of prey individuals and the total number of crawling Hymenoptera trapped were analysed using Generalized Linear Models (GLM) assuming a Poisson distribution with a log-link function. We tested, as explanatory variables, Plant taxon, Pitcher Stage, Pitcher length, Aperture width, the ratio Pitcher length / Aperture width, and Capture area (as defined in Bhattarai & Horner (2009) i.e. the area of a circle of diameter Aperture width) and Season. For morphological variables, which were partly correlated (for instance the ratio Pitcher length / Aperture width was negatively correlated with Pitcher length : correlation-coefficient = −0.83, t = −8.27, p-value < 0.001), we chose to include in the same model only the least correlated variables, that is Pitcher length and Aperture width (which were not correlated : correlation-coefficient = −0.20, t = −1.15, p-value = 0.261). For this analysis, an outlier was identified and excluded (a *S.* × *leucophylla* pitcher with 153 prey individuals including 54 ants).

Furthermore, we tested whether the number of flying Hymenoptera prey trapped was also linked to visual signals. The number of flying Hymenoptera was analysed using similar Poisson regression models. As Plant taxon and Pitcher stage absorbed the variation in visual signals, they were not included in the analysis. Season was retained as it was likely to affect the abundance of prey available. As the visual signals measured were numerous compared to the sample size, we focused on the variables of interest, i.e. the contrasts related to areoles and also included in the analysis only the least correlated contrasts (<50%, see Supporting Information Fig. S1). Hence, we tested the effects of Season, Pitcher length, Aperture width, Areoles brightness contrast with background, Areoles-Peristome colour contrast, Areoles-Peristome brightness contrast, Areoles-Tube colour contrast.

For all models, we selected the best model with the lowest AIC, and we did that for the fixed and random effects separately. In all mixed models, only Pitcher identity was retained as a random effect. Backward selection of models and type III tests were carried out. For each significant factor, we estimated the regression coefficients and post-hoc tests were carried out between any two factor levels, when necessary.

## RESULTS

### PITCHER MORPHOLOGY AND COLOURATION

Plant taxa differed both in morphology and colouration (Fig. 1, Supporting Information Fig. S2, Fig. S3). *S.* × *leucophylla* and *S.* × Juthatip soper tended to have long pitchers with a narrow aperture (Fig. S3) and well-defined clear areoles, which respectively appeared white and pink to a human eye (Fig. 1, Fig. S3). The other two taxa, *S.* × *mitchelliana* and *S. purpurea*, had short pitchers with a large aperture (Fig. S3), and areas defined as areoles had a green colouration to a human eye, which was similar to that of the pitcher tube (Fig. 1, Fig. S2). In all taxa, the tube generally reflected light in the middle wavelengths while the peristome and veins reflected more in the long wavelengths, appearing respectively green and red to a human eye (Fig. 1, Fig. S2).

### CONTRASTS WITH A GREEN BACKGROUND

The visual contrasts – colour contrast and brightness contrast – produced by pitcher areas against a green background as seen by flying Hymenoptera varied with taxon and stage (Table 1). More specifically, to flying Hymenoptera, peristome and tube showed no significant difference in colour contrast between taxa (Fig. 2A, (Supporting Information Table S1), and veins and tube showed no significant difference in brightness contrast between taxa (Fig. 2B, Supporting Information Table S2). Conversely, areoles and veins differed in colour contrast and areoles and peristome differed in brightness contrast between taxa (Fig. 2, Table S1, S2). The veins contrasted more in colour with a green background in S. × *mitchelliana* than in *S.* × *leucophylla*. The peristome contrasted more in brightness with a green background in *S.* × *leucophylla* than in *S.* × Juthatip soper. The areoles contrasted more in colour with a green background in *S.* x *leucophylla* and S. x Juthatip soper than in *S.* x *mitchelliana* and *S. purpurea* while *S.* × *leucophylla* had the most contrasting areoles of all four taxa. Hence, areoles accounted for most of the visual differences observed between taxa as they differed both in colour and brightness among taxa (Fig. 2). Yet, the area that contrasted most strongly against a green background was not always the same from one taxon to another (Table S1, S2): for the two long-leaved taxa, *S.* × *leucophylla* and *S.* × Juthatip soper, areoles were the most contrasting area either for colour or for brightness (Table S1, S2). Conversely, the most contrasting area for colour was the veins for *S.* × *mitchelliana* while it was the peristome for *S. purpurea* (Table S1, S2).

**Table 1.** Factors of variation in colour and brightness contrasts with a green background. Only the best models are presented. Both dependent variables were square-root transformed to match residuals normality. “Chisq” values refer to type III Wald chi-square tests. Symbols describe various levels of p-values *: p<0.05, **: p<0.01, and ***: p<0.001. Contrasts are detailed in ESM, Tables S1 (colour contrast) and S2 (brightness contrast). “ - ” means that the variable was not retained in the best model. “ : ” refers to the interaction between variables.

| Explanatory variables | Colour contrast |  |  |  | Brightness contrast |  |  |  |
| --- | --- | --- | --- | --- | --- | --- | --- | --- |
|  | Chisq | Df | p-value |  | Chisq | Df | p-value |  |
| Intercept | 80.11 | 1 | <0.001 | *** | 74.33 | 1 | <0.001 | *** |
| Plant taxon | 9.69 | 3 | 0.0214 | * | 1.38 | 3 | 0.7107 |  |
| Area | 32.28 | 3 | <0.001 | *** | 15.68 | 3 | 0.0013 | ** |
| Pitcher stage | 3.06 | 3 | 0.3823 |  | 5.27 | 3 | 0.1532 |  |
| Season | 31.79 | 1 | <0.001 | *** | - | - | - |  |
| Plant taxon: Area | 72.44 | 9 | <0.001 | *** | 26.96 | 9 | 0.0014 | ** |
| Plant taxon: Pitcher stage | 11.20 | 9 | 0.2622 |  | 17.55 | 9 | 0.0407 | * |
| Area: Pitcher stage | 45.83 | 9 | <0.001 | *** | 40.48 | 9 | <0.001 | *** |
| Plant taxon: Area : Pitcher stage | 103.08 | 27 | <0.001 | *** | 136.96 | 27 | <0.001 | *** |

**Figure 2.**
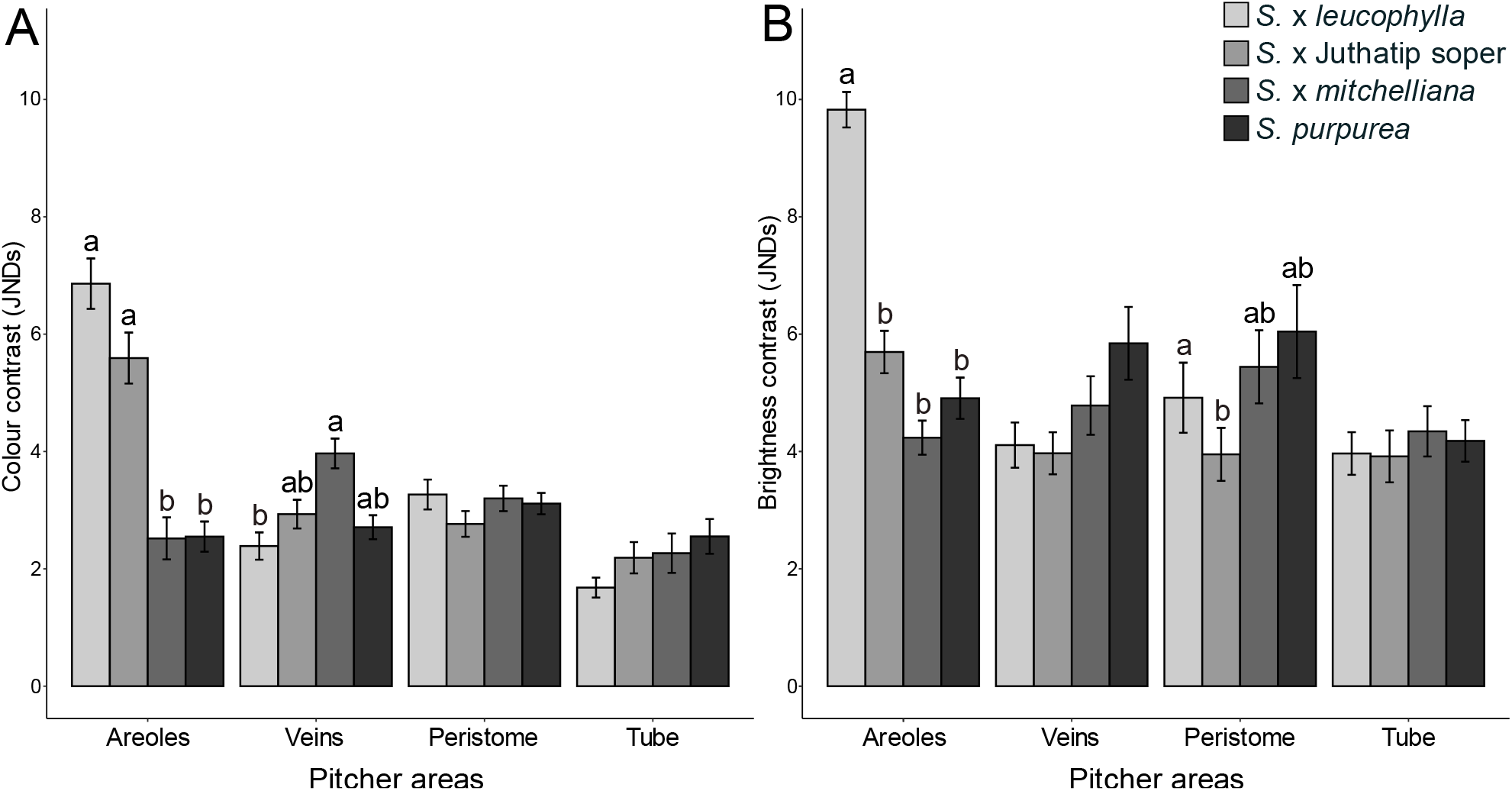
Variation in colour contrast (A) and brightness contrast (B) displayed by the different pitcher areas against the green background, as perceived by flying Hymenoptera. Mean values are presented with their associated standard errors. Contrasts are expressed in Just Noticeable Differences (JNDs). Different letters above bars show statistically significant differences in means between plant taxa for the contrasts of each specific area, no letter meaning that differences are not statistically different (p>0.05).

The change in colour and brightness contrasts with pitcher stage was not similar in all taxa (Table 1). However, areoles were the area most contrasting in colour at stage 1 (~5 days after pitcher opening) and at stage 2 (~15 days after pitcher opening) while the peristome was the area most contrasting in colour at stage 3 (~40 days after pitcher opening) and at stage 4 (~60 days after pitcher opening) (Supporting Information Fig. S4A, Table S1). Regarding brightness, areoles were the most contrasting pitcher areas against background at stage 1, 3 and 4 but only the second most contrasting areas at stage 2, veins being the most contrasting areas for this stage (Fig. S4B, Table S2). Season also directly affected the colour but not brightness contrasts with background, the colour contrasts being more pronounced in autumn than in summer (Table 1, Table S1).

### CONTRASTS AMONG PITCHER AREAS

The visual contrasts – colour contrast and brightness contrast –between pitcher areas seen by flying Hymenoptera varied with taxon and stage (Table 2). More specifically, as seen by flying Hymenoptera, areoles produced a higher colour contrast with any other area in *S.* × *leucophylla* than in the 3 other taxa (Fig. 3A). These contrasts were also generally higher for *S.* × Juthatip soper than for *S.* × *mitchelliana* and *S. purpurea* but this was not the case for the areoles-veins contrast, which was not significantly different between *S.* × Juthatip soper and *S.* × *mitchelliana* (Fig. 3A). Areoles displayed a generally higher brightness contrast against any other area in *S.* × *leucophylla* than in the 3 other taxa but this was not the case for the areoles-veins contrast, which was not significantly different between *S.* × *leucophylla* and *S.* × *mitchelliana* and for the areoles-tube contrast, which was not significantly different between *S.* × *leucophylla* and *S.* × Juthatip soper (Fig. 3B). Overall, colour contrasts values between any two pitcher areas were always higher when they involved the areoles than when they involved any other areas for these two taxa (Supporting Information Table S3, Table S4). Yet, the pair of areas that displayed the strongest contrasts was not always the same from one taxon to another (Table S3, Table S4). For the two long-leaved taxa, *S.* × *leucophylla* and *S.* × Juthatip soper, the strongest contrast for colour or brightness was produced by areoles against peristome (Table S3, Table S4). For *S*. × *mitchelliana* the strongest colour contrast was produced by veins against tube and peristome against tube and the strongest brightness contrast was produced between any red areas (peristome or veins) against any green area (areoles or tube). For *S. purpurea,* the strongest colour or brightness contrast was produced by areoles against peristome and peristome against tube, the brightness contrast being higher between areoles and peristome than between tube and peristome.

**Table 2.** Factors of variation in colour and brightness contrasts between two any pitcher areas. The best model is presented. Both dependent variables were square-root transformed to match residual normality. “Chisq” values refer to type III Wald chi-square tests. *: p<0.05, **: p<0.01, and ***: p<0.001. Contrasts are detailed in ESM, Table S3 for colour contrast and S4 for brightness contrast. “ : ” refers to the interaction between variables.

| Explanatory variables | Colour contrast |  |  | Brightness contrast |  |  |
| --- | --- | --- | --- | --- | --- | --- |
|  | Chisq | Df | p-value | Chisq | Df | p-value |
| Intercept | 132.85 | 1 | <0.001 *** | 55.23 | 1 | <0.001 *** |
| Plant taxon | 13.14 | 3 | 0.0043 ** | 2.55 | 3 | 0.4662 |
| Pair of areas | 347.46 | 5 | <0.001 *** | 259.38 | 5 | <0.001 *** |
| Pitcher stage | 2.12 | 3 | 0.5481 | 4.09 | 3 | 0.2520 |
| Season | 56.42 | 1 | <0.001 *** | 7.03 | 1 | 0.0080 ** |
| Plant taxon: Pair of areas | 553.75 | 15 | <0.001 *** | 174.22 | 15 | <0.001 *** |
| Plant taxon: Pitcher stage | 30.21 | 9 | <0.001 *** | 6.45 | 9 | 0.6941 |
| Pair of areas: Pitcher stage | 241.65 | 15 | <0.001 *** | 199.66 | 15 | <0.001 *** |
| Plant taxon: Pair of areas : Pitcher stage | 1114.93 | 45 | <0.001 *** | 674.58 | 45 | <0.001 *** |

**Figure 3.**
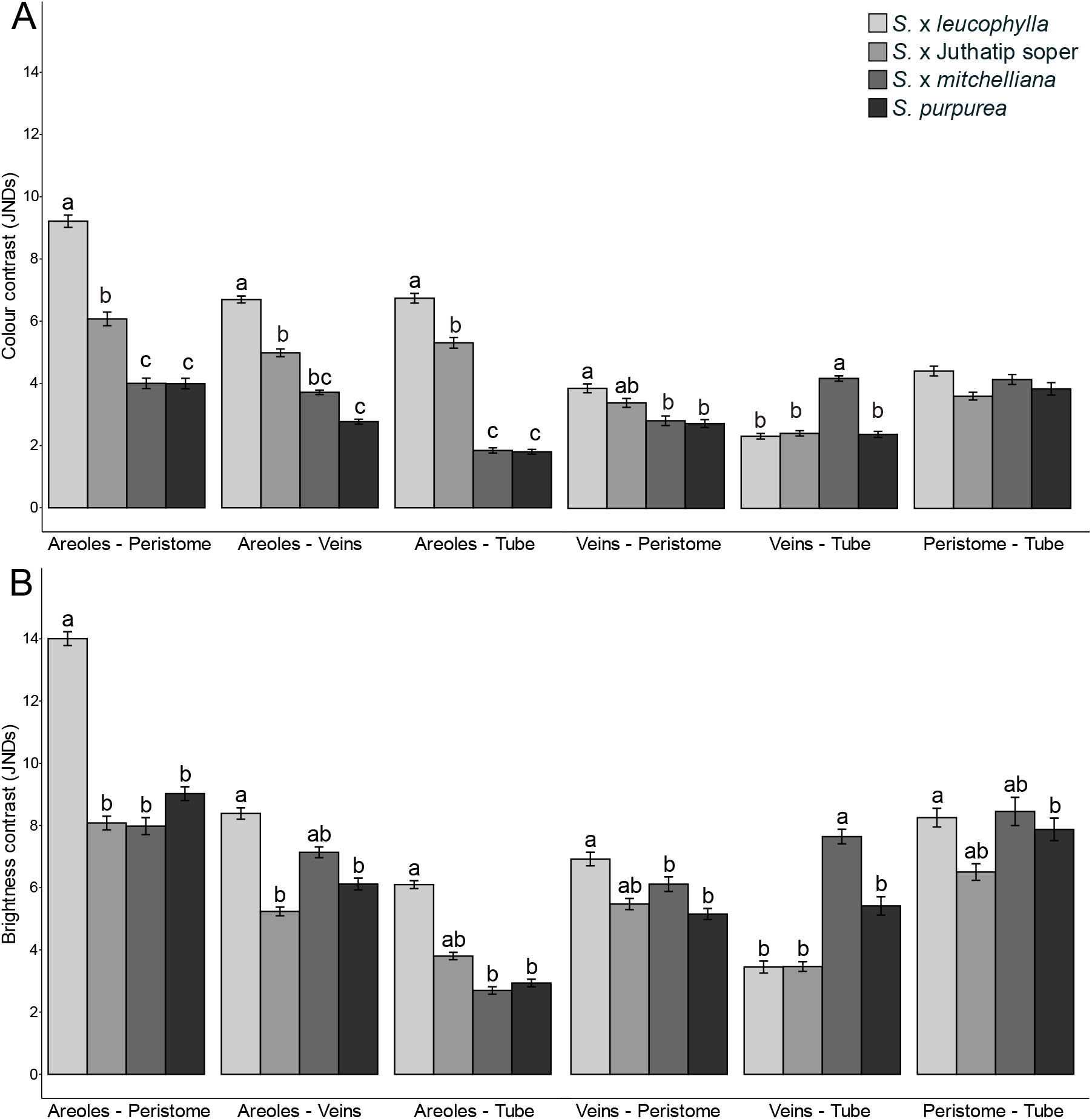
Variation among plant taxa in colour contrast (A) and brightness contrast (B) between any two pitcher areas (areoles, peristome, veins, or tube), as perceived by flying Hymenoptera. Mean values are presented with their associated standard errors. Contrasts are expressed in Just Noticeable Differences (JNDs). Different letters above bars show statistically significant differences in means between plant taxa for the contrasts of each specific area, no letter meaning that differences are not statistically different (p>0.05).

The differences in contrast produced by each pair of areas depended on the pitcher stage (Table 2, Supporting Information Fig. S5) but areoles and peristome always produced the highest contrast at stages 2 and 3 regarding colour and at stages 1, 2 and 3 regarding brightness (Table S3, S4). At stage 4, the most contrasting pairs of areas regarding colour and brightness contrasts involved the peristome against both areoles and tube (Table S3, S4). Overall, pitcher stages did not show many differences (Table 2), but differences occurred according to the pair of areas considered (Table 2, Fig. S5). Pitchers of stage 1 showed a weaker brightness contrast than pitchers of other stages, and the contrast displayed by areoles or tube against peristome or veins was always the lowest for that stage (Table S4). Regarding contrasts produced by areoles against peristome, stage 2 was the highest for colour but stage 3 was the highest for brightness. Moreover, stage 2 displayed the highest brightness contrast between areoles and veins, while the most brightness contrasting pairs of areas of stages 3 and 4 were peristome and tube. Finally, colour and brightness contrasts between areas varied with season and were generally more pronounced in autumn than in summer (Table 2, S3, and S4).

### PREY CAPTURE AND PITCHER FEATURE

Prey spectra varied with plant taxon (Supporting Information Fig. S6). The prey trapped by pitchers were mostly composed of flying Hymenoptera and Diptera in *S.* × *leucophylla* (35 ± 8 %, 23 ± 10%); flying Hymenoptera, ants (crawling Hymenoptera) in *S.* × Juthatip soper (39 ± 7%, 39 ± 8%); ants and Diptera in *S.* × *mitchelliana* (32 ± 13 % and 21 ± 8%) and ants in *S. purpurea* (36 ± 16 %). Flying Hymenoptera trapped were mostly solitary bees (93 ± 6%) with a clear dominance of sweet bees (Halictidae).

The total number of prey individuals and crawling Hymenoptera caught in pitchers depended on plant taxon and pitcher stage. Likewise, pitcher morphology affected the total number of prey individuals (Supporting Information Table S5). Pitchers of *S.* × *leucophylla* and *S.* × Juthatip soper trapped the highest total number of prey individuals while pitchers of *S.* × *mitchelliana* and *S. purpurea* trapped the lowest number of prey individuals (Fig. 4, Table 3). As regards crawling Hymenoptera, *S.* × Juthatip soper pitchers trapped more individuals than other taxa (Fig. 4, Table 3). Pitchers of stage 1 trapped the lowest number of prey individuals and crawling Hymenoptera individuals while pitchers of stage 3 trapped the highest total number of prey individuals followed by pitchers of stage 2 (Table 3). Stage 4 trapped more crawling Hymenoptera individuals than other stages (Table 3). Regarding morphology, the total number of prey individuals increased with aperture width and pitcher length (Table 3).

**Figure 4.**
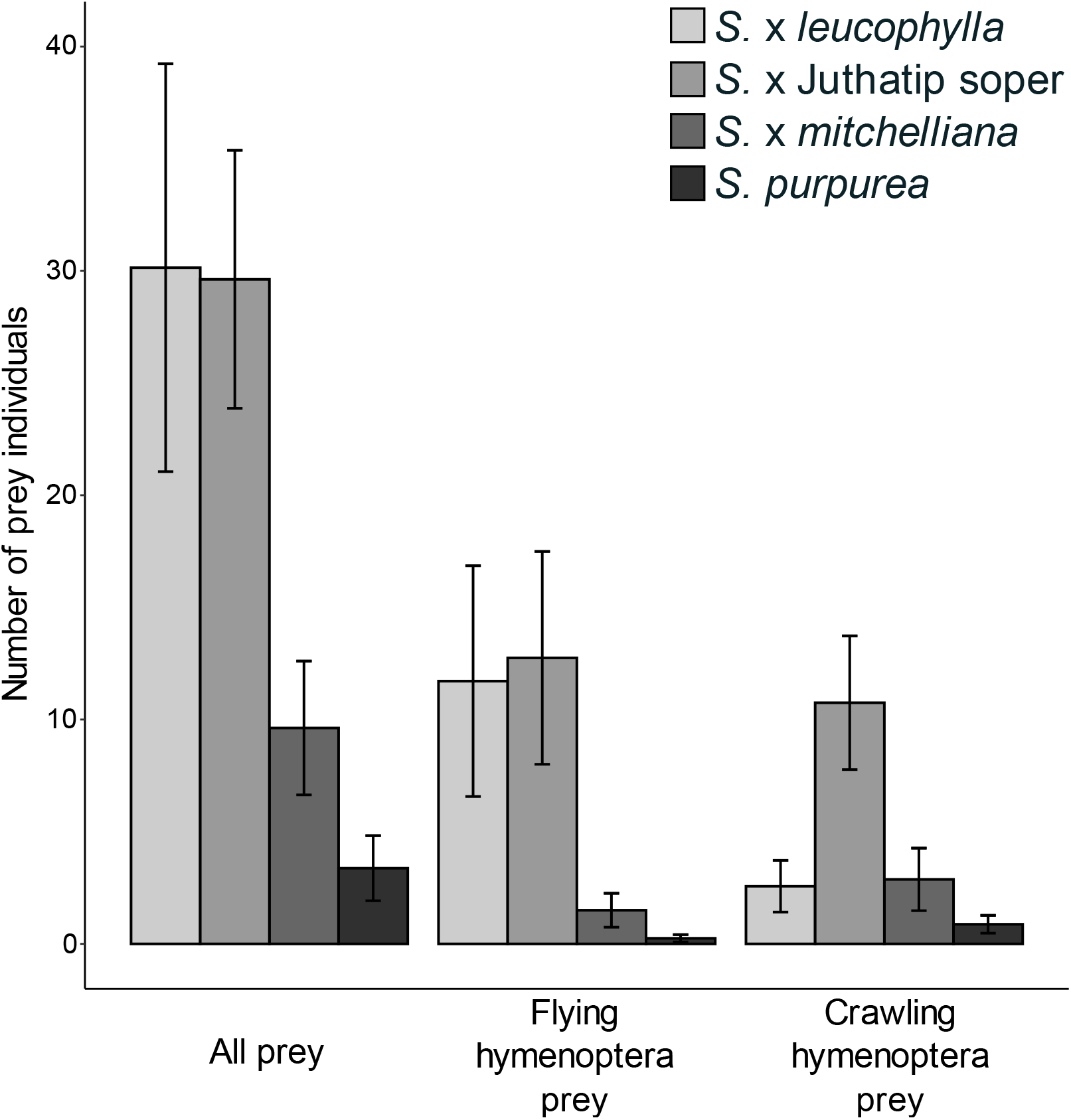
Variation among plant taxa in the number of prey individuals captured. Mean values are presented with their associated standard errors. We consider all prey, flying Hymenoptera prey and crawling Hymenoptera prey (ants).

**Table 3.**
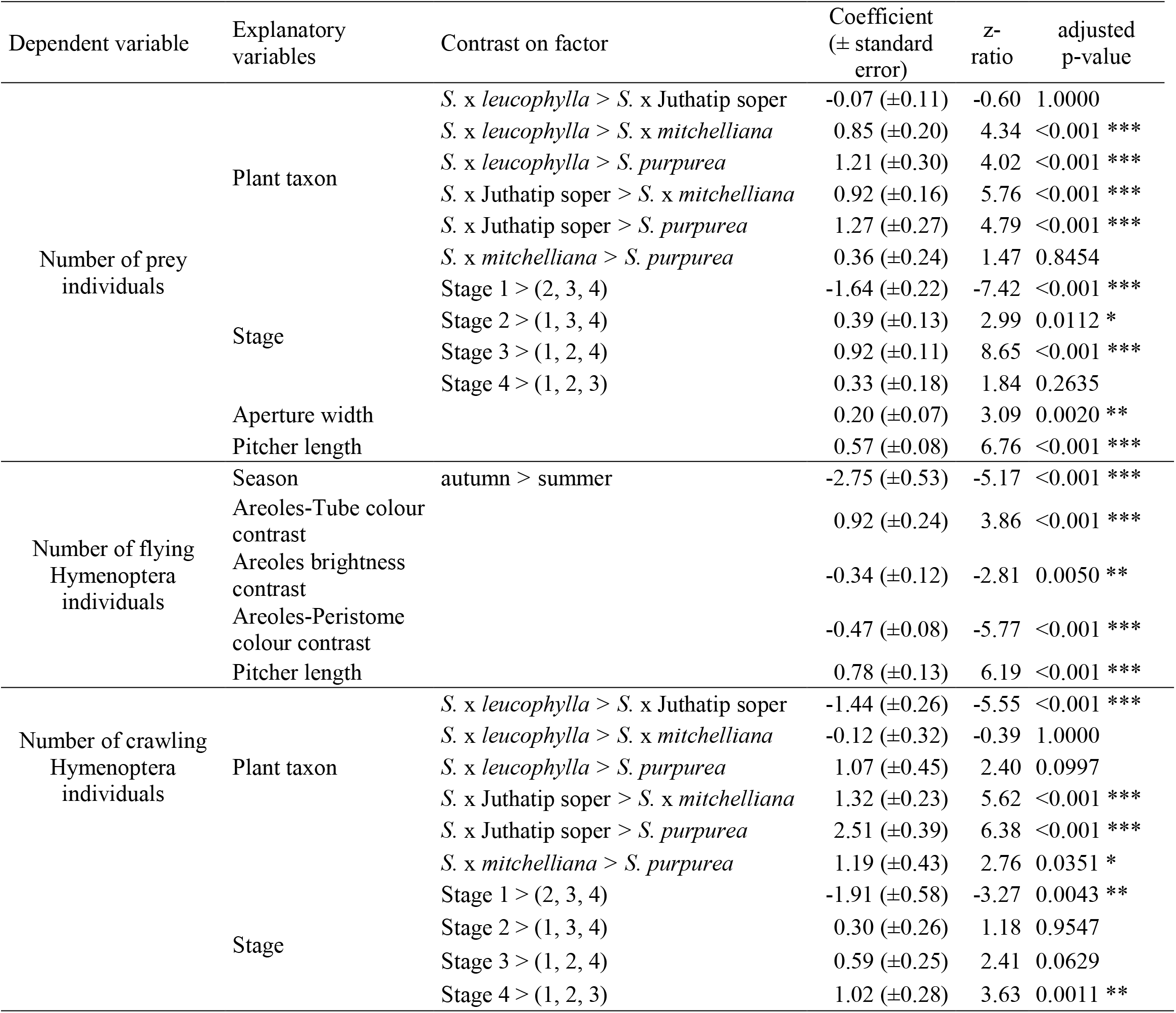
Effects of covariates included in the best linear models presented in Table S5 to explain the variations in the number of prey individuals, the number of flying Hymenoptera individuals and the number of crawling Hymenoptera individuals caught in the pitchers. We used a GLM with a Poisson distribution. P-values were adjusted for multiple comparisons with the Bonferroni correction, *: p<0.05, **: p<0.01, and ***: p<0.001. For qualitative variables, we tested the contrast between two levels of factor or between 2 gathered levels of factor. For instance, we tested whether the number of prey individuals in S. × *leucophylla* was higher than the number of prey individuals in S. × Juthatip soper and it was significantly different (p-value<0.001). For quantitative variables, which were centred and scaled before analysis to assess relative effect sizes, we showed the estimated regression coefficients of the model.

The number of flying Hymenoptera individuals caught in pitchers depended on season, visual signals and pitcher morphology (Table S5). Regarding morphology, the number of flying Hymenoptera individuals increased with pitcher length (Table 3). Pitchers also trapped less flying Hymenoptera individuals in autumn than in summer (Table 3). In terms of visual signals, the colour contrast between areoles and tube had a positive effect on the number of flying hymenopterans caught in pitchers while the colour contrast between areoles and peristome and the areoles brightness contrast with background had a negative one (Table 3).

## DISCUSSION

### PITCHER COLOURATION AND ATTRACTION OF FLYING HYMENOPTERA

Beyond the broad range of colouration patterns within pitchers and between plant taxa, our findings support the hypothesis that the clear areoles – a characteristic of the two long-leaved taxa related to *S. leucophylla, S.* × *leucophylla* and *S.* × Juthatip soper – may potentially help *Sarracenia* pitcher plants attracting flying Hymenoptera. (i) Clear areoles contrasted strongly both in colour and brightness with the other parts of the pitcher and were the areas that contrasted the most against the green background, making them the most conspicuous area in the eyes of flying Hymenoptera. Flying Hymenoptera, such as honeybees and bumblebees, use both colour and brightness signals for flower detection (Giurfa *et al*., 1996; Giurfa & Vorobyev, 1998; Dyer *et al*., 2008). Brightness contrast is used for small or distant objects (subtending a small visual angle in the eye) while colour contrast is used for large or close objects (subtending a large visual angle in the eye) (Giurfa *et al*., 1996; Dyer *et al*., 2008). Hence, bright areoles may play a role as a long-distance attractant for flying Hymenoptera, because they make the upper part with the aperture of the trap more visible compared to the rest of the pitcher. (ii) Furthermore, the two taxa that sported clear areoles also trapped more prey in total and trapped a majority of flying hymenoptera compared to the two smallest-leafed taxa without clear areoles, *S.* × *mitchelliana* and *S. purpurea,* that trapped a majority of ants. (iii) In addition, the colour contrast between areoles and tube explained an important part of the variation in the number of flying Hymenoptera trapped, the number of captures increasing with this contrast. On the other hand, we observed a negative effect of areoles brightness contrast and areoles-peristome colour contrast. Such a seemingly contradictory effect can be explained by the fact that colour and brightness contrasts are not particularly correlated. The pitchers that contrasted strongly in colour between areoles and tube are not the ones that contrasted the most in brightness for their areoles or in colour between areoles and peristome. In any case, the effect of the areoles-tube contrast outweighed the other two negative effects combined and is thus the most reliable effect. Considering the spectrum of attracted insects instead of the spectrum of trapped insects would have helped to better assess the relative contribution of the interacting effects, since only a few insects attracted are bound to be trapped (Joel, 1988; Newell & Nastase, 1998). However, attraction is a prerequisite to capture and our results support the hypothesis that areoles help attraction by contrasting highly in colour against the tube, a result coherent with that found by Schaefer & Ruxton (2014). *Sarracenia* form dense groups of plants (Gibson, 1983b) and areoles are often seen against a background of other pitcher tubes. Areoles would thus lure flying insects to the deadly traps, as also shown by McGregor *et al.* (2016) and Moran *et al.* (2012a). Although we cannot exclude a role of areoles as light traps, in this study we suggest that areole high visual contrasts may help long-range attraction, a potential role, which has rarely been considered previously. Most of the insects caught in the traps of carnivorous plants are hymenopterans and dipterans (Juniper *et al*., 1989; Ellison & Gotelli, 2009). While Hymenoptera cannot detect long wavelengths (red) and use brightness to detect red colours since the latter poorly reflect at shorter wavelengths, appearing them dark (Martínez-Harms *et al*., 2010), many flies have a tetrachromatic or even pentachromatic vision (Kooi *et al*., 2021) and they are able to see red wavelengths both in colour and brightness. While most studies regarding carnivorous pitcher plants have overlooked how pitcher colours are perceived by potential prey (Newell & Nastase, 1998; Green & Horner, 2007; Schaefer & Ruxton, 2008; Bennett & Ellison, 2009), our results underline the importance of red colouration, already considered in these previous studies, in enhancing the visual contrast even seen by flying Hymenoptera. Indeed, for all plant taxa, whether for colour or brightness, the peristome produced one of the most important contrast either against the tube or against the areoles. The red peristome, which borders the mouth of the carnivorous pitcher, is a strategic area (Moran, 1996; Bohn & Federle, 2004; Kurup *et al*., 2013) and any contrasting patterns that concerns this area is of high advantage in terms of plant fitness, as it is likely to favour insect guidance to the deadly trap. In the two short-leaf plants *S.* × *mitchelliana* and *S. purpurea*, the red peristome provides similar brightness and colour contrast with green areoles or with green tube. In the two long-leaved plants, *S.* × *leucophylla and S.* × Juthatip soper, the red peristome contrasts more in brightness and in colour with white-pinkish areoles than with green tube, which increases pitcher visibility. Areoles when distinct in colouration from the pitcher tube, add to pitcher visibility and are associated to higher prey capture in the two long-leaved plants compared to the two short-leaf plants.

In our study, all measured pitcher areas show very poor UV-reflection in all taxa (Fig. S2), thus showing virtually no contrast in the UV range. Whether a UV contrast may exist locally between highly-absorbing (like the UV-induced fluorescence documented in Sarraceniaceae by Golos (2020)) and highly-reflecting areas remains to be studied. Such highly-localised UV-contrasts may help orientation, as suggested in Kurup *et al.* (2013), but their role, if any, in attraction and capture is controversial and remains to be elucidated (Jansen, 2017).

### INFLUENCE OF OTHER PITCHER TRAITS ON PREY CAPTURE AND COMPOSITION

Besides colouration, our results show that pitcher morphology likely plays a role in prey capture. Pitchers with a longer size and a larger aperture trapped significantly more prey in total. This is consistent with the results of Cresswell (1993) and Heard (1998) on *S. purpurea* and of Green & Horner (2007) and Bhattarai & Horner (2009) on *S. alata,* who showed that larger pitchers trapped more prey. Bhattarai & Horner (2009) supposed that the positive relationship often observed between pitcher size and prey capture was due to a greater quantity of attractants produced by larger pitchers and not merely a larger area of capture. Indeed, the capture area as defined by Bhattarai & Horner (2009) did not better explain variation in our models than aperture width. Larger pitcher apertures also mean larger surface areas of lid, which was the most nectar-rewarding area of the pitchers in the studied plants (TH, JP and LG personal observations). Nevertheless, previous studies were limited to a single taxon. By considering different taxa, we showed that the effect of pitcher length was still more important than the effect of aperture width. The hypothesis of Bhattarai & Horner (2009) is therefore plausible on an intraspecific scale, but on an interspecific scale, the shape of the trap must also play an important role in prey capture.

Pitcher length was also positively linked to the number of flying Hymenoptera individuals, which suggests that taller pitchers may be more efficient at retaining flying Hymenoptera and/or may be more attractive, for instance by their colour signals as shown above. The conspicuousness of taller pitchers can arise from their size and/or their colouration: here clear areoles were only present in long-leaved taxa. Both aspects – size and colouration – can be under selection. Such multi-component signals are frequent in the contexts where plant appearance is under selection. For instance, in a prey-pollinator conflict regarding sundew carnivorous plants, El-Sayed *et al.* (2016) showed that in addition to different colouration between flower and trap, spatial separation is necessary to attract pollinator to the flowers rather than to the traps. Likewise, in *Nepenthes* liana, which explore both terrestrial and aerial strata, two pitcher types “lower” and “upper” are produced, each one targeting a different group of prey, the “upper” type trapping more flying insects than the “lower” type (Moran, 1996; Di Giusto *et al*., 2008; Gaume *et al*., 2016). Similarly, in long-leaved *Sarracenia* plants, pitchers are likely more accessible to flying insects. In *Nepenthes rafflesiana* (Jack.), this is also explained by the emission by “upper” pitchers of volatile compounds, which attract a diversity of flying insects (Di Giusto *et al*., 2010). Whether *Sarracenia* taxa differ in their emission of odour attractive to prey remains to be studied.

### VARIATION IN VISUAL SIGNALS AND PREY CAPTURE OVER PITCHER LIFE SPAN

Our study also show that both colour contrasting patterns and prey capture changed during pitcher life span. We observed an increase in capture at stage 2 and a peak at stage 3 for the number of prey individuals. Although our sampling is weak for stages 1 and 4, mixed models control for unbalanced observations and show that capture increases over the first part of pitcher life (stages 1 to 3) and later (stage 4) decreases, which implies that when aging, pitchers allocated their resources more to the digestive process and less to attraction and capture. Heard (1998) observed a similar peak in prey capture in 12-33-day-old pitchers (corresponding to stages 2 and 3 in our study) in *S. purpurea*. Several studies on pitcher plants have reported that the accumulation of ammonia released by dead prey when prey capture is too important leads to digestion disruption and pitcher putrefaction (Clarke & Kitching, 1995). Increase in ammonia with prey capture has been recorded for *Sarracenia* (Wakefield *et al.*, 2005), suggesting prey capture should not exceed a threshold. Enzyme activity increases with pitcher age in *Sarracenia* (Luciano & Newell (2017); hydrolase secretion increases after pitcher opening for *S. purpurea* but stops in absence of prey according to Gallie & Chang (1997). Hence, younger stages are likely specialized in attracting and capturing prey, a hypothesis that is supported by the variation in colouration that we observed. (i) Areoles are the most contrasting area for colour in younger stages (1 and 2) while peristome is the most contrasting area for colour in older stages, (ii) areoles-peristome contrast is stronger at stage 2 for colour and stronger at stage 3 for brightness and areoles-veins brightness contrast is stronger at stage 2 (iii) areoles against peristome produced the highest contrast on pitchers for colour at stages 2, 3 and for brightness at stages 1, 2, 3, while at stage 4 areoles do not contrast more than tube against peristome for colour and brightness.

Variation in colouration with pitcher age has been mentioned by Bhattarai & Horner (2009) and has been already shown in *S. alata* by Horner *et al.* (2012). Leaf colour plasticity is already known in pitcher plants: nitrogen supply or food intake has been shown to affect foliar reflectance pattern (Moran & Moran, 1998; Yoon *et al*., 2019) or chlorophyll activity (Farnsworth & Ellison, 2008). Therefore, it remains an open question whether the change in colouration of pitchers as they age is a consequence of prey capture. We also observed a seasonal variation in pitcher colouration with more pronounced contrasts in autumn, although our design allows us to this only for stage 2 pitchers. In parallel, pitchers seem to capture more flying hymenoptera individuals in summer than in autumn. Bees representing the majority of flying hymenoptera trapped by our pitchers. Like many other Hymenoptera, bees are thermophile species and are more active in summer than in autumn. Summer good weather conditions maximize time spent foraging (Vicens & Bosch, 2000). As season did seem to have neither an effect on the total number of prey individuals nor on the total number of ants, an increase in the capture of other non-Hymenoptera prey is likely to have compensated for the observed decrease in the capture of flying Hymenoptera (see Supporting Information Fig. S7). An increase in the capture of non-Hymenoptera flower visitors for instance may result from a reduced availability of flowers redirecting them toward nectar-producing pitchers - or from pitcher attraction towards different groups of insects with various visual ability - or capture efficiency, a question that deserves proper study. Yet, as we focused on the relation between areoles and flying Hymenoptera and used a specific vision model for that purpose, it is difficult to consider the effect of visual signals on the whole prey spectrum. For instance, ant workers, which represent a large part of the prey spectra of *Sarracenia* are often dichromate (Aksoy & Camlitepe, 2018). It would therefore be interesting to model their vision to better understand pitcher colours as perceived by a completely different type of prey, especially since their capture variation among stage and taxon differed a lot from that of flying Hymenoptera. Moreover, carnivorous plants have been suggested to lure prey by other attractants such as nectar (Bennett & Ellison, 2009) and odours (Jürgens *et al*., 2009). For example, Bhattarai & Horner (2009) showed that volatiles from *Sarracenia* pitchers attract a majority of flying insects but few crawling ones, suggesting an important role of odours in the attraction of flying prey.

Studies of these other attractants could help us to better understand the variation in prey capture and complete our knowledge on the attraction strategy of *Sarracenia* pitcher plants.

## Supporting information

Dupont_supporting_information

## ACKNOWLEDGMENTS

We thank Jean-Jacques Labat from the “Nature et Paysages” nursery for providing us with *Sarracenia* pitcher plants, Stéphane Fourtier, Merlin Ramel, and the technical staff from the AMAP lab for setting up, and growing plants.

## SUPPORTING INFORMATION

Dupont_supporting_information.pdf

## REFERENCES

Adlassnig W, Lendl T, Peroutka M, Lang I. 2010. Deadly glue—adhesive traps of carnivorous plants. In: Byern J and Grunwald I, eds. Biological Adhesive Systems From Nature To Technical And Medical Application. Vienna, Austria: Springer, 15–28.

Aguiar JMRBV, Maciel AA, Santana PC, Telles FJ, Bergamo PJ, Oliveira PE, Brito VLG. 2020. Intrafloral color modularity in a bee-pollinated orchid. Frontiers in plant science 11.

Aksoy V, Camlitepe Y. 2018. Spectral sensitivities of ants–a review. Animal Biology 68: 55–73.

Bauer U, Scharmann M, Skepper J, Federle W. 2013. ‘Insect aquaplaning’ on a superhydrophilic hairy surface: how *Heliamphora nutans* Benth. pitcher plants capture prey. Proceedings of the Royal Society B: Biological Sciences 280: 20122569.

Bennett KF, Ellison AM. 2009. Nectar, not colour, may lure insects to their death. Biology Letters 5: 469–472.

Bhattarai GP, Horner JD. 2009. The importance of pitcher size in prey capture in the carnivorous plant, *Sarracenia alata* Wood (Sarraceniaceae). The American Midland Naturalist 163: 264–272.

Bohn HF, Federle W. 2004. Insect aquaplaning: Nepenthes pitcher plants capture prey with the peristome, a fully wettable water-lubricated anisotropic surface. Proceedings of the National Academy of Sciences 101: 14138–14143.

Bonhomme V, Pelloux-Prayer H, Jousselin E, Forterre Y, Labat JJ, Gaume L. 2011. Slippery or sticky? Functional diversity in the trapping strategy of *Nepenthes* carnivorous plants. New Phytologist 191: 545–554.

Briscoe AD, Chittka L. 2001. The evolution of color vision in insects. Annual review of entomology 46: 471–510.

Chittka L, Raine NE. 2006. Recognition of flowers by pollinators. Current opinion in plant biology 9: 428–435.

Clarke CM, Kitching RL. 1995. Swimming ants and pitcher plants: a unique ant-plant interaction from Borneo. Journal of Tropical Ecology: 589–602.

Cresswell JE. 1991. Capture rates and composition of insect prey of the pitcher plant *Sarracenia purpurea*. The American Midland Naturalist 125: 1–9.

Cresswell JE. 1993. The morphological correlates of prey capture and resource parasitism in pitchers of the carnivorous plant *Sarracenia purpurea*. The American Midland Naturalist 129: 35–41.

Dafni A. 1984. Mimicry and deception in pollination. Annual review of ecology and systematics 15: 259–278.

De Ibarra HN, Vorobyev M, Menzel R. 2014. Mechanisms, functions and ecology of colour vision in the honeybee. Journal of Comparative Physiology A 200: 411–433.

Di Giusto B, Bessière JM, Guéroult M, Lim LBL, Marshall DJ, Hossaert-McKey M, Gaume L. 2010. Flower-scent mimicry masks a deadly trap in the carnivorous plant *Nepenthes rafflesiana*. Journal of Ecology 98: 845–856.

Di Giusto B, Grosbois V, Fargeas E, Marshall DJ, Gaume L. 2008. Contribution of pitcher fragrance and fluid viscosity to high prey diversity in a *Nepenthes* carnivorous plant from Borneo. Journal of Biosciences 33: 121–136.

Dobson HEM. 1994. Floral volatiles in insect biology. Insect-plant interactions 5: 47–81.

Dress W, Newell S, Nastase A, Ford J. 1997. Analysis of amino acids in nectar from pitchers of *Sarracenia purpurea* (Sarraceniaceae). American Journal of Botany 84: 1701.

Dyer AG, Spaethe J, Prack S. 2008. Comparative psychophysics of bumblebee and honeybee colour discrimination and object detection. Journal of Comparative Physiology A 194: 617.

El-Sayed AM, Byers JA, Suckling DM. 2016. Pollinator-prey conflicts in carnivorous plants: When flower and trap properties mean life or death. Scientific reports 6: 1–11.

Ellison AM, Adamec L. 2018. Carnivorous plants: physiology, ecology, and evolution. Oxford University Press.

Ellison AM, Gotelli NJ. 2009. Energetics and the evolution of carnivorous plants--Darwin’s ‘most wonderful plants in the world’. J. Exp. Bot. 60: 19–42.

Farnsworth EJ, Ellison AM. 2008. Prey availability directly affects physiology, growth, nutrient allocation and scaling relationships among leaf traits in 10 carnivorous plant species. Journal of Ecology 96: 213–221.

Forterre Y, Skotheim JM, Dumais J, Mahadevan L. 2005. How the Venus flytrap snaps. Nature 433: 421–425.

Gallie DR, Chang SC. 1997. Signal transduction in the carnivorous plant *Sarracenia purpurea*. Regulation of secretory hydrolase expression during development and in response to resources. Plant Physiol. 115: 1461–1471.

Gaume L, Bazile V, Huguin M, Bonhomme V. 2016. Different pitcher shapes and trapping syndromes explain resource partitioning in *Nepenthes* species. Ecology and Evolution 6: 1378–1392.

Gaume L, Forterre Y. 2007. A viscoelastic deadly fluid in carnivorous pitcher plants. PloS one 2: e1185.

Gaume L, Perret P, Gorb E, Gorb S, Labat JJ, Rowe N. 2004. How do plant waxes cause flies to slide? Experimental tests of wax-based trapping mechanisms in three pitfall carnivorous plants. Arthropod structure & development 33: 103–111.

Gibson TC. 1983a. Competition, disturbance and the carnivorous plant community in south eastern US. Unpublished D. Phil. Thesis, University of Utah.

Gibson TC. 1983b. Competition, disturbance, and the carnivorous plant community in the southeastern United States. Unpublished Ph.D. Thesis Thesis, Univ. of Utah.

Giurfa M, Vorobyev M. 1998. The angular range of achromatic target detection by honey bees. Journal of Comparative Physiology A 183: 101–110.

Giurfa M, Vorobyev M, Kevan P, Menzel R. 1996. Detection of coloured stimuli by honeybees: minimum visual angles and receptor specific contrasts. Journal of Comparative Physiology A 178: 699–709.

Golos MR. 2020. First observations of UV-induced fluorescence in *Heliamphora* (Sarraceniaceae) and other tepui flora. Carnivorous Plant Newsletter 49.

Green ML, Horner JD. 2007. The relationship between prey capture and characteristics of the carnivorous pitcher plant, *Sarracenia alata* Wood. The American Midland Naturalist 158: 424–431.

Grimaldi D. 1999. The co-radiations of pollinating insects and angiosperms in the Cretaceous. Annals of the Missouri Botanical Garden: 373–406.

Heard SB. 1998. Capture rates of invertebrate prey by the pitcher plant, *Sarracenia purpurea* L. The American Midland Naturalist 139: 79–89.

Horner JD, Steele JC, Underwood CA, Lingamfelter D. 2012. Age-related changes in characteristics and prey capture of seasonal cohorts of *Sarracenia alata* pitchers. The American Midland Naturalist 167: 13–27.

Jaffé K, Blum MS, Fales HM, Mason RT, Cabrera A. 1995. On insect attractants from pitcher plants of the genus *Heliamphora* (Sarraceniaceae). Journal of chemical ecology 21: 379–384.

Jansen MAK. 2017. Carnivorous plants and UV-radiation: a captivating story? UV4Plants Bulletin 2017: 11–16.

Joel DM. 1988. Mimicry and mutualism in carnivorous pitcher plants (Sarraceniaceae, Nepenthaceae, Cephalotaceae, Bromeliaceae). Biological Journal of the Linnean Society 35: 185–197.

Joel DM, Juniper BE, Dafni A. 1985. Ultraviolet patterns in the traps of carnivorous plants. New Phytologist 101: 585–593.

Juniper BE, Robins RJ, Joel D. 1989. The carnivorous plants. Academic press: London, UK.

Jürgens A, El-Sayed AM, Suckling DM. 2009. Do carnivorous plants use volatiles for attracting prey insects? Functional Ecology 23: 875–887.

Kevan PG, Baker HG. 1983. Insects as flower visitors and pollinators. Annual review of entomology 28: 407–453.

Kooi CJvd, Stavenga DG, Arikawa K, Belušič G, Kelber A. 2021. Evolution of Insect Color Vision: From Spectral Sensitivity to Visual Ecology. Annual Review of Entomology 66: 435–461.

Kurup R, Johnson AJ, Sankar S, Hussain AA, Kumar CS, Sabulal B. 2013. Fluorescent prey traps in carnivorous plants. Plant Biology 15: 611–615.

Lehrer M. 1993. Parallel processing of motion, shape and colour in the visual system of the bee. Sensory systems of arthropods: 266–272.

Luciano CS, Newell SJ. 2017. Effects of prey, pitcher age, and microbes on acid phosphatase activity in fluid from pitchers of *Sarracenia purpurea (Sarraceniaceae*). PloS one 12: e0181252.

Lunau K, Maier EJ. 1995. Innate colour preferences of flower visitors. Journal of Comparative Physiology A 177: 1–19.

Maia R, Gruson H, Endler JA, White TE. 2019. pavo 2: new tools for the spectral and spatial analysis of colour in R. Methods in Ecology and Evolution 10: 1097–1107.

Martínez-Harms J, Palacios AG, Márquez N, Estay P, Arroyo MTK, Mpodozis J. 2010. Can red flowers be conspicuous to bees? *Bombus dahlbomii* and South American temperate forest flowers as a case in point. Journal of Experimental Biology 213: 564–571.

McGregor JP, Moon DC, Rossi AM. 2016. Role of areoles on prey abundance and diversity in the hooded pitcher plant *(Sarracenia minor*, Sarraceniaceae). Arthropod-Plant Interactions 10: 133–141.

McPherson S. 2006. Sarracenia pitcher plants of the Americas. McDonald & Woodward Publishing Company: Virginia, USA.

Meurgey F, Perrocheau R. 2015. Les Sarracénies, pièges pour le Frelon asiatique à pattes jaunes. Insectes 177.

Moran JA. 1996. Pitcher dimorphism, prey composition and the mechanisms of prey attraction in the pitcher plant *Nepenthes rafflesiana* in Borneo. The Journal of ecology 84: 515–525.

Moran JA, Booth WE, Charles JK. 1999. Aspects of pitcher morphology and spectral characteristics of six bornean *Nepenthes* pitcher plant species: implications for prey capture. Annals of Botany 83: 521–528.

Moran JA, Clarke C, Gowen BE. 2012a. The use of light in prey capture by the tropical pitcher plant *Nepenthes aristolochioides*. Plant signaling & behavior 7: 957–960.

Moran JA, Clarke C, Greenwood M, Chin L. 2012b. Tuning of color contrast signals to visual sensitivity maxima of tree shrews by three Bornean highland Nepenthes species. Plant signaling & behavior 7: 1267–1270.

Moran JA, Moran AJ. 1998. Foliar reflectance and vector analysis reveal nutrient stress in prey-deprived pitcher plants *(Nepenthes rafflesiana)*. International Journal of Plant Sciences 159: 996–1001.

Newell SJ, Nastase AJ. 1998. Efficiency of insect capture by *Sarracenia purpurea (Sarraceniaceae)*, the northern pitcher plant. American Journal of Botany 85: 88–91.

Pavlovič A, Masarovičová E, Hudák J. 2007. Carnivorous syndrome in Asian pitcher plants of the genus *Nepenthes*. Annals of Botany 100: 527–536.

Peitsch D, Fietz A, Hertel H, De Souza J, Ventura DF, Menzel R. 1992. The spectral input systems of hymenopteran insects and their receptor-based colour vision. Journal of Comparative Physiology A 170: 23–40.

R Core Team. 2020. R: A language and environment for statistical computing. Vienna, Austria: R Foundation for Statistical Computing. https://www.R-project.org/

Raguso RA. 2008. Wake up and smell the roses: the ecology and evolution of floral scent. Annual Review of Ecology, Evolution, and Systematics 39: 549–569.

Schaefer HM, Ruxton GD. 2008. Fatal attraction: carnivorous plants roll out the red carpet to lure insects. Biology Letters 4: 153–155.

Schaefer HM, Ruxton GD. 2009. Deception in plants: mimicry or perceptual exploitation? Trends in Ecology & Evolution 24: 676–685.

Schaefer HM, Ruxton GD. 2014. Fenestration: a window of opportunity for carnivorous plants. Biology Letters 10: 20140134.

Schiestl FP. 2010. Pollination: sexual mimicry abounds. Current Biology 20: R1020–R1022.

Schnell DE. 2002. Carnivorous plants of the United States and Canada. Timber Press Inc.: Portland, Oregon, USA.

Spaethe J, Schmidt A, Hickelsberger A, Chittka L. 2001. Adaptation, constraint, and chance in the evolution of flower color and pollinator color vision. In: Thomson J and Chittka L, eds. Cognitive ecology of pollination: animal behavior and floral evolution: Cambridge University Press., 106 – 126.

Streinzer M, Ellis T, Paulus HF, Spaethe J. 2010. Visual discrimination between two sexually deceptive *Ophrys* species by a bee pollinator. Arthropod-Plant Interactions 4: 141–148.

Streinzer M, Paulus HF, Spaethe J. 2009. Floral colour signal increases short-range detectability of a sexually deceptive orchid to its bee pollinator. Journal of Experimental Biology 212: 1365–1370.

Van der Niet T, Johnson SD. 2012. Phylogenetic evidence for pollinator-driven diversification of angiosperms. Trends in Ecology & Evolution 27: 353–361.

Vicens N, Bosch J. 2000. Weather-dependent pollinator activity in an apple orchard, with special reference to *Osmia cornuta* and *Apis mellifera (*Hymenoptera: Megachilidae and Apidae). Environmental Entomology 29: 413–420.

Vorobyev M, Osorio D. 1998. Receptor noise as a determinant of colour thresholds. Proceedings of the Royal Society of London. Series B: Biological Sciences 265: 351–358.

Wakefield AE, Gotelli NJ, Wittman SE, Ellison AM. 2005. Prey addition alters nutrient stoichiometry of the carnivorous plant *Sarracenia purpurea*. Ecology 86: 1737–1743.

Yoon JS, Riu YS, Kong SG. 2019. Regulation of Anthocyanin Biosynthesis by Light and Nitrogen in *Sarracenia purpurea*. Journal of Life Science 29: 1055–1061.

