## Supplementary material for "Variation in colour signals among *Sarracenia* pitcher plants and the potential role of areoles in the attraction of flying Hymenoptera": Dupont_supporting_information

**Figure S1.** Correlation table relative to visual areole-contrast variables, size variables (Pitcher length and Aperture Width) and number of flying Hymenoptera. The correlation coefficients between each pair of variables, estimated using the Pearson method, are presented with their associated p-values, *: p<0.05, **: p<0.01, and ***: p<0.001.

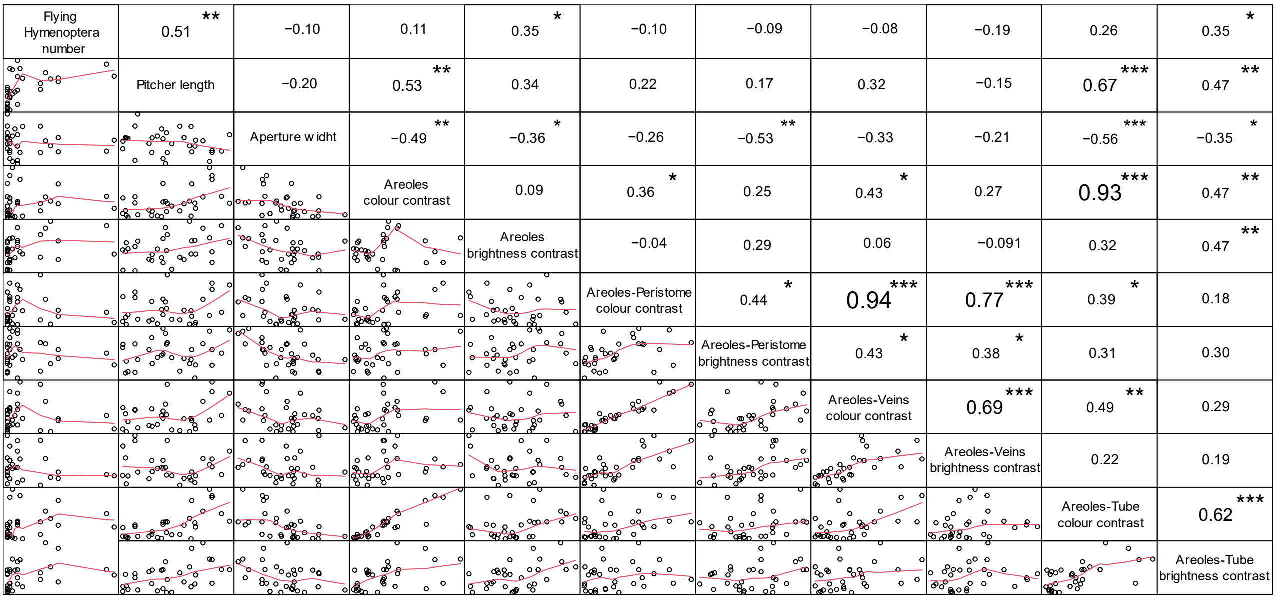

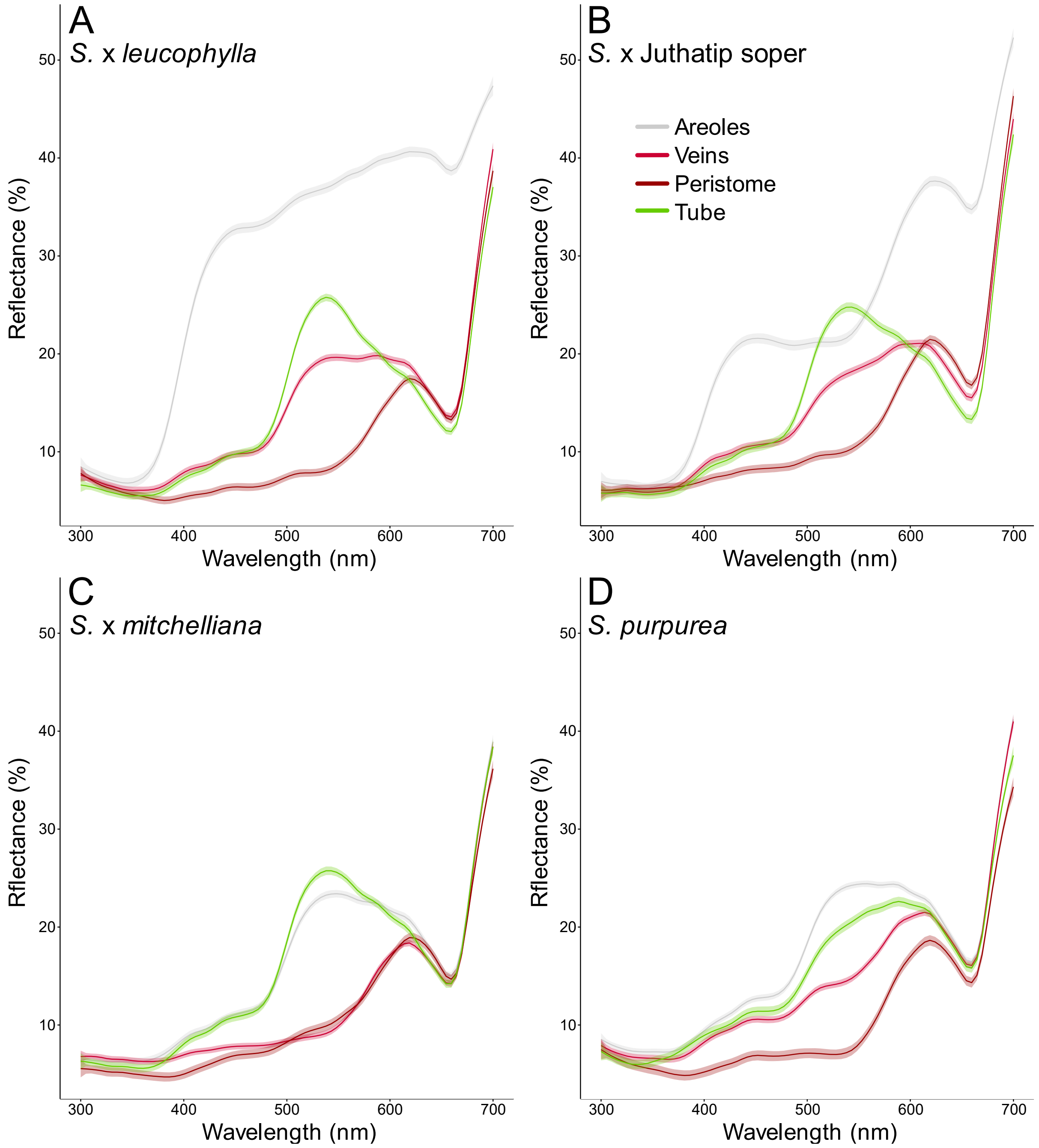


**Figure S2.** Mean reflectance spectra (± SE) of the pitcher areas in the four *Sarracenia* taxa: (A) *S.* x *leucophylla*, (B) *S.* x Juthatip soper, (C) *S.* x *mitchelliana*, (D) *S. purpurea*: areoles, veins, peristome and tube. For each taxon and pitcher area, we presented the estimated means and confident intervals of all spectral measurements obtained for all stages and seasons.

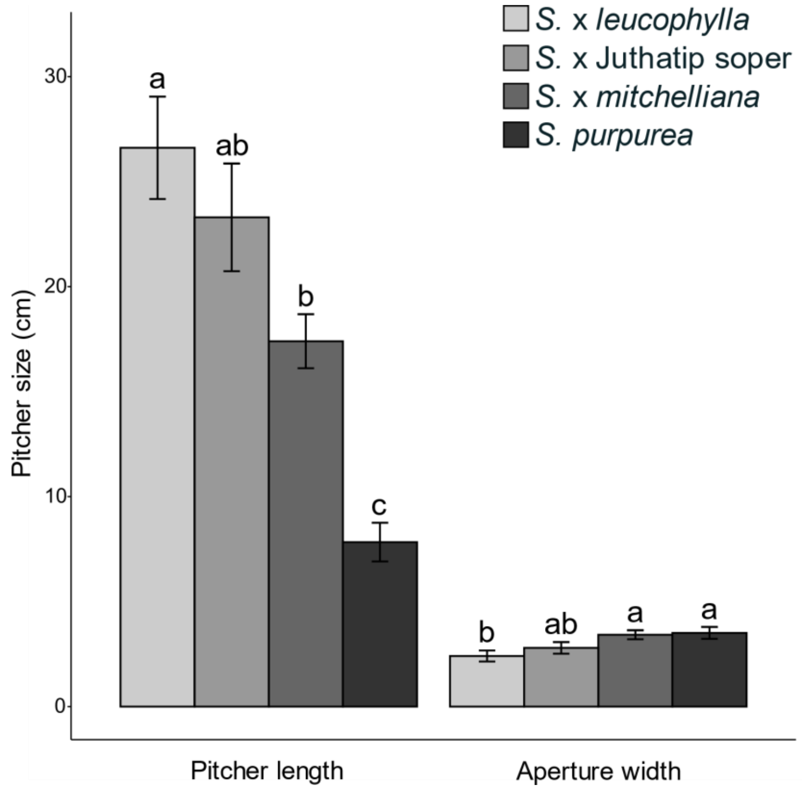


**Figure S3**. Variation among plant taxa of pitcher length and pitcher width. Mean values are presented with their associated standard errors. For each measurement, different letters above bars show statistically significant differences in means between plant taxa ($p<0.05$). For instance, pitcher length is significantly greater in S. x *leucophylla* than in S. x *mitchelliana* ($a \neq b$) and greater than in *S. purpurea* ($a \neq c$) while the length of the pitchers of S. x Juthatip soper is not different from that of S. x *leucophylla* or of S. x *mitchelliana* (ab is not different from a or from b).

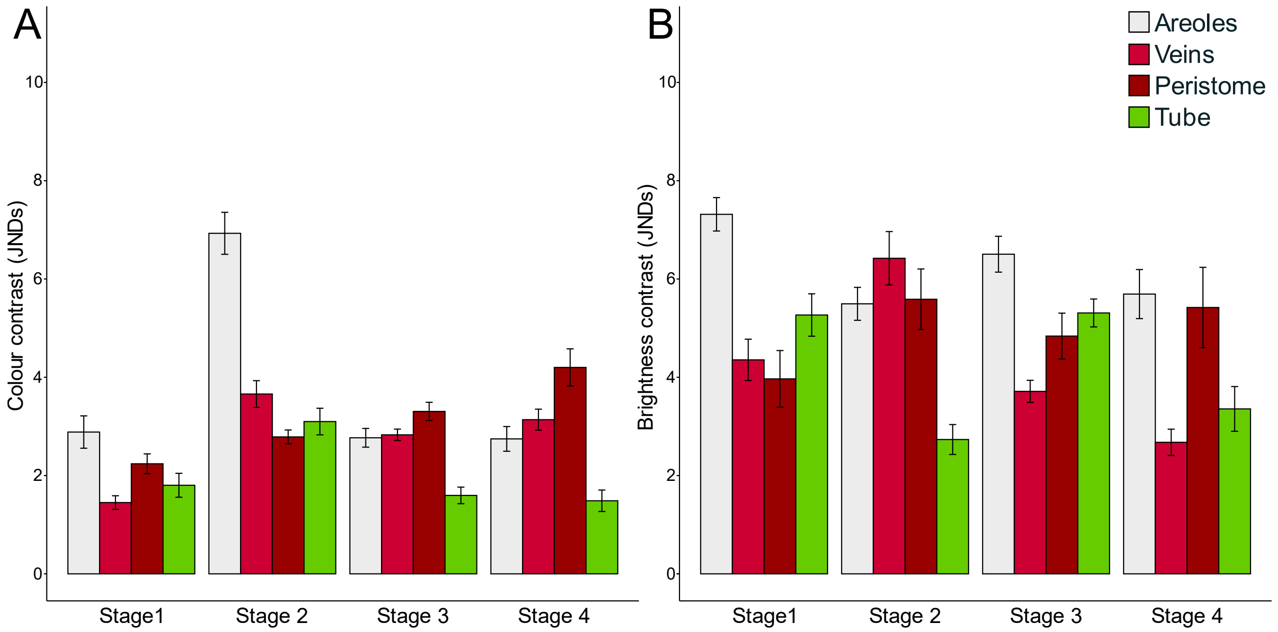


**Figure S4.** Variation among pitcher stages of colour contrast (A) and brightness contrast (B) displayed by the different pitcher areas against the green background, as perceived by flying Hymenoptera. Mean values are presented with their associated standard errors. Contrasts are expressed in Just Noticeable Differences (JNDs). Colours of the graphs do not correspond to the actual colours of the pitcher areas.

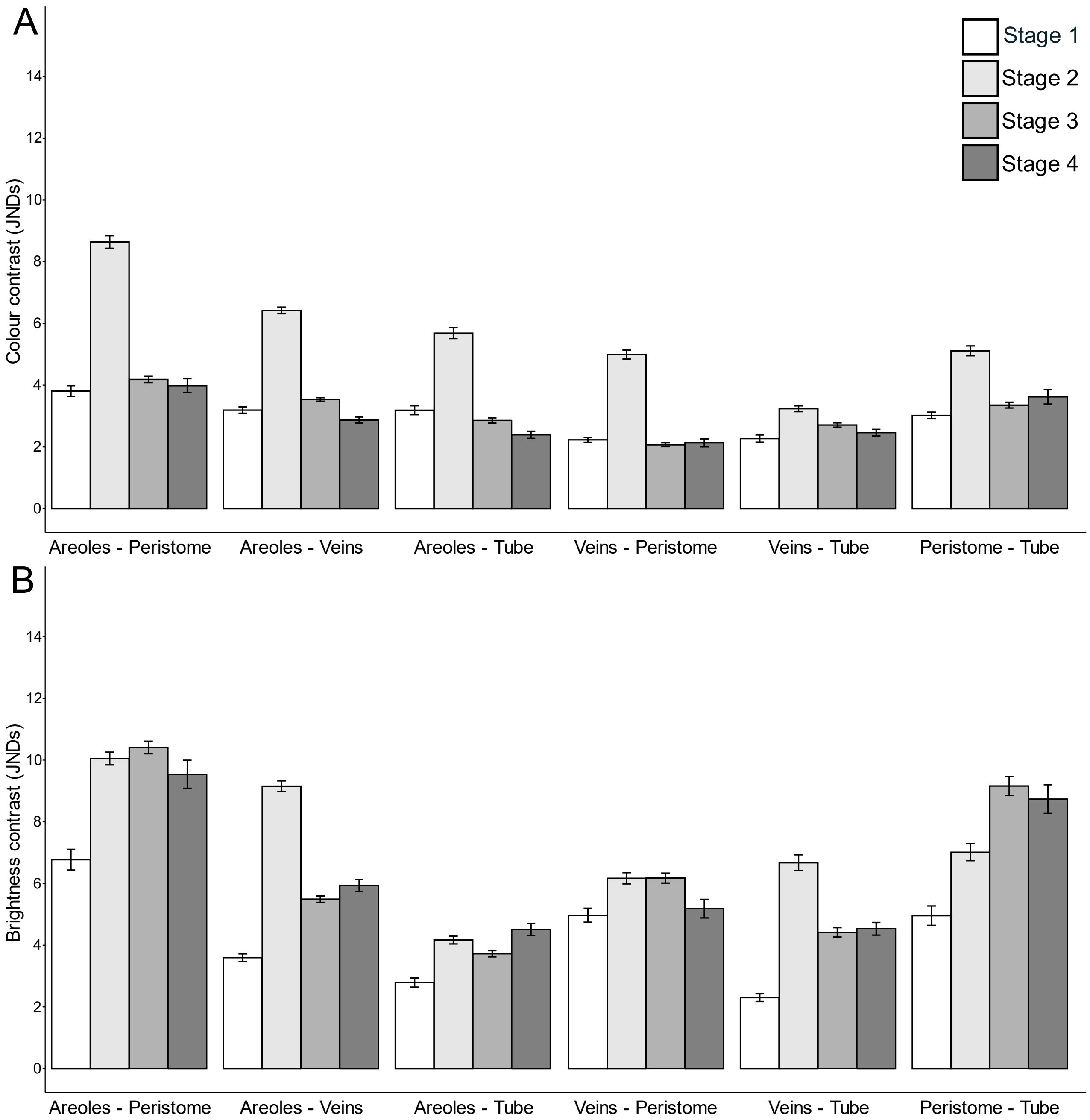


**Figure S5.** Variation among pitcher stages of colour contrast (A) and brightness contrast (B) between any two pitcher areas (areoles, peristome, veins, or tube), as perceived by flying Hymenoptera. Mean values are presented with their associated standard errors. Contrasts are expressed in Just Noticeable Differences (JNDs).

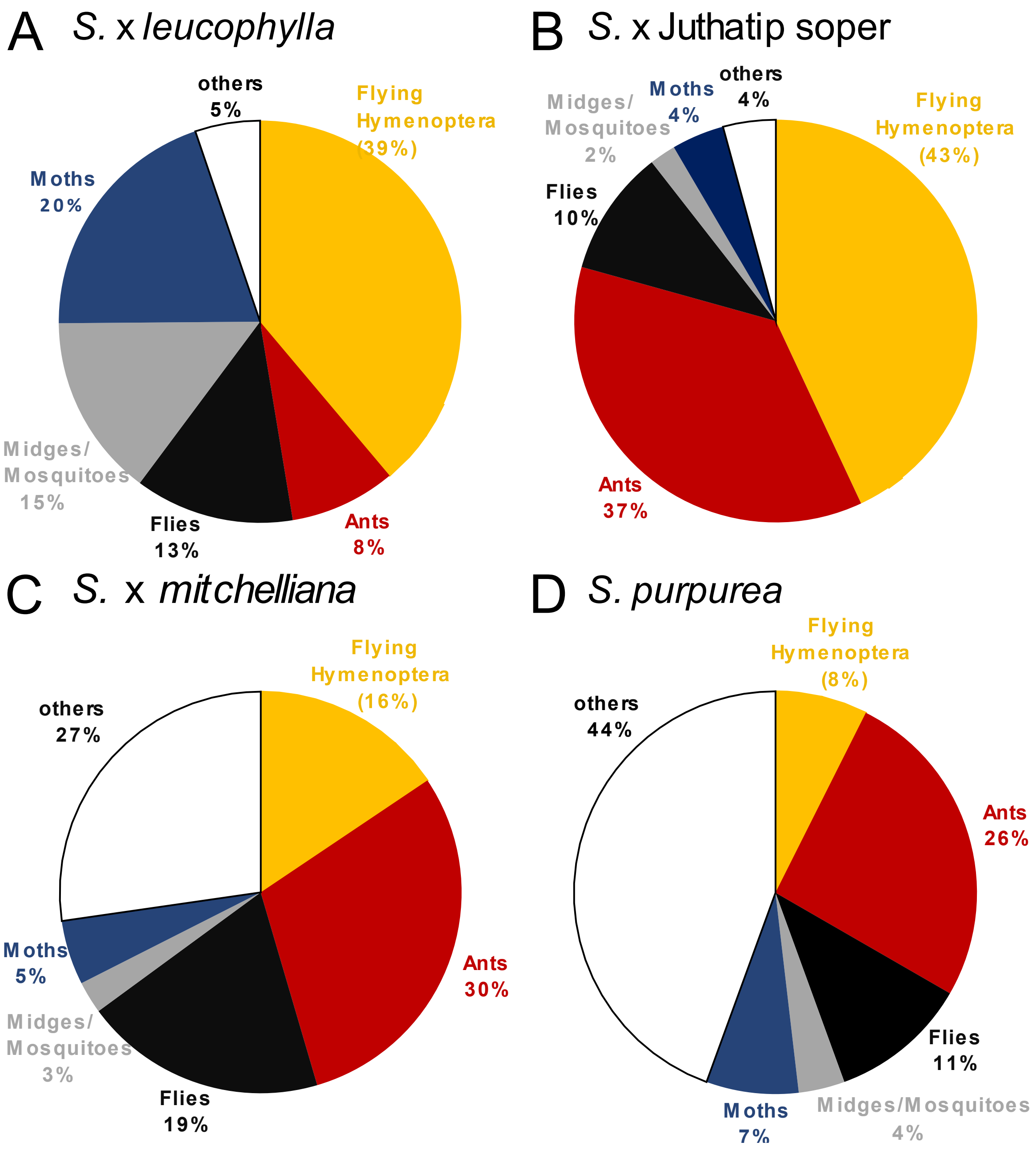


**Figure S6.** Total proportion of prey trapped in all pitchers for the four *Sarracenia* taxa: (A) *S.* x *leucophylla*, (B) *S.* x Juthatip soper, (C) *S.* x *mitchelliana*, (D) *S. purpurea*. Prey could be Diptera (Flies in black or Midges/Mosquitoes in grey), flying Hymenoptera (Bees, Bumblebees, Solitary Bees, Wasps and Parasitoid wasps in yellow), or crawling Hymenoptera (ants in red), Lepidoptera (Moths in blue) or other prey (in white, including Coleoptera, Hemiptera, Blattodea, Springtails, Snails and Spiders).

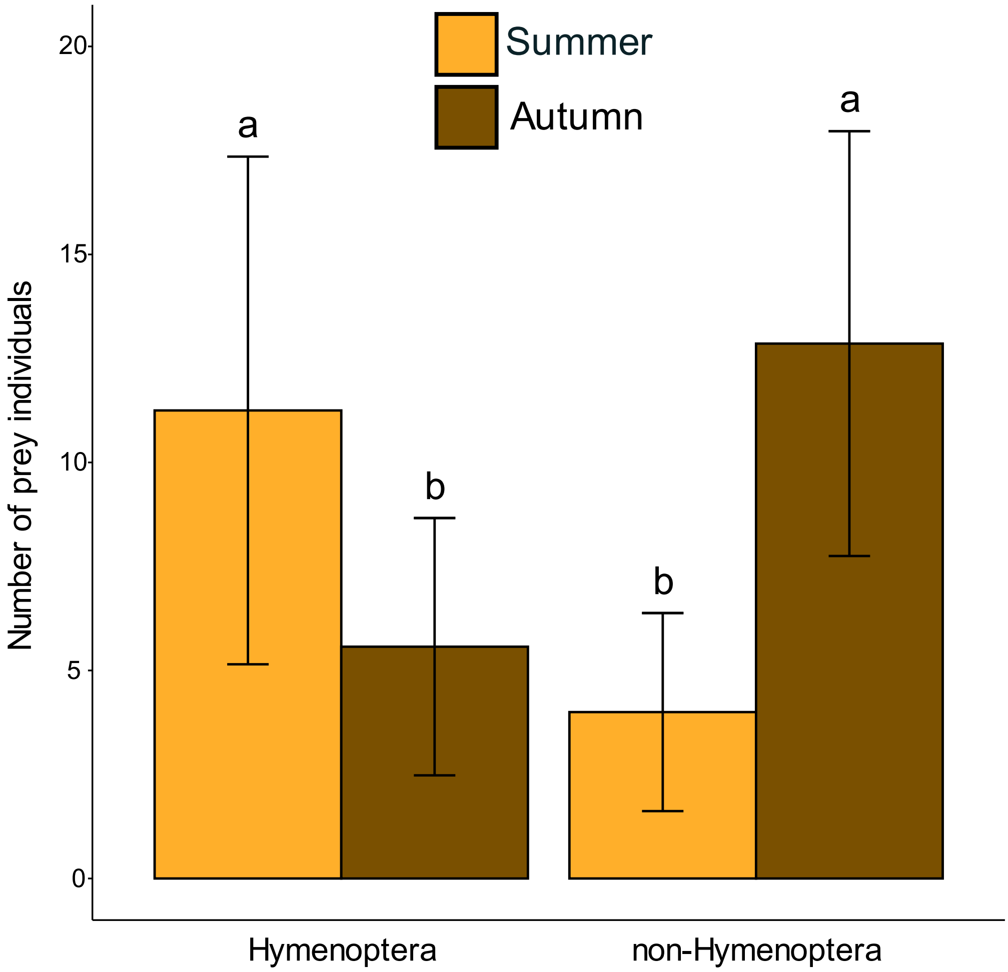


**Figure S7**. Seasonal variation in the number of Hymenoptera and non-Hymenoptera individuals captured. Mean values are presented with their associated standard errors. We consider only prey captured in pitchers at stage 2 because it was the only stage sampled for the two seasons (n = 4 in summer, n =7 in autumn). For each prey category, different letters above bars show statistically significant differences in means between seasons ($p<0.05$), as output by one-way ANOVAs.

**Table S1.** Results of the post-hoc tests associated to the best linear mixed model presented in Table 1 to explain the variation in colour contrast with a green background. The variable Colour contrast was square-root transformed to match residual normality. P-values were adjusted for multiple comparisons with the Bonferroni correction, *: p<0.05, **: p<0.01, and ***: p<0.001. Within a level of factor 1, we tested the contrast between two levels of factor 2 or between 2 gathered levels of factor 2. For instance, for areoles, we tested whether the colour contrast of *S.* x *leucophylla* was higher than the colour contrast of *S.* x Juthatip soper and it was not significantly different (p-value=0.585). Symbol "-" means that the contrasts were tested overall and not within a level of another factor. For instance, we tested whether the colour contrast of *S.* x *leucophylla* was higher overall than the colour contrast of *S.* x Juthatip soper.

| Level on factor 1 | Contrast on factor 2 | Coefficient (± standard error) | t-ratio | adjusted p-value | |
|---|---|---|---|---|---|
| - | *S.* x *leucophylla* > *S.* x Juthatip soper | 0.08 (±0.15) | 0.51 | 1.0000 | |
| - | *S.* x *leucophylla* > *S.* x *mitchelliana* | 0.08 (±0.15) | 0.55 | 1.0000 | |
| - | *S.* x *leucophylla* > *S. purpurea* | 0.11 (±0.15) | 0.74 | 1.0000 | |
| - | *S.* x Juthatip soper > *S.* x *mitchelliana* | 0.01 (±0.15) | 0.04 | 1.0000 | |
| - | *S.* x Juthatip soper > *S. purpurea* | 0.03 (±0.15) | 0.22 | 1.0000 | |
| - | *S.* x *mitchelliana* > *S. purpurea* | 0.03 (±0.15) | 0.18 | 1.0000 | |
| - | Areoles > Veins | 0.23 (±0.04) | 6.22 | <0.001 | *** |
| - | Areoles > Peristome | 0.08 (±0.04) | 1.68 | 0.5605 | |
| - | Areoles > Tube | 0.47 (±0.04) | 10.60 | <0.001 | *** |
| - | Veins > Peristome | -0.15 (±0.04) | -3.37 | 0.0048 | ** |
| - | Veins > Tube | 0.24 (±0.04) | 5.52 | <0.001 | *** |
| - | Peristome > Tube | 0.4 (±0.05) | 7.67 | <0.001 | *** |
| - | Stage 1 > Stage 2 | 0.08 (±0.18) | 0.43 | 1.0000 | |
| - | Stage 1 > Stage 3 | -0.15 (±0.15) | -1.02 | 1.0000 | |
| - | Stage 1 > Stage 4 | -0.24 (±0.18) | -1.34 | 1.0000 | |
| - | Stage 2 > Stage 3 | -0.23 (±0.15) | -1.55 | 0.8456 | |
| - | Stage 2 > Stage 4 | -0.32 (±0.18) | -1.77 | 0.5784 | |
| - | Stage 3 > Stage 4 | -0.09 (±0.15) | -0.62 | 1.0000 | |
| - | autumn > summer | 0.87 (±0.15) | 5.64 | <0.001 | *** |
| Areoles | *S.* x *leucophylla* > *S.* x Juthatip soper | 0.29 (±0.16) | 1.81 | 0.5085 | |
| Areoles | *S.* x *leucophylla* > *S.* x *mitchelliana* | 1 (±0.16) | 6.32 | <0.001 | *** |
| Areoles | *S.* x *leucophylla* > *S. purpurea* | 0.98 (±0.16) | 6.20 | <0.001 | *** |
| Areoles | *S.* x Juthatip soper > *S.* x *mitchelliana* | 0.71 (±0.16) | 4.53 | 0.0012 | ** |
| Areoles | *S.* x Juthatip soper > *S. purpurea* | 0.69 (±0.16) | 4.40 | 0.0016 | ** |
| Areoles | *S.* x *mitchelliana* > *S. purpurea* | -0.02 (±0.16) | -0.13 | 1.0000 | |

| | | | | | |
|---|---|---|---|---|---|
| Veins | *S.* x *leucophylla* > *S.* x Juthatip soper | -0.15 (±0.16) | -0.94 | 1.0000 | |
| Veins | *S.* x *leucophylla* > *S.* x *mitchelliana* | -0.51 (±0.16) | -3.24 | 0.0241 | * |
| Veins | *S.* x *leucophylla* > *S. purpurea* | -0.25 (±0.16) | -1.58 | 0.7796 | |
| Veins | *S.* x Juthatip soper > *S.* x *mitchelliana* | -0.36 (±0.16) | -2.31 | 0.1898 | |
| Veins | *S.* x Juthatip soper > *S. purpurea* | -0.1 (±0.16) | -0.64 | 1.0000 | |
| Veins | *S.* x *mitchelliana* > *S. purpurea* | 0.26 (±0.16) | 1.67 | 0.6621 | |
| Peristome | *S.* x *leucophylla* > *S.* x Juthatip soper | 0.24 (±0.17) | 1.41 | 1.0000 | |
| Peristome | *S.* x *leucophylla* > *S.* x *mitchelliana* | 0.08 (±0.17) | 0.47 | 1.0000 | |
| Peristome | *S.* x *leucophylla* > *S. purpurea* | -0.03 (±0.17) | -0.15 | 1.0000 | |
| Peristome | *S.* x Juthatip soper > *S.* x *mitchelliana* | -0.16 (±0.17) | -0.92 | 1.0000 | |
| Peristome | *S.* x Juthatip soper > *S. purpurea* | -0.27 (±0.17) | -1.56 | 0.7747 | |
| Peristome | *S.* x *mitchelliana* > *S. purpurea* | -0.11 (±0.17) | -0.62 | 1.0000 | |
| Tube | *S.* x *leucophylla* > *S.* x Juthatip soper | -0.08 (±0.17) | -0.47 | 1.0000 | |
| Tube | *S.* x *leucophylla* > *S.* x *mitchelliana* | -0.25 (±0.17) | -1.44 | 0.9696 | |
| Tube | *S.* x *leucophylla* > *S. purpurea* | -0.27 (±0.17) | -1.58 | 0.7498 | |
| Tube | *S.* x Juthatip soper > *S.* x *mitchelliana* | -0.17 (±0.17) | -0.96 | 1.0000 | |
| Tube | *S.* x Juthatip soper > *S. purpurea* | -0.19 (±0.17) | -1.10 | 1.0000 | |
| Tube | *S.* x *mitchelliana* > *S. purpurea* | -0.02 (±0.17) | -0.14 | 1.0000 | |
| *S.* x *leucophylla* | Areoles > (Veins, Peristome, Tube) | 0.92 (±0.07) | 14.02 | <0.001 | *** |
| *S.* x *leucophylla* | Veins > (Areoles, Peristome, Tube) | -0.43 (±0.07) | -6.67 | <0.001 | *** |
| *S.* x *leucophylla* | Peristome > (Areoles, Veins, Tube) | 0.17 (±0.08) | 2.11 | 0.1422 | |
| *S.* x *leucophylla* | Tube > (Areoles, Veins, Peristome) | -0.66 (±0.08) | -8.37 | <0.001 | *** |
| *S.* x Juthatip soper | Areoles > (Veins, Peristome, Tube) | 0.64 (±0.06) | 10.02 | <0.001 | *** |
| *S.* x Juthatip soper | Veins > (Areoles, Peristome, Tube) | -0.14 (±0.06) | -2.15 | 0.1260 | |
| *S.* x Juthatip soper | Peristome > (Areoles, Veins, Tube) | -0.06 (±0.08) | -0.69 | 1.0000 | |
| *S.* x Juthatip soper | Tube > (Areoles, Veins, Peristome) | -0.45 (±0.08) | -5.59 | <0.001 | *** |
| *S.* x *mitchelliana* | Areoles > (Veins, Peristome, Tube) | -0.3 (±0.06) | -4.62 | <0.001 | *** |
| *S.* x *mitchelliana* | Veins > (Areoles, Peristome, Tube) | 0.35 (±0.06) | 5.46 | <0.001 | *** |
| *S.* x *mitchelliana* | Peristome > (Areoles, Veins, Tube) | 0.17 (±0.08) | 2.00 | 0.1854 | |
| *S.* x *mitchelliana* | Tube > (Areoles, Veins, Peristome) | -0.22 (±0.08) | -2.74 | 0.0253 | * |
| S. *purpurea* | Areoles > (Veins, Peristome, Tube) | -0.24 (±0.06) | -3.71 | <0.001 | *** |
| S. *purpurea* | Veins > (Areoles, Peristome, Tube) | 0.04 (±0.06) | 0.63 | 1.0000 | |
| S. *purpurea* | Peristome > (Areoles, Veins, Tube) | 0.35 (±0.08) | 4.33 | <0.001 | *** |
| S. *purpurea* | Tube > (Areoles, Veins, Peristome) | -0.15 (±0.08) | -1.90 | 0.2331 | |
| Areoles | Stage 1 > (2, 3, 4) | -0.07 (±0.15) | -0.49 | 1.0000 | |
| Areoles | Stage 2 > (1, 3, 4) | 0.29 (±0.15) | 2.01 | 0.2415 | |
| Areoles | Stage 3 > (1, 2, 4) | -0.16 (±0.11) | -1.43 | 0.6787 | |
| Areoles | Stage 4 > (1, 2, 3) | -0.06 (±0.15) | -0.41 | 1.0000 | |
| Veins | Stage 1 > (2, 3, 4) | -0.37 (±0.15) | -2.41 | 0.1022 | |
| Veins | Stage 2 > (1, 3, 4) | -0.29 (±0.15) | -2.00 | 0.2470 | |
| Veins | Stage 3 > (1, 2, 4) | 0.27 (±0.11) | 2.48 | 0.0897 | |
| Veins | Stage 4 > (1, 2, 3) | 0.39 (±0.15) | 2.54 | 0.0778 | |

| | | | | | |
|---|---|---|---|---|---|
| Peristome | Stage 1 > (2, 3, 4) | -0.14 (±0.17) | -0.84 | 1.0000 | |
| Peristome | Stage 2 > (1, 3, 4) | -0.7 (±0.15) | -4.50 | <0.001 | *** |
| Peristome | Stage 3 > (1, 2, 4) | 0.26 (±0.12) | 2.18 | 0.1501 | |
| Peristome | Stage 4 > (1, 2, 3) | 0.57 (±0.17) | 3.42 | 0.0074 | ** |
| Tube | Stage 1 > (2, 3, 4) | 0.17 (±0.17) | 1.01 | 1.0000 | |
| Tube | Stage 2 > (1, 3, 4) | -0.13 (±0.15) | -0.86 | 1.0000 | |
| Tube | Stage 3 > (1, 2, 4) | 0 (±0.12) | 0.01 | 1.0000 | |
| Tube | Stage 4 > (1, 2, 3) | -0.04 (±0.17) | -0.22 | 1.0000 | |
| Stage 1 | Areoles > (Veins, Peristome, Tube) | 0.29 (±0.08) | 3.63 | 0.0012 | ** |
| Stage 1 | Veins > (Areoles, Peristome, Tube) | -0.31 (±0.08) | -3.94 | <0.001 | *** |
| Stage 1 | Peristome > (Areoles, Veins, Tube) | 0.12 (±0.1) | 1.22 | 0.8938 | |
| Stage 1 | Tube > (Areoles, Veins, Peristome) | -0.1 (±0.1) | -1.03 | 1.0000 | |
| Stage 2 | Areoles > (Veins, Peristome, Tube) | 0.76 (±0.05) | 16.72 | <0.001 | *** |
| Stage 2 | Veins > (Areoles, Peristome, Tube) | -0.13 (±0.05) | -2.87 | 0.0166 | * |
| Stage 2 | Peristome > (Areoles, Veins, Tube) | -0.33 (±0.06) | -5.85 | <0.001 | *** |
| Stage 2 | Tube > (Areoles, Veins, Peristome) | -0.3 (±0.06) | -5.20 | <0.001 | *** |
| Stage 3 | Areoles > (Veins, Peristome, Tube) | 0.01 (±0.05) | 0.12 | 1.0000 | |
| Stage 3 | Veins > (Areoles, Peristome, Tube) | 0.13 (±0.05) | 2.93 | 0.0138 | * |
| Stage 3 | Peristome > (Areoles, Veins, Tube) | 0.32 (±0.06) | 5.74 | <0.001 | *** |
| Stage 3 | Tube > (Areoles, Veins, Peristome) | -0.46 (±0.06) | -8.15 | <0.001 | *** |
| Stage 4 | Areoles > (Veins, Peristome, Tube) | -0.02 (±0.08) | -0.26 | 1.0000 | |
| Stage 4 | Veins > (Areoles, Peristome, Tube) | 0.13 (±0.08) | 1.60 | 0.4445 | |
| Stage 4 | Peristome > (Areoles, Veins, Tube) | 0.52 (±0.1) | 5.11 | <0.001 | *** |
| Stage 4 | Tube > (Areoles, Veins, Peristome) | -0.62 (±0.1) | -6.29 | <0.001 | *** |
| *S.* x *leucophylla* | Stage 1 > (2, 3, 4) | -0.24 (±0.28) | -0.88 | 1.0000 | |
| *S.* x *leucophylla* | Stage 2 > (1, 3, 4) | -0.09 (±0.22) | -0.41 | 1.0000 | |
| *S.* x *leucophylla* | Stage 3 > (1, 2, 4) | 0.07 (±0.2) | 0.37 | 1.0000 | |
| *S.* x *leucophylla* | Stage 4 > (1, 2, 3) | 0.26 (±0.28) | 0.94 | 1.0000 | |
| *S.* x Juthatip soper | Stage 1 > (2, 3, 4) | 0 (±0.28) | -0.01 | 1.0000 | |
| *S.* x Juthatip soper | Stage 2 > (1, 3, 4) | -0.16 (±0.22) | -0.71 | 1.0000 | |
| *S.* x Juthatip soper | Stage 3 > (1, 2, 4) | 0.4 (±0.2) | 2.00 | 0.2542 | |
| *S.* x Juthatip soper | Stage 4 > (1, 2, 3) | -0.24 (±0.28) | -0.85 | 1.0000 | |
| *S.* x *mitchelliana* | Stage 1 > (2, 3, 4) | 0.04 (±0.28) | 0.14 | 1.0000 | |
| *S.* x *mitchelliana* | Stage 2 > (1, 3, 4) | -0.28 (±0.22) | -1.28 | 0.8834 | |
| *S.* x *mitchelliana* | Stage 3 > (1, 2, 4) | -0.01 (±0.2) | -0.05 | 1.0000 | |
| *S.* x *mitchelliana* | Stage 4 > (1, 2, 3) | 0.25 (±0.28) | 0.91 | 1.0000 | |
| S. *purpurea* | Stage 1 > (2, 3, 4) | -0.21 (±0.28) | -0.75 | 1.0000 | |
| S. *purpurea* | Stage 2 > (1, 3, 4) | -0.3 (±0.22) | -1.37 | 0.7655 | |
| S. *purpurea* | Stage 3 > (1, 2, 4) | -0.08 (±0.2) | -0.39 | 1.0000 | |
| S. *purpurea* | Stage 4 > (1, 2, 3) | 0.58 (±0.28) | 2.11 | 0.2081 | |
| Stage 1 | *S.* x *leucophylla* > *S.* x Juthatip soper | -0.1 (±0.36) | -0.29 | 1.0000 | |
| Stage 1 | *S.* x *leucophylla* > *S.* x *mitchelliana* | -0.13 (±0.36) | -0.36 | 1.0000 | |

| | | | | |
|---|---|---|---|---|
| Stage 1 | *S.* x *leucophylla* > *S. purpurea* | 0.08 (±0.36) | 0.23 | 1.0000 |
| Stage 1 | *S.* x Juthatip soper > *S.* x *mitchelliana* | -0.03 (±0.36) | -0.07 | 1.0000 |
| Stage 1 | *S.* x Juthatip soper > *S. purpurea* | 0.18 (±0.36) | 0.52 | 1.0000 |
| Stage 1 | *S.* x *mitchelliana* > *S. purpurea* | 0.21 (±0.36) | 0.59 | 1.0000 |
| Stage 2 | *S.* x *leucophylla* > *S.* x Juthatip soper | 0.12 (±0.21) | 0.60 | 1.0000 |
| Stage 2 | *S.* x *leucophylla* > *S.* x *mitchelliana* | 0.22 (±0.21) | 1.09 | 1.0000 |
| Stage 2 | *S.* x *leucophylla* > *S. purpurea* | 0.27 (±0.21) | 1.29 | 1.0000 |
| Stage 2 | *S.* x Juthatip soper > *S.* x *mitchelliana* | 0.1 (±0.21) | 0.48 | 1.0000 |
| Stage 2 | *S.* x Juthatip soper > *S. purpurea* | 0.14 (±0.21) | 0.69 | 1.0000 |
| Stage 2 | *S.* x *mitchelliana* > *S. purpurea* | 0.04 (±0.21) | 0.20 | 1.0000 |
| Stage 3 | *S.* x *leucophylla* > *S.* x Juthatip soper | -0.17 (±0.21) | -0.81 | 1.0000 |
| Stage 3 | *S.* x *leucophylla* > *S.* x *mitchelliana* | 0.14 (±0.21) | 0.69 | 1.0000 |
| Stage 3 | *S.* x *leucophylla* > *S. purpurea* | 0.22 (±0.21) | 1.06 | 1.0000 |
| Stage 3 | *S.* x Juthatip soper > *S.* x *mitchelliana* | 0.31 (±0.21) | 1.50 | 0.9213 |
| Stage 3 | *S.* x Juthatip soper > *S. purpurea* | 0.39 (±0.21) | 1.87 | 0.4831 |
| Stage 3 | *S.* x *mitchelliana* > *S. purpurea* | 0.08 (±0.21) | 0.37 | 1.0000 |
| Stage 4 | *S.* x *leucophylla* > *S.* x Juthatip soper | 0.45 (±0.36) | 1.25 | 1.0000 |
| Stage 4 | *S.* x *leucophylla* > *S.* x *mitchelliana* | 0.08 (±0.36) | 0.24 | 1.0000 |
| Stage 4 | *S.* x *leucophylla* > *S. purpurea* | -0.14 (±0.36) | -0.38 | 1.0000 |
| Stage 4 | *S.* x Juthatip soper > *S.* x *mitchelliana* | -0.36 (±0.36) | -1.01 | 1.0000 |
| Stage 4 | *S.* x Juthatip soper > *S. purpurea* | -0.58 (±0.36) | -1.63 | 0.7396 |
| Stage 4 | *S.* x *mitchelliana* > *S. purpurea* | -0.22 (±0.36) | -0.62 | 1.0000 |

**Table S2.** Results of the post-hoc tests associated to the best linear mixed model presented in Table 1 to explain the variation in brightness contrast with a green background. The variable Brightness contrast was square-root transformed to match residual normality. P-values were adjusted for multiple comparisons with the Bonferroni correction, *: p<0.05, **: p<0.01, and ***: p<0.001. Within a level of factor 1, we tested the contrast between two levels of factor 2 or between 2 gathered levels of factor 2. For instance, for areoles, we tested whether the brightness contrast of *S.* x *leucophylla* was higher than the brightness contrast of *S.* x Juthatip soper and we found that it was significantly different (associated with a corrected p-value=0.0034). Symbol "-" means that the contrasts were tested overall and not within a level of another factor. For instance, we tested whether the brightness contrast of *S.* x *leucophylla* was higher than the brightness contrast of *S.* x Juthatip soper overall.

| Level on factor 1 | Contrast on factor 2 | Coefficient (± standard error) | t-ratio | adjusted p-value | |
|---|---|---|---|---|---|
| - | *S.* x *leucophylla* > *S.* x Juthatip soper | 0.32 (±0.14) | 2.21 | 0.2481 | |
| - | *S.* x *leucophylla* > *S.* x *mitchelliana* | 0.23 (±0.14) | 1.57 | 0.8025 | |
| - | *S.* x *leucophylla* > *S. purpurea* | 0.14 (±0.14) | 0.95 | 1.0000 | |
| - | *S.* x Juthatip soper > *S.* x *mitchelliana* | -0.09 (±0.14) | -0.63 | 1.0000 | |
| - | *S.* x Juthatip soper > *S. purpurea* | -0.18 (±0.14) | -1.26 | 1.0000 | |
| - | *S.* x *mitchelliana* > *S. purpurea* | -0.09 (±0.14) | -0.63 | 1.0000 | |
| - | Areoles > Veins | 0.5 (±0.07) | 7.63 | <0.001 | *** |
| - | Areoles > Peristome | 0.35 (±0.08) | 4.28 | <0.001 | *** |
| - | Areoles > Tube | 0.45 (±0.08) | 5.65 | <0.001 | *** |
| - | Veins > Peristome | -0.15 (±0.08) | -1.91 | 0.3381 | |
| - | Veins > Tube | -0.05 (±0.08) | -0.61 | 1.0000 | |
| - | Peristome > Tube | 0.11 (±0.09) | 1.14 | 1.0000 | |
| - | Stage 1 > Stage 2 | 0.17 (±0.14) | 1.21 | 1.0000 | |
| - | Stage 1 > Stage 3 | 0.06 (±0.14) | 0.44 | 1.0000 | |
| - | Stage 1 > Stage 4 | 0.25 (±0.18) | 1.44 | 1.0000 | |
| - | Stage 2 > Stage 3 | -0.11 (±0.1) | -1.08 | 1.0000 | |
| - | Stage 2 > Stage 4 | 0.08 (±0.14) | 0.56 | 1.0000 | |
| - | Stage 3 > Stage 4 | 0.19 (±0.14) | 1.32 | 1.0000 | |
| Areoles | *S.* x *leucophylla* > *S.* x Juthatip soper | 0.67 (±0.18) | 3.74 | 0.0034 | ** |
| Areoles | *S.* x *leucophylla* > *S.* x *mitchelliana* | 0.95 (±0.18) | 5.27 | <0.001 | *** |
| Areoles | *S.* x *leucophylla* > *S. purpurea* | 0.81 (±0.18) | 4.52 | <0.001 | *** |
| Areoles | *S.* x Juthatip soper > *S.* x *mitchelliana* | 0.28 (±0.18) | 1.55 | 0.7744 | |
| Areoles | *S.* x Juthatip soper > *S. purpurea* | 0.14 (±0.18) | 0.77 | 1.0000 | |
| Areoles | *S.* x *mitchelliana* > *S. purpurea* | -0.14 (±0.18) | -0.78 | 1.0000 | |

| | | | | | |
|---|---|---|---|---|---|
| Veins | *S.* x *leucophylla* > *S.* x Juthatip soper | -0.01 (±0.18) | -0.04 | 1.0000 | |
| Veins | *S.* x *leucophylla* > *S.* x *mitchelliana* | -0.02 (±0.18) | -0.11 | 1.0000 | |
| Veins | *S.* x *leucophylla* > *S. purpurea* | -0.3 (±0.18) | -1.68 | 0.6086 | |
| Veins | *S.* x Juthatip soper > *S.* x *mitchelliana* | -0.01 (±0.18) | -0.07 | 1.0000 | |
| Veins | *S.* x Juthatip soper > *S. purpurea* | -0.29 (±0.18) | -1.66 | 0.6352 | |
| Veins | *S.* x *mitchelliana* > *S. purpurea* | -0.28 (±0.18) | -1.58 | 0.7395 | |
| Peristome | *S.* x *leucophylla* > *S.* x Juthatip soper | 0.54 (±0.22) | 2.44 | 0.1009 | |
| Peristome | *S.* x *leucophylla* > *S.* x *mitchelliana* | 0.09 (±0.22) | 0.39 | 1.0000 | |
| Peristome | *S.* x *leucophylla* > *S. purpurea* | 0.14 (±0.22) | 0.65 | 1.0000 | |
| Peristome | *S.* x Juthatip soper > *S.* x *mitchelliana* | -0.45 (±0.22) | -2.00 | 0.2879 | |
| Peristome | *S.* x Juthatip soper > *S. purpurea* | -0.39 (±0.22) | -1.78 | 0.4682 | |
| Peristome | *S.* x *mitchelliana* > *S. purpurea* | 0.06 (±0.22) | 0.25 | 1.0000 | |
| Tube | *S.* x *leucophylla* > *S.* x Juthatip soper | 0.07 (±0.22) | 0.32 | 1.0000 | |
| Tube | *S.* x *leucophylla* > *S.* x *mitchelliana* | -0.11 (±0.22) | -0.49 | 1.0000 | |
| Tube | *S.* x *leucophylla* > *S. purpurea* | -0.11 (±0.22) | -0.49 | 1.0000 | |
| Tube | *S.* x Juthatip soper > *S.* x *mitchelliana* | -0.18 (±0.22) | -0.79 | 1.0000 | |
| Tube | *S.* x Juthatip soper > *S. purpurea* | -0.18 (±0.22) | -0.80 | 1.0000 | |
| Tube | *S.* x *mitchelliana* > *S. purpurea* | 0 (±0.22) | 0.00 | 1.0000 | |
| *S.* x *leucophylla* | Areoles > (Veins, Peristome, Tube) | 1.02 (±0.12) | 8.57 | <0.001 | *** |
| *S.* x *leucophylla* | Veins > (Areoles, Peristome, Tube) | -0.57 (±0.12) | -4.87 | <0.001 | *** |
| *S.* x *leucophylla* | Peristome > (Areoles, Veins, Tube) | 0 (±0.15) | 0.00 | 1.0000 | |
| *S.* x *leucophylla* | Tube > (Areoles, Veins, Peristome) | -0.45 (±0.14) | -3.14 | 0.0070 | ** |
| *S.* x Juthatip soper | Areoles > (Veins, Peristome, Tube) | 0.54 (±0.12) | 4.71 | <0.001 | *** |
| *S.* x Juthatip soper | Veins > (Areoles, Peristome, Tube) | -0.14 (±0.11) | -1.20 | 0.9151 | |
| *S.* x Juthatip soper | Peristome > (Areoles, Veins, Tube) | -0.29 (±0.14) | -2.02 | 0.1733 | |
| *S.* x Juthatip soper | Tube > (Areoles, Veins, Peristome) | -0.11 (±0.15) | -0.78 | 1.0000 | |
| *S.* x *mitchelliana* | Areoles > (Veins, Peristome, Tube) | 0.06 (±0.12) | 0.48 | 1.0000 | |
| *S.* x *mitchelliana* | Veins > (Areoles, Peristome, Tube) | -0.24 (±0.12) | -2.07 | 0.1547 | |
| *S.* x *mitchelliana* | Peristome > (Areoles, Veins, Tube) | 0.19 (±0.15) | 1.23 | 0.8698 | |
| *S.* x *mitchelliana* | Tube > (Areoles, Veins, Peristome) | 0 (±0.15) | 0.00 | 1.0000 | |
| S. *purpurea* | Areoles > (Veins, Peristome, Tube) | 0.12 (±0.11) | 1.05 | 1.0000 | |
| S. *purpurea* | Veins > (Areoles, Peristome, Tube) | 0.01 (±0.11) | 0.09 | 1.0000 | |
| S. *purpurea* | Peristome > (Areoles, Veins, Tube) | -0.01 (±0.14) | -0.07 | 1.0000 | |
| S. *purpurea* | Tube > (Areoles, Veins, Peristome) | -0.12 (±0.14) | -0.83 | 1.0000 | |
| Areoles | Stage 1 > (2, 3, 4) | 0.39 (±0.17) | 2.29 | 0.1089 | |
| Areoles | Stage 2 > (1, 3, 4) | -0.28 (±0.12) | -2.37 | 0.0916 | |
| Areoles | Stage 3 > (1, 2, 4) | 0.03 (±0.12) | 0.22 | 1.0000 | |
| Areoles | Stage 4 > (1, 2, 3) | -0.13 (±0.17) | -0.79 | 1.0000 | |
| Veins | Stage 1 > (2, 3, 4) | 0.08 (±0.17) | 0.47 | 1.0000 | |
| Veins | Stage 2 > (1, 3, 4) | 0.49 (±0.12) | 4.18 | <0.001 | *** |
| Veins | Stage 3 > (1, 2, 4) | -0.11 (±0.12) | -0.89 | 1.0000 | |
| Veins | Stage 4 > (1, 2, 3) | -0.47 (±0.17) | -2.79 | 0.0326 | * |

| | | | | | |
|---|---|---|---|---|---|
| Peristome | Stage 1 > (2, 3, 4) | -0.26 (±0.21) | -1.26 | 0.8413 | |
| Peristome | Stage 2 > (1, 3, 4) | 0.11 (±0.15) | 0.75 | 1.0000 | |
| Peristome | Stage 3 > (1, 2, 4) | -0.02 (±0.15) | -0.13 | 1.0000 | |
| Peristome | Stage 4 > (1, 2, 3) | 0.17 (±0.21) | 0.81 | 1.0000 | |
| Tube | Stage 1 > (2, 3, 4) | 0.46 (±0.21) | 2.21 | 0.1186 | |
| Tube | Stage 2 > (1, 3, 4) | -0.6 (±0.15) | -4.05 | <0.001 | *** |
| Tube | Stage 3 > (1, 2, 4) | 0.41 (±0.15) | 2.82 | 0.0235 | * |
| Tube | Stage 4 > (1, 2, 3) | -0.27 (±0.21) | -1.32 | 0.7619 | |
| Stage 1 | Areoles > (Veins, Peristome, Tube) | 0.66 (±0.14) | 4.61 | <0.001 | *** |
| Stage 1 | Veins > (Areoles, Peristome, Tube) | -0.32 (±0.14) | -2.29 | 0.0899 | |
| Stage 1 | Peristome > (Areoles, Veins, Tube) | -0.46 (±0.18) | -2.57 | 0.0411 | * |
| Stage 1 | Tube > (Areoles, Veins, Peristome) | 0.12 (±0.17) | 0.69 | 1.0000 | |
| Stage 2 | Areoles > (Veins, Peristome, Tube) | 0.22 (±0.08) | 2.70 | 0.0286 | * |
| Stage 2 | Veins > (Areoles, Peristome, Tube) | 0.33 (±0.08) | 4.00 | <0.001 | *** |
| Stage 2 | Peristome > (Areoles, Veins, Tube) | 0.15 (±0.1) | 1.46 | 0.5846 | |
| Stage 2 | Tube > (Areoles, Veins, Peristome) | -0.7 (±0.1) | -6.80 | <0.001 | *** |
| Stage 3 | Areoles > (Veins, Peristome, Tube) | 0.38 (±0.08) | 4.66 | <0.001 | *** |
| Stage 3 | Veins > (Areoles, Peristome, Tube) | -0.42 (±0.08) | -5.12 | <0.001 | *** |
| Stage 3 | Peristome > (Areoles, Veins, Tube) | -0.13 (±0.1) | -1.25 | 0.8488 | |
| Stage 3 | Tube > (Areoles, Veins, Peristome) | 0.17 (±0.1) | 1.61 | 0.4280 | |
| Stage 4 | Areoles > (Veins, Peristome, Tube) | 0.48 (±0.14) | 3.35 | 0.0034 | ** |
| Stage 4 | Veins > (Areoles, Peristome, Tube) | -0.53 (±0.14) | -3.69 | <0.001 | *** |
| Stage 4 | Peristome > (Areoles, Veins, Tube) | 0.32 (±0.18) | 1.75 | 0.3245 | |
| Stage 4 | Tube > (Areoles, Veins, Peristome) | -0.27 (±0.18) | -1.51 | 0.5278 | |
| *S.* x *leucophylla* | Stage 1 > (2, 3, 4) | -0.27 (±0.27) | -1.01 | 1.0000 | |
| *S.* x *leucophylla* | Stage 2 > (1, 3, 4) | -0.24 (±0.19) | -1.27 | 0.8835 | |
| *S.* x *leucophylla* | Stage 3 > (1, 2, 4) | 0.19 (±0.19) | 1.00 | 1.0000 | |
| *S.* x *leucophylla* | Stage 4 > (1, 2, 3) | 0.33 (±0.27) | 1.20 | 0.9928 | |
| *S.* x Juthatip soper | Stage 1 > (2, 3, 4) | 0.13 (±0.27) | 0.49 | 1.0000 | |
| *S.* x Juthatip soper | Stage 2 > (1, 3, 4) | -0.03 (±0.19) | -0.17 | 1.0000 | |
| *S.* x Juthatip soper | Stage 3 > (1, 2, 4) | 0.15 (±0.19) | 0.77 | 1.0000 | |
| *S.* x Juthatip soper | Stage 4 > (1, 2, 3) | -0.25 (±0.27) | -0.91 | 1.0000 | |
| *S.* x *mitchelliana* | Stage 1 > (2, 3, 4) | 0.21 (±0.27) | 0.79 | 1.0000 | |
| *S.* x *mitchelliana* | Stage 2 > (1, 3, 4) | -0.27 (±0.19) | -1.39 | 0.7262 | |
| *S.* x *mitchelliana* | Stage 3 > (1, 2, 4) | 0.25 (±0.19) | 1.31 | 0.8256 | |
| *S.* x *mitchelliana* | Stage 4 > (1, 2, 3) | -0.2 (±0.28) | -0.72 | 1.0000 | |
| S. *purpurea* | Stage 1 > (2, 3, 4) | 0.58 (±0.27) | 2.15 | 0.1859 | |
| S. *purpurea* | Stage 2 > (1, 3, 4) | 0.27 (±0.19) | 1.43 | 0.6840 | |
| S. *purpurea* | Stage 3 > (1, 2, 4) | -0.28 (±0.19) | -1.44 | 0.6723 | |
| S. *purpurea* | Stage 4 > (1, 2, 3) | -0.58 (±0.27) | -2.14 | 0.1886 | |
| Stage 1 | *S.* x *leucophylla* > *S.* x Juthatip soper | 0.01 (±0.35) | 0.04 | 1.0000 | |
| Stage 1 | *S.* x *leucophylla* > *S.* x *mitchelliana* | -0.14 (±0.35) | -0.39 | 1.0000 | |

| | | | | |
|---|---|---|---|---|
| Stage 1 | *S.* x *leucophylla* > *S. purpurea* | -0.51 (±0.35) | -1.44 | 1.0000 |
| Stage 1 | *S.* x Juthatip soper > *S.* x *mitchelliana* | -0.15 (±0.35) | -0.43 | 1.0000 |
| Stage 1 | *S.* x Juthatip soper > *S. purpurea* | -0.52 (±0.35) | -1.47 | 0.9583 |
| Stage 1 | *S.* x *mitchelliana* > *S. purpurea* | -0.37 (±0.35) | -1.04 | 1.0000 |
| Stage 2 | *S.* x *leucophylla* > *S.* x Juthatip soper | 0.16 (±0.2) | 0.78 | 1.0000 |
| Stage 2 | *S.* x *leucophylla* > *S.* x *mitchelliana* | 0.25 (±0.2) | 1.20 | 1.0000 |
| Stage 2 | *S.* x *leucophylla* > *S. purpurea* | -0.25 (±0.2) | -1.24 | 1.0000 |
| Stage 2 | *S.* x Juthatip soper > *S.* x *mitchelliana* | 0.09 (±0.2) | 0.42 | 1.0000 |
| Stage 2 | *S.* x Juthatip soper > *S. purpurea* | -0.41 (±0.2) | -2.02 | 0.3586 |
| Stage 2 | *S.* x *mitchelliana* > *S. purpurea* | -0.5 (±0.2) | -2.44 | 0.1550 |
| Stage 3 | *S.* x *leucophylla* > *S.* x Juthatip soper | 0.35 (±0.2) | 1.72 | 0.6201 |
| Stage 3 | *S.* x *leucophylla* > *S.* x *mitchelliana* | 0.18 (±0.2) | 0.89 | 1.0000 |
| Stage 3 | *S.* x *leucophylla* > *S. purpurea* | 0.49 (±0.2) | 2.39 | 0.1708 |
| Stage 3 | *S.* x Juthatip soper > *S.* x *mitchelliana* | -0.17 (±0.2) | -0.83 | 1.0000 |
| Stage 3 | *S.* x Juthatip soper > *S. purpurea* | 0.14 (±0.2) | 0.68 | 1.0000 |
| Stage 3 | *S.* x *mitchelliana* > *S. purpurea* | 0.31 (±0.2) | 1.50 | 0.9048 |
| Stage 4 | *S.* x *leucophylla* > *S.* x Juthatip soper | 0.75 (±0.35) | 2.12 | 0.2950 |
| Stage 4 | *S.* x *leucophylla* > *S.* x *mitchelliana* | 0.62 (±0.36) | 1.74 | 0.5913 |
| Stage 4 | *S.* x *leucophylla* > *S. purpurea* | 0.82 (±0.35) | 2.31 | 0.2012 |
| Stage 4 | *S.* x Juthatip soper > *S.* x *mitchelliana* | -0.13 (±0.36) | -0.36 | 1.0000 |
| Stage 4 | *S.* x Juthatip soper > *S. purpurea* | 0.07 (±0.35) | 0.19 | 1.0000 |
| Stage 4 | *S.* x *mitchelliana* > *S. purpurea* | 0.2 (±0.36) | 0.55 | 1.0000 |

**Table S3.** Results of the post-hoc tests associated to the best linear mixed model, presented in Table 2, to explain the variation in colour contrast between any two pitcher areas. The variable colour contrast was square-root transformed to match residual normality. P-values were adjusted for multiple comparisons with the Bonferroni correction. Symbols describe various levels of p-values *: p<0.05, **: p<0.01, and ***: p<0.001. Within a level of factor 1, we tested the oriented contrast between two levels of factor 2 or between 2 grouped levels of factor 2. For instance, for the contrast produced by areoles against veins, we tested whether the colour contrast of *S.* x *leucophylla* was higher than the colour contrast of *S.* x Juthatip soper and it was significantly different (p-value=0.0039). Symbols "-" mean that the oriented contrasts were tested overall and not within a level of another factor. For instance, we tested whether the colour contrast of *S.* x *leucophylla* was higher than the colour contrast of *S.* x Juthatip soper overall.

| Level on factor 1 | Contrast on factor 2 | Coefficient (± standard error) | t-ratio | adjusted p-value | |
|---|---|---|---|---|---|
| - | *S.* x *leucophylla* > *S.* x Juthatip soper | 0.31 (±0.1) | 3.11 | 0.0427 | * |
| - | *S.* x *leucophylla* > *S.* x *mitchelliana* | 0.47 (±0.1) | 4.69 | 0.0017 | ** |
| - | *S.* x *leucophylla* > *S. purpurea* | 0.64 (±0.1) | 6.35 | <0.001 | *** |
| - | *S.* x Juthatip soper > *S.* x *mitchelliana* | 0.16 (±0.1) | 1.58 | 0.8049 | |
| - | *S.* x Juthatip soper > *S. purpurea* | 0.33 (±0.1) | 3.24 | 0.0327 | * |
| - | *S.* x *mitchelliana* > *S. purpurea* | 0.17 (±0.1) | 1.66 | 0.7099 | |
| - | Stage 1 > Stage 2 | -0.05 (±0.12) | -0.41 | 1.0000 | |
| - | Stage 1 > Stage 3 | -0.03 (±0.1) | -0.32 | 1.0000 | |
| - | Stage 1 > Stage 4 | 0.03 (±0.12) | 0.26 | 1.0000 | |
| - | Stage 2 > Stage 3 | 0.02 (±0.1) | 0.18 | 1.0000 | |
| - | Stage 2 > Stage 4 | 0.08 (±0.12) | 0.67 | 1.0000 | |
| - | Stage 3 > Stage 4 | 0.06 (±0.1) | 0.64 | 1.0000 | |
| - | Areoles-Veins > Areoles-Peristome | -0.22 (±0.02) | -13.49 | <0.001 | *** |
| - | Areoles-Veins > Areoles-Tube | 0.16 (±0.02) | 10.16 | <0.001 | *** |
| - | Areoles-Veins > Veins-Peristome | 0.34 (±0.02) | 21.06 | <0.001 | *** |
| - | Areoles-Veins > Veins-Tube | 0.35 (±0.02) | 22.01 | <0.001 | *** |
| - | Areoles-Veins > Peristome-Tube | 0.03 (±0.02) | 1.34 | 1.0000 | |
| - | Areoles-Peristome > Areoles-Tube | 0.38 (±0.02) | 20.47 | <0.001 | *** |
| - | Areoles-Peristome > Veins-Peristome | 0.56 (±0.02) | 29.82 | <0.001 | *** |
| - | Areoles-Peristome > Veins-Tube | 0.57 (±0.02) | 30.68 | <0.001 | *** |
| - | Areoles-Peristome > Peristome-Tube | 0.25 (±0.02) | 10.80 | <0.001 | *** |
| - | Areoles-Tube > Veins-Peristome | 0.18 (±0.02) | 9.54 | <0.001 | *** |
| - | Areoles-Tube > Veins-Tube | 0.19 (±0.02) | 10.23 | <0.001 | *** |
| - | Areoles-Tube > Peristome-Tube | -0.14 (±0.02) | -5.95 | <0.001 | *** |

| | | | | | |
|---|---|---|---|---|---|
| - | Veins-Peristome > Veins-Tube | 0.01 (±0.02) | 0.60 | 1.0000 | |
| - | Veins-Peristome > Peristome-Tube | -0.31 (±0.02) | -13.70 | <0.001 | *** |
| - | Veins-Tube > Peristome-Tube | -0.33 (±0.02) | -14.27 | <0.001 | *** |
| - | summer > autumn | -0.8 (±0.11) | -7.51 | <0.001 | *** |
| Areoles-Veins | *S.* x *leucophylla* > *S.* x Juthatip soper | 0.43 (±0.1) | 4.19 | 0.0039 | ** |
| Areoles-Veins | *S.* x *leucophylla* > *S.* x *mitchelliana* | 0.64 (±0.1) | 6.21 | <0.001 | *** |
| Areoles-Veins | *S.* x *leucophylla* > *S. purpurea* | 0.91 (±0.1) | 8.83 | <0.001 | *** |
| Areoles-Veins | *S.* x Juthatip soper > *S.* x *mitchelliana* | 0.21 (±0.1) | 2.03 | 0.3569 | |
| Areoles-Veins | *S.* x Juthatip soper > *S. purpurea* | 0.48 (±0.1) | 4.65 | 0.0015 | ** |
| Areoles-Veins | *S.* x *mitchelliana* > *S. purpurea* | 0.27 (±0.1) | 2.62 | 0.1099 | |
| Areoles-Peristome | *S.* x *leucophylla* > *S.* x Juthatip soper | 0.72 (±0.11) | 6.69 | <0.001 | *** |
| Areoles-Peristome | *S.* x *leucophylla* > *S.* x *mitchelliana* | 1.18 (±0.11) | 10.98 | <0.001 | *** |
| Areoles-Peristome | *S.* x *leucophylla* > *S. purpurea* | 1.09 (±0.11) | 10.16 | <0.001 | *** |
| Areoles-Peristome | *S.* x Juthatip soper > *S.* x *mitchelliana* | 0.46 (±0.11) | 4.32 | 0.0022 | ** |
| Areoles-Peristome | *S.* x Juthatip soper > *S. purpurea* | 0.37 (±0.11) | 3.48 | 0.0156 | * |
| Areoles-Peristome | *S.* x *mitchelliana* > *S. purpurea* | -0.09 (±0.11) | -0.86 | 1.0000 | |
| Areoles-Tube | *S.* x *leucophylla* > *S.* x Juthatip soper | 0.37 (±0.11) | 3.46 | 0.0159 | * |
| Areoles-Tube | *S.* x *leucophylla* > *S.* x *mitchelliana* | 1.08 (±0.11) | 10.12 | <0.001 | *** |
| Areoles-Tube | *S.* x *leucophylla* > *S. purpurea* | 1.19 (±0.11) | 11.16 | <0.001 | *** |
| Areoles-Tube | *S.* x Juthatip soper > *S.* x *mitchelliana* | 0.71 (±0.11) | 6.66 | <0.001 | *** |
| Areoles-Tube | *S.* x Juthatip soper > *S. purpurea* | 0.82 (±0.11) | 7.70 | <0.001 | *** |
| Areoles-Tube | *S.* x *mitchelliana* > *S. purpurea* | 0.11 (±0.11) | 1.03 | 1.0000 | |
| Veins-Peristome | *S.* x *leucophylla* > *S.* x Juthatip soper | 0.16 (±0.11) | 1.48 | 0.9419 | |
| Veins-Peristome | *S.* x *leucophylla* > *S.* x *mitchelliana* | 0.42 (±0.11) | 3.94 | 0.0053 | ** |
| Veins-Peristome | *S.* x *leucophylla* > *S. purpurea* | 0.35 (±0.11) | 3.26 | 0.0250 | * |
| Veins-Peristome | *S.* x Juthatip soper > *S.* x *mitchelliana* | 0.26 (±0.11) | 2.47 | 0.1388 | |
| Veins-Peristome | *S.* x Juthatip soper > *S. purpurea* | 0.19 (±0.11) | 1.79 | 0.5366 | |
| Veins-Peristome | *S.* x *mitchelliana* > *S. purpurea* | -0.07 (±0.11) | -0.69 | 1.0000 | |
| Veins-Tube | *S.* x *leucophylla* > *S.* x Juthatip soper | -0.06 (±0.11) | -0.53 | 1.0000 | |
| Veins-Tube | *S.* x *leucophylla* > *S.* x *mitchelliana* | -0.58 (±0.11) | -5.45 | <0.001 | *** |
| Veins-Tube | *S.* x *leucophylla* > *S. purpurea* | 0.08 (±0.11) | 0.73 | 1.0000 | |
| Veins-Tube | *S.* x Juthatip soper > *S.* x *mitchelliana* | -0.52 (±0.11) | -4.92 | <0.001 | *** |
| Veins-Tube | *S.* x Juthatip soper > *S. purpurea* | 0.13 (±0.11) | 1.26 | 1.0000 | |
| Veins-Tube | *S.* x *mitchelliana* > *S. purpurea* | 0.66 (±0.11) | 6.18 | <0.001 | *** |
| Peristome-Tube | *S.* x *leucophylla* > *S.* x Juthatip soper | 0.27 (±0.11) | 2.38 | 0.1558 | |
| Peristome-Tube | *S.* x *leucophylla* > *S.* x *mitchelliana* | 0.11 (±0.11) | 0.95 | 1.0000 | |
| Peristome-Tube | *S.* x *leucophylla* > *S. purpurea* | 0.24 (±0.11) | 2.13 | 0.2640 | |
| Peristome-Tube | *S.* x Juthatip soper > *S.* x *mitchelliana* | -0.16 (±0.11) | -1.41 | 1.0000 | |
| Peristome-Tube | *S.* x Juthatip soper > *S. purpurea* | -0.03 (±0.11) | -0.25 | 1.0000 | |
| Peristome-Tube | *S.* x *mitchelliana* > *S. purpurea* | 0.13 (±0.11) | 1.17 | 1.0000 | |
| *S.* x *leucophylla* | Areoles-Veins > Areoles-Peristome | -0.47 (±0.03) | -13.78 | <0.001 | *** |
| *S.* x *leucophylla* | Areoles-Veins > Areoles-Tube | 0 (±0.03) | 0.06 | 1.0000 | |
| *S.* x *leucophylla* | Areoles-Veins > Veins-Peristome | 0.61 (±0.03) | 18.15 | <0.001 | *** |
| *S.* x *leucophylla* | Areoles-Veins > Veins-Tube | 0.99 (±0.03) | 30.16 | <0.001 | *** |
| *S.* x *leucophylla* | Areoles-Veins > Peristome-Tube | 0.37 (±0.04) | 9.02 | <0.001 | *** |

| | | | | | |
|---|---|---|---|---|---|
| *S.* x *leucophylla* | Areoles-Peristome > Areoles-Tube | 0.47 (±0.04) | 12.27 | <0.001 | *** |
| *S.* x *leucophylla* | Areoles-Peristome > Veins-Peristome | 1.08 (±0.04) | 27.93 | <0.001 | *** |
| *S.* x *leucophylla* | Areoles-Peristome > Veins-Tube | 1.46 (±0.04) | 38.43 | <0.001 | *** |
| *S.* x *leucophylla* | Areoles-Peristome > Peristome-Tube | 0.84 (±0.05) | 18.50 | <0.001 | *** |
| *S.* x *leucophylla* | Areoles-Tube > Veins-Peristome | 0.61 (±0.04) | 16.01 | <0.001 | *** |
| *S.* x *leucophylla* | Areoles-Tube > Veins-Tube | 0.99 (±0.04) | 26.51 | <0.001 | *** |
| *S.* x *leucophylla* | Areoles-Tube > Peristome-Tube | 0.37 (±0.04) | 8.25 | <0.001 | *** |
| *S.* x *leucophylla* | Veins-Peristome > Veins-Tube | 0.38 (±0.04) | 10.18 | <0.001 | *** |
| *S.* x *leucophylla* | Veins-Peristome > Peristome-Tube | -0.24 (±0.05) | -5.25 | <0.001 | *** |
| *S.* x *leucophylla* | Veins-Tube > Peristome-Tube | -0.62 (±0.04) | -13.88 | <0.001 | *** |
| *S.* x Juthatip soper | Areoles-Veins > Areoles-Peristome | -0.19 (±0.03) | -5.81 | <0.001 | *** |
| *S.* x Juthatip soper | Areoles-Veins > Areoles-Tube | -0.06 (±0.03) | -1.95 | 0.7761 | |
| *S.* x Juthatip soper | Areoles-Veins > Veins-Peristome | 0.33 (±0.03) | 10.48 | <0.001 | *** |
| *S.* x Juthatip soper | Areoles-Veins > Veins-Tube | 0.5 (±0.03) | 15.77 | <0.001 | *** |
| *S.* x Juthatip soper | Areoles-Veins > Peristome-Tube | 0.21 (±0.04) | 4.97 | <0.001 | *** |
| *S.* x Juthatip soper | Areoles-Peristome > Areoles-Tube | 0.12 (±0.04) | 3.32 | 0.0135 | * |
| *S.* x Juthatip soper | Areoles-Peristome > Veins-Peristome | 0.52 (±0.04) | 14.06 | <0.001 | *** |
| *S.* x Juthatip soper | Areoles-Peristome > Veins-Tube | 0.69 (±0.04) | 18.61 | <0.001 | *** |
| *S.* x Juthatip soper | Areoles-Peristome > Peristome-Tube | 0.39 (±0.05) | 8.61 | <0.001 | *** |
| *S.* x Juthatip soper | Areoles-Tube > Veins-Peristome | 0.39 (±0.04) | 10.68 | <0.001 | *** |
| *S.* x Juthatip soper | Areoles-Tube > Veins-Tube | 0.56 (±0.04) | 15.23 | <0.001 | *** |
| *S.* x Juthatip soper | Areoles-Tube > Peristome-Tube | 0.27 (±0.05) | 5.89 | <0.001 | *** |
| *S.* x Juthatip soper | Veins-Peristome > Veins-Tube | 0.17 (±0.04) | 4.62 | <0.001 | *** |
| *S.* x Juthatip soper | Veins-Peristome > Peristome-Tube | -0.13 (±0.05) | -2.78 | 0.0806 | |
| *S.* x Juthatip soper | Veins-Tube > Peristome-Tube | -0.29 (±0.05) | -6.51 | <0.001 | *** |
| *S.* x *mitchelliana* | Areoles-Veins > Areoles-Peristome | 0.07 (±0.03) | 2.02 | 0.6497 | |
| *S.* x *mitchelliana* | Areoles-Veins > Areoles-Tube | 0.44 (±0.03) | 13.61 | <0.001 | *** |
| *S.* x *mitchelliana* | Areoles-Veins > Veins-Peristome | 0.39 (±0.03) | 11.62 | <0.001 | *** |
| *S.* x *mitchelliana* | Areoles-Veins > Veins-Tube | -0.23 (±0.03) | -7.22 | <0.001 | *** |
| *S.* x *mitchelliana* | Areoles-Veins > Peristome-Tube | -0.16 (±0.04) | -3.79 | 0.0022 | ** |
| *S.* x *mitchelliana* | Areoles-Peristome > Areoles-Tube | 0.37 (±0.04) | 9.75 | <0.001 | *** |
| *S.* x *mitchelliana* | Areoles-Peristome > Veins-Peristome | 0.32 (±0.04) | 8.20 | <0.001 | *** |
| *S.* x *mitchelliana* | Areoles-Peristome > Veins-Tube | -0.3 (±0.04) | -7.87 | <0.001 | *** |
| *S.* x *mitchelliana* | Areoles-Peristome > Peristome-Tube | -0.23 (±0.05) | -4.84 | <0.001 | *** |
| *S.* x *mitchelliana* | Areoles-Tube > Veins-Peristome | -0.05 (±0.04) | -1.36 | 1.0000 | |
| *S.* x *mitchelliana* | Areoles-Tube > Veins-Tube | -0.67 (±0.04) | -18.02 | <0.001 | *** |
| *S.* x *mitchelliana* | Areoles-Tube > Peristome-Tube | -0.6 (±0.05) | -12.80 | <0.001 | *** |
| *S.* x *mitchelliana* | Veins-Peristome > Veins-Tube | -0.62 (±0.04) | -16.24 | <0.001 | *** |
| *S.* x *mitchelliana* | Veins-Peristome > Peristome-Tube | -0.55 (±0.05) | -11.53 | <0.001 | *** |
| *S.* x *mitchelliana* | Veins-Tube > Peristome-Tube | 0.07 (±0.05) | 1.47 | 1.0000 | |
| S. *purpurea* | Areoles-Veins > Areoles-Peristome | -0.3 (±0.03) | -9.36 | <0.001 | *** |
| S. *purpurea* | Areoles-Veins > Areoles-Tube | 0.28 (±0.03) | 8.81 | <0.001 | *** |
| S. *purpurea* | Areoles-Veins > Veins-Peristome | 0.04 (±0.03) | 1.35 | 1.0000 | |
| S. *purpurea* | Areoles-Veins > Veins-Tube | 0.15 (±0.03) | 4.91 | <0.001 | *** |
| S. *purpurea* | Areoles-Veins > Peristome-Tube | -0.3 (±0.04) | -7.29 | <0.001 | *** |

| | | | | | |
|---|---|---|---|---|---|
| S. *purpurea* | Areoles-Peristome > Areoles-Tube | 0.57 (±0.04) | 15.64 | <0.001 | *** |
| S. *purpurea* | Areoles-Peristome > Veins-Peristome | 0.34 (±0.04) | 9.23 | <0.001 | *** |
| S. *purpurea* | Areoles-Peristome > Veins-Tube | 0.45 (±0.04) | 12.29 | <0.001 | *** |
| S. *purpurea* | Areoles-Peristome > Peristome-Tube | -0.01 (±0.05) | -0.14 | 1.0000 | |
| S. *purpurea* | Areoles-Tube > Veins-Peristome | -0.23 (±0.04) | -6.42 | <0.001 | *** |
| S. *purpurea* | Areoles-Tube > Veins-Tube | -0.12 (±0.04) | -3.35 | 0.0121 | * |
| S. *purpurea* | Areoles-Tube > Peristome-Tube | -0.58 (±0.05) | -12.77 | <0.001 | *** |
| S. *purpurea* | Veins-Peristome > Veins-Tube | 0.11 (±0.04) | 3.06 | 0.0329 | * |
| S. *purpurea* | Veins-Peristome > Peristome-Tube | -0.34 (±0.05) | -7.59 | <0.001 | *** |
| S. *purpurea* | Veins-Tube > Peristome-Tube | -0.46 (±0.05) | -10.06 | <0.001 | *** |
| Stage 1 | Areoles-Veins > Areoles-Peristome | -0.15 (±0.04) | -3.70 | 0.0032 | ** |
| Stage 1 | Areoles-Veins > Areoles-Tube | 0.03 (±0.04) | 0.86 | 1.0000 | |
| Stage 1 | Areoles-Veins > Veins-Peristome | 0.27 (±0.04) | 6.97 | <0.001 | *** |
| Stage 1 | Areoles-Veins > Veins-Tube | 0.28 (±0.04) | 7.32 | <0.001 | *** |
| Stage 1 | Areoles-Veins > Peristome-Tube | -0.01 (±0.05) | -0.14 | 1.0000 | |
| Stage 1 | Areoles-Peristome > Areoles-Tube | 0.18 (±0.05) | 3.97 | 0.0011 | ** |
| Stage 1 | Areoles-Peristome > Veins-Peristome | 0.42 (±0.05) | 9.22 | <0.001 | *** |
| Stage 1 | Areoles-Peristome > Veins-Tube | 0.43 (±0.05) | 9.53 | <0.001 | *** |
| Stage 1 | Areoles-Peristome > Peristome-Tube | 0.14 (±0.06) | 2.54 | 0.1665 | |
| Stage 1 | Areoles-Tube > Veins-Peristome | 0.24 (±0.05) | 5.31 | <0.001 | *** |
| Stage 1 | Areoles-Tube > Veins-Tube | 0.25 (±0.04) | 5.58 | <0.001 | *** |
| Stage 1 | Areoles-Tube > Peristome-Tube | -0.04 (±0.05) | -0.75 | 1.0000 | |
| Stage 1 | Veins-Peristome > Veins-Tube | 0.01 (±0.04) | 0.22 | 1.0000 | |
| Stage 1 | Veins-Peristome > Peristome-Tube | -0.28 (±0.05) | -5.11 | <0.001 | *** |
| Stage 1 | Veins-Tube > Peristome-Tube | -0.29 (±0.05) | -5.33 | <0.001 | *** |
| Stage 2 | Areoles-Veins > Areoles-Peristome | -0.37 (±0.02) | -16.07 | <0.001 | *** |
| Stage 2 | Areoles-Veins > Areoles-Tube | 0.23 (±0.02) | 10.03 | <0.001 | *** |
| Stage 2 | Areoles-Veins > Veins-Peristome | 0.33 (±0.02) | 14.36 | <0.001 | *** |
| Stage 2 | Areoles-Veins > Veins-Tube | 0.73 (±0.02) | 32.15 | <0.001 | *** |
| Stage 2 | Areoles-Veins > Peristome-Tube | 0.25 (±0.03) | 8.51 | <0.001 | *** |
| Stage 2 | Areoles-Peristome > Areoles-Tube | 0.6 (±0.03) | 22.61 | <0.001 | *** |
| Stage 2 | Areoles-Peristome > Veins-Peristome | 0.69 (±0.03) | 26.35 | <0.001 | *** |
| Stage 2 | Areoles-Peristome > Veins-Tube | 1.1 (±0.03) | 41.76 | <0.001 | *** |
| Stage 2 | Areoles-Peristome > Peristome-Tube | 0.62 (±0.03) | 19.14 | <0.001 | *** |
| Stage 2 | Areoles-Tube > Veins-Peristome | 0.1 (±0.03) | 3.74 | 0.0027 | ** |
| Stage 2 | Areoles-Tube > Veins-Tube | 0.5 (±0.03) | 19.15 | <0.001 | *** |
| Stage 2 | Areoles-Tube > Peristome-Tube | 0.02 (±0.03) | 0.68 | 1.0000 | |
| Stage 2 | Veins-Peristome > Veins-Tube | 0.41 (±0.03) | 15.41 | <0.001 | *** |
| Stage 2 | Veins-Peristome > Peristome-Tube | -0.08 (±0.03) | -2.38 | 0.2608 | |
| Stage 2 | Veins-Tube > Peristome-Tube | -0.48 (±0.03) | -14.96 | <0.001 | *** |
| Stage 3 | Areoles-Veins > Areoles-Peristome | -0.14 (±0.02) | -6.25 | <0.001 | *** |
| Stage 3 | Areoles-Veins > Areoles-Tube | 0.24 (±0.02) | 10.41 | <0.001 | *** |
| Stage 3 | Areoles-Veins > Veins-Peristome | 0.46 (±0.02) | 20.13 | <0.001 | *** |
| Stage 3 | Areoles-Veins > Veins-Tube | 0.26 (±0.02) | 11.36 | <0.001 | *** |
| Stage 3 | Areoles-Veins > Peristome-Tube | 0.04 (±0.03) | 1.33 | 1.0000 | |

| | | | | | |
|---|---|---|---|---|---|
| Stage 3 | Areoles-Peristome > Areoles-Tube | 0.38 (±0.03) | 14.48 | <0.001 | *** |
| Stage 3 | Areoles-Peristome > Veins-Peristome | 0.6 (±0.03) | 22.94 | <0.001 | *** |
| Stage 3 | Areoles-Peristome > Veins-Tube | 0.4 (±0.03) | 15.29 | <0.001 | *** |
| Stage 3 | Areoles-Peristome > Peristome-Tube | 0.18 (±0.03) | 5.65 | <0.001 | *** |
| Stage 3 | Areoles-Tube > Veins-Peristome | 0.22 (±0.03) | 8.40 | <0.001 | *** |
| Stage 3 | Areoles-Tube > Veins-Tube | 0.02 (±0.03) | 0.84 | 1.0000 | |
| Stage 3 | Areoles-Tube > Peristome-Tube | -0.2 (±0.03) | -6.20 | <0.001 | *** |
| Stage 3 | Veins-Peristome > Veins-Tube | -0.2 (±0.03) | -7.54 | <0.001 | *** |
| Stage 3 | Veins-Peristome > Peristome-Tube | -0.42 (±0.03) | -13.10 | <0.001 | *** |
| Stage 3 | Veins-Tube > Peristome-Tube | -0.22 (±0.03) | -6.88 | <0.001 | *** |
| Stage 4 | Areoles-Veins > Areoles-Peristome | -0.22 (±0.04) | -5.55 | <0.001 | *** |
| Stage 4 | Areoles-Veins > Areoles-Tube | 0.15 (±0.04) | 3.89 | 0.0015 | ** |
| Stage 4 | Areoles-Veins > Veins-Peristome | 0.31 (±0.04) | 7.61 | <0.001 | *** |
| Stage 4 | Areoles-Veins > Veins-Tube | 0.14 (±0.04) | 3.41 | 0.0098 | ** |
| Stage 4 | Areoles-Veins > Peristome-Tube | -0.17 (±0.05) | -3.27 | 0.0164 | * |
| Stage 4 | Areoles-Peristome > Areoles-Tube | 0.38 (±0.05) | 8.16 | <0.001 | *** |
| Stage 4 | Areoles-Peristome > Veins-Peristome | 0.53 (±0.05) | 11.30 | <0.001 | *** |
| Stage 4 | Areoles-Peristome > Veins-Tube | 0.36 (±0.05) | 7.75 | <0.001 | *** |
| Stage 4 | Areoles-Peristome > Peristome-Tube | 0.05 (±0.06) | 0.93 | 1.0000 | |
| Stage 4 | Areoles-Tube > Veins-Peristome | 0.15 (±0.05) | 3.31 | 0.0143 | * |
| Stage 4 | Areoles-Tube > Veins-Tube | -0.02 (±0.05) | -0.42 | 1.0000 | |
| Stage 4 | Areoles-Tube > Peristome-Tube | -0.33 (±0.06) | -5.70 | <0.001 | *** |
| Stage 4 | Veins-Peristome > Veins-Tube | -0.17 (±0.05) | -3.72 | 0.0030 | ** |
| Stage 4 | Veins-Peristome > Peristome-Tube | -0.48 (±0.06) | -8.31 | <0.001 | *** |
| Stage 4 | Veins-Tube > Peristome-Tube | -0.31 (±0.06) | -5.36 | <0.001 | *** |
| Stage 1 | *S.* x *leucophylla* > *S.* x Juthatip soper | -0.14 (±0.25) | -0.57 | 1.0000 | |
| Stage 1 | *S.* x *leucophylla* > *S.* x *mitchelliana* | 0.19 (±0.25) | 0.76 | 1.0000 | |
| Stage 1 | *S.* x *leucophylla* > *S. purpurea* | 0.34 (±0.25) | 1.36 | 1.0000 | |
| Stage 1 | *S.* x Juthatip soper > *S.* x *mitchelliana* | 0.33 (±0.25) | 1.33 | 1.0000 | |
| Stage 1 | *S.* x Juthatip soper > *S. purpurea* | 0.48 (±0.25) | 1.93 | 0.4394 | |
| Stage 1 | *S.* x *mitchelliana* > *S. purpurea* | 0.15 (±0.25) | 0.60 | 1.0000 | |
| Stage 2 | *S.* x *leucophylla* > *S.* x Juthatip soper | 0.21 (±0.14) | 1.49 | 0.9426 | |
| Stage 2 | *S.* x *leucophylla* > *S.* x *mitchelliana* | 0.67 (±0.14) | 4.71 | 0.0017 | ** |
| Stage 2 | *S.* x *leucophylla* > *S. purpurea* | 0.51 (±0.14) | 3.53 | 0.0179 | * |
| Stage 2 | *S.* x Juthatip soper > *S.* x *mitchelliana* | 0.46 (±0.14) | 3.22 | 0.0342 | * |
| Stage 2 | *S.* x Juthatip soper > *S. purpurea* | 0.29 (±0.14) | 2.05 | 0.3519 | |
| Stage 2 | *S.* x *mitchelliana* > *S. purpurea* | -0.17 (±0.14) | -1.17 | 1.0000 | |
| Stage 3 | *S.* x *leucophylla* > *S.* x Juthatip soper | 0.28 (±0.14) | 1.96 | 0.4101 | |
| Stage 3 | *S.* x *leucophylla* > *S.* x *mitchelliana* | 0.31 (±0.14) | 2.18 | 0.2753 | |
| Stage 3 | *S.* x *leucophylla* > *S. purpurea* | 0.79 (±0.14) | 5.56 | <0.001 | *** |
| Stage 3 | *S.* x Juthatip soper > *S.* x *mitchelliana* | 0.03 (±0.14) | 0.21 | 1.0000 | |
| Stage 3 | *S.* x Juthatip soper > *S. purpurea* | 0.51 (±0.14) | 3.60 | 0.0158 | * |
| Stage 3 | *S.* x *mitchelliana* > *S. purpurea* | 0.48 (±0.14) | 3.38 | 0.0244 | * |
| Stage 4 | *S.* x *leucophylla* > *S.* x Juthatip soper | 0.91 (±0.25) | 3.65 | 0.0139 | * |
| Stage 4 | *S.* x *leucophylla* > *S.* x *mitchelliana* | 0.73 (±0.25) | 2.93 | 0.0619 | |

| | | | | | |
|---|---|---|---|---|---|
| Stage 4 | *S.* x *leucophylla* > *S. purpurea* | 0.93 (±0.25) | 3.76 | 0.0111 | * |
| Stage 4 | *S.* x Juthatip soper > *S.* x *mitchelliana* | -0.18 (±0.25) | -0.73 | 1.0000 | |
| Stage 4 | *S.* x Juthatip soper > *S. purpurea* | 0.03 (±0.25) | 0.11 | 1.0000 | |
| Stage 4 | *S.* x *mitchelliana* > *S. purpurea* | 0.21 (±0.25) | 0.83 | 1.0000 | |
| Areoles-Veins | Stage 1 > (2, 3, 4) | -0.07 (±0.1) | -0.68 | 1.0000 | |
| Areoles-Veins | Stage 2 > (1, 3, 4) | 0.16 (±0.1) | 1.64 | 0.4805 | |
| Areoles-Veins | Stage 3 > (1, 2, 4) | 0.07 (±0.07) | 0.94 | 1.0000 | |
| Areoles-Veins | Stage 4 > (1, 2, 3) | -0.16 (±0.1) | -1.63 | 0.4902 | |
| Areoles-Peristome | Stage 1 > (2, 3, 4) | -0.16 (±0.1) | -1.59 | 0.5123 | |
| Areoles-Peristome | Stage 2 > (1, 3, 4) | 0.36 (±0.1) | 3.56 | 0.0099 | ** |
| Areoles-Peristome | Stage 3 > (1, 2, 4) | -0.04 (±0.07) | -0.48 | 1.0000 | |
| Areoles-Peristome | Stage 4 > (1, 2, 3) | -0.16 (±0.1) | -1.52 | 0.5816 | |
| Areoles-Tube | Stage 1 > (2, 3, 4) | 0.11 (±0.1) | 1.02 | 1.0000 | |
| Areoles-Tube | Stage 2 > (1, 3, 4) | 0.08 (±0.1) | 0.76 | 1.0000 | |
| Areoles-Tube | Stage 3 > (1, 2, 4) | -0.03 (±0.07) | -0.41 | 1.0000 | |
| Areoles-Tube | Stage 4 > (1, 2, 3) | -0.15 (±0.1) | -1.46 | 0.6447 | |
| Veins-Peristome | Stage 1 > (2, 3, 4) | 0.02 (±0.1) | 0.23 | 1.0000 | |
| Veins-Peristome | Stage 2 > (1, 3, 4) | 0.18 (±0.1) | 1.81 | 0.3532 | |
| Veins-Peristome | Stage 3 > (1, 2, 4) | -0.09 (±0.07) | -1.17 | 1.0000 | |
| Veins-Peristome | Stage 4 > (1, 2, 3) | -0.12 (±0.1) | -1.14 | 1.0000 | |
| Veins-Tube | Stage 1 > (2, 3, 4) | 0.03 (±0.1) | 0.24 | 1.0000 | |
| Veins-Tube | Stage 2 > (1, 3, 4) | -0.34 (±0.1) | -3.42 | 0.0134 | * |
| Veins-Tube | Stage 3 > (1, 2, 4) | 0.19 (±0.07) | 2.56 | 0.0776 | |
| Veins-Tube | Stage 4 > (1, 2, 3) | 0.13 (±0.1) | 1.23 | 0.9329 | |
| Peristome-Tube | Stage 1 > (2, 3, 4) | -0.02 (±0.11) | -0.20 | 1.0000 | |
| Peristome-Tube | Stage 2 > (1, 3, 4) | -0.13 (±0.1) | -1.29 | 0.8463 | |
| Peristome-Tube | Stage 3 > (1, 2, 4) | 0.05 (±0.08) | 0.68 | 1.0000 | |
| Peristome-Tube | Stage 4 > (1, 2, 3) | 0.1 (±0.11) | 0.93 | 1.0000 | |
| *S.* x *leucophylla* | Stage 1 > (2, 3, 4) | -0.37 (±0.19) | -1.91 | 0.3034 | |
| *S.* x *leucophylla* | Stage 2 > (1, 3, 4) | 0.04 (±0.15) | 0.24 | 1.0000 | |
| *S.* x *leucophylla* | Stage 3 > (1, 2, 4) | 0.01 (±0.14) | 0.09 | 1.0000 | |
| *S.* x *leucophylla* | Stage 4 > (1, 2, 3) | 0.32 (±0.19) | 1.65 | 0.4772 | |
| *S.* x Juthatip soper | Stage 1 > (2, 3, 4) | 0.24 (±0.19) | 1.26 | 0.9133 | |
| *S.* x Juthatip soper | Stage 2 > (1, 3, 4) | 0.17 (±0.15) | 1.13 | 1.0000 | |
| *S.* x Juthatip soper | Stage 3 > (1, 2, 4) | 0.06 (±0.14) | 0.42 | 1.0000 | |
| *S.* x Juthatip soper | Stage 4 > (1, 2, 3) | -0.47 (±0.19) | -2.45 | 0.1080 | |
| *S.* x *mitchelliana* | Stage 1 > (2, 3, 4) | 0.02 (±0.19) | 0.08 | 1.0000 | |
| *S.* x *mitchelliana* | Stage 2 > (1, 3, 4) | -0.23 (±0.15) | -1.49 | 0.6249 | |
| *S.* x *mitchelliana* | Stage 3 > (1, 2, 4) | 0.23 (±0.14) | 1.68 | 0.4565 | |
| *S.* x *mitchelliana* | Stage 4 > (1, 2, 3) | -0.02 (±0.19) | -0.09 | 1.0000 | |
| S. *purpurea* | Stage 1 > (2, 3, 4) | 0.04 (±0.19) | 0.22 | 1.0000 | |
| S. *purpurea* | Stage 2 > (1, 3, 4) | 0.22 (±0.15) | 1.44 | 0.6844 | |
| S. *purpurea* | Stage 3 > (1, 2, 4) | -0.19 (±0.14) | -1.40 | 0.7251 | |
| S. *purpurea* | Stage 4 > (1, 2, 3) | -0.07 (±0.19) | -0.36 | 1.0000 | |

**Table S4.** Results of the post-hoc tests associated to the best linear mixed model, presented in Table 2, to explain the variation in brightness contrast between any two pitcher areas. The variable brightness contrast was square-root transformed to match residual normality. P-values were adjusted for multiple comparisons with the Bonferroni correction. Symbols describe various levels of p-values *: p<0.05, **: p<0.01, and ***: p<0.001. Within a level of factor 1, we tested the oriented contrast between two levels of factor 2 or between 2 grouped levels of factor 2. For instance, for the contrast produced by areoles against veins, we tested whether the brightness contrast of *S.* x *leucophylla* was higher than the brightness contrast of *S.* x Juthatip soper and it was significantly different (p-value=0.0136). Symbols "-" mean that the oriented contrasts were tested overall and not within a level of another factor. For instance, we tested whether the brightness contrast of *S.* x *leucophylla* was higher than the brightness contrast of *S.* x Juthatip soper overall.

| Level on factor 1 | Contrast on factor 2 | Coefficient (± standard error) | t-ratio | adjusted p-value | |
|---|---|---|---|---|---|
| - | *S.* x *leucophylla* > *S.* x Juthatip soper | 0.52 (±0.16) | 3.20 | 0.0357 | * |
| - | *S.* x *leucophylla* > *S.* x *mitchelliana* | 0.45 (±0.16) | 2.81 | 0.0790 | |
| - | *S.* x *leucophylla* > *S. purpurea* | 0.55 (±0.16) | 3.40 | 0.0238 | * |
| - | *S.* x Juthatip soper > *S.* x *mitchelliana* | -0.06 (±0.16) | -0.39 | 1.0000 | |
| - | *S.* x Juthatip soper > *S. purpurea* | 0.03 (±0.16) | 0.20 | 1.0000 | |
| - | *S.* x *mitchelliana* > *S. purpurea* | 0.09 (±0.16) | 0.59 | 1.0000 | |
| - | Stage 1 > Stage 2 | -0.29 (±0.2) | -1.47 | 0.9718 | |
| - | Stage 1 > Stage 3 | -0.49 (±0.16) | -3.04 | 0.0489 | * |
| - | Stage 1 > Stage 4 | -0.47 (±0.2) | -2.39 | 0.1803 | |
| - | Stage 2 > Stage 3 | -0.2 (±0.16) | -1.25 | 1.0000 | |
| - | Stage 2 > Stage 4 | -0.18 (±0.2) | -0.93 | 1.0000 | |
| - | Stage 3 > Stage 4 | 0.02 (±0.16) | 0.11 | 1.0000 | |
| - | Areoles-Veins > Areoles-Peristome | -0.58 (±0.03) | -21.31 | <0.001 | *** |
| - | Areoles-Veins > Areoles-Tube | 0.48 (±0.03) | 17.87 | <0.001 | *** |
| - | Areoles-Veins > Veins-Peristome | 0.08 (±0.03) | 3.02 | 0.0386 | * |
| - | Areoles-Veins > Veins-Tube | 0.39 (±0.03) | 14.56 | <0.001 | *** |
| - | Areoles-Veins > Peristome-Tube | -0.27 (±0.04) | -7.73 | <0.001 | *** |
| - | Areoles-Peristome > Areoles-Tube | 1.07 (±0.03) | 33.91 | <0.001 | *** |
| - | Areoles-Peristome > Veins-Peristome | 0.67 (±0.03) | 21.06 | <0.001 | *** |
| - | Areoles-Peristome > Veins-Tube | 0.98 (±0.03) | 31.13 | <0.001 | *** |
| - | Areoles-Peristome > Peristome-Tube | 0.31 (±0.04) | 8.12 | <0.001 | *** |
| - | Areoles-Tube > Veins-Peristome | -0.4 (±0.03) | -12.81 | <0.001 | *** |
| - | Areoles-Tube > Veins-Tube | -0.09 (±0.03) | -2.95 | 0.0477 | * |
| - | Areoles-Tube > Peristome-Tube | -0.75 (±0.04) | -19.68 | <0.001 | *** |

| | | | | | |
|---|---|---|---|---|---|
| - | Veins-Peristome > Veins-Tube | 0.31 (±0.03) | 9.92 | <0.001 | *** |
| - | Veins-Peristome > Peristome-Tube | -0.35 (±0.04) | -9.18 | <0.001 | *** |
| - | Veins-Tube > Peristome-Tube | -0.66 (±0.04) | -17.34 | <0.001 | *** |
| - | summer > autumn | -0.45 (±0.17) | -2.65 | 0.0181 | * |
| Areoles-Veins | *S.* x *leucophylla* > *S.* x Juthatip soper | 0.59 (±0.17) | 3.60 | 0.0136 | * |
| Areoles-Veins | *S.* x *leucophylla* > *S.* x *mitchelliana* | 0.34 (±0.17) | 2.03 | 0.3510 | |
| Areoles-Veins | *S.* x *leucophylla* > *S. purpurea* | 0.54 (±0.17) | 3.25 | 0.0290 | * |
| Areoles-Veins | *S.* x Juthatip soper > *S.* x *mitchelliana* | -0.26 (±0.16) | -1.57 | 0.8092 | |
| Areoles-Veins | *S.* x Juthatip soper > *S. purpurea* | -0.06 (±0.16) | -0.36 | 1.0000 | |
| Areoles-Veins | *S.* x *mitchelliana* > *S. purpurea* | 0.2 (±0.16) | 1.22 | 1.0000 | |
| Areoles-Peristome | *S.* x *leucophylla* > *S.* x Juthatip soper | 1.07 (±0.17) | 6.24 | <0.001 | *** |
| Areoles-Peristome | *S.* x *leucophylla* > *S.* x *mitchelliana* | 1.36 (±0.17) | 7.91 | <0.001 | *** |
| Areoles-Peristome | *S.* x *leucophylla* > *S. purpurea* | 1.05 (±0.17) | 6.16 | <0.001 | *** |
| Areoles-Peristome | *S.* x Juthatip soper > *S.* x *mitchelliana* | 0.29 (±0.17) | 1.69 | 0.6405 | |
| Areoles-Peristome | *S.* x Juthatip soper > *S. purpurea* | -0.01 (±0.17) | -0.09 | 1.0000 | |
| Areoles-Peristome | *S.* x *mitchelliana* > *S. purpurea* | -0.3 (±0.17) | -1.78 | 0.5443 | |
| Areoles-Tube | *S.* x *leucophylla* > *S.* x Juthatip soper | 0.49 (±0.17) | 2.85 | 0.0614 | |
| Areoles-Tube | *S.* x *leucophylla* > *S.* x *mitchelliana* | 0.78 (±0.17) | 4.57 | 0.0012 | ** |
| Areoles-Tube | *S.* x *leucophylla* > *S. purpurea* | 0.74 (±0.17) | 4.32 | 0.0022 | ** |
| Areoles-Tube | *S.* x Juthatip soper > *S.* x *mitchelliana* | 0.29 (±0.17) | 1.72 | 0.6104 | |
| Areoles-Tube | *S.* x Juthatip soper > *S. purpurea* | 0.25 (±0.17) | 1.47 | 0.9513 | |
| Areoles-Tube | *S.* x *mitchelliana* > *S. purpurea* | -0.04 (±0.17) | -0.25 | 1.0000 | |
| Veins-Peristome | *S.* x *leucophylla* > *S.* x Juthatip soper | 0.48 (±0.17) | 2.84 | 0.0633 | |
| Veins-Peristome | *S.* x *leucophylla* > *S.* x *mitchelliana* | 0.57 (±0.17) | 3.30 | 0.0219 | * |
| Veins-Peristome | *S.* x *leucophylla* > *S. purpurea* | 0.62 (±0.17) | 3.61 | 0.0111 | * |
| Veins-Peristome | *S.* x Juthatip soper > *S.* x *mitchelliana* | 0.08 (±0.17) | 0.48 | 1.0000 | |
| Veins-Peristome | *S.* x Juthatip soper > *S. purpurea* | 0.13 (±0.17) | 0.78 | 1.0000 | |
| Veins-Peristome | *S.* x *mitchelliana* > *S. purpurea* | 0.05 (±0.17) | 0.29 | 1.0000 | |
| Veins-Tube | *S.* x *leucophylla* > *S.* x Juthatip soper | -0.05 (±0.17) | -0.27 | 1.0000 | |
| Veins-Tube | *S.* x *leucophylla* > *S.* x *mitchelliana* | -0.77 (±0.17) | -4.53 | 0.0014 | ** |
| Veins-Tube | *S.* x *leucophylla* > *S. purpurea* | -0.19 (±0.17) | -1.10 | 1.0000 | |
| Veins-Tube | *S.* x Juthatip soper > *S.* x *mitchelliana* | -0.73 (±0.17) | -4.27 | 0.0025 | ** |
| Veins-Tube | *S.* x Juthatip soper > *S. purpurea* | -0.14 (±0.17) | -0.83 | 1.0000 | |
| Veins-Tube | *S.* x *mitchelliana* > *S. purpurea* | 0.59 (±0.17) | 3.43 | 0.0167 | * |
| Peristome-Tube | *S.* x *leucophylla* > *S.* x Juthatip soper | 0.5 (±0.18) | 2.76 | 0.0645 | |
| Peristome-Tube | *S.* x *leucophylla* > *S.* x *mitchelliana* | 0.45 (±0.18) | 2.45 | 0.1303 | |
| Peristome-Tube | *S.* x *leucophylla* > *S. purpurea* | 0.52 (±0.18) | 2.90 | 0.0475 | * |
| Peristome-Tube | *S.* x Juthatip soper > *S.* x *mitchelliana* | -0.05 (±0.18) | -0.30 | 1.0000 | |
| Peristome-Tube | *S.* x Juthatip soper > *S. purpurea* | 0.02 (±0.18) | 0.13 | 1.0000 | |
| Peristome-Tube | *S.* x *mitchelliana* > *S. purpurea* | 0.08 (±0.18) | 0.43 | 1.0000 | |
| *S.* x *leucophylla* | Areoles-Veins > Areoles-Peristome | -1.09 (±0.06) | -19.12 | <0.001 | *** |
| *S.* x *leucophylla* | Areoles-Veins > Areoles-Tube | 0.35 (±0.06) | 6.27 | <0.001 | *** |
| *S.* x *leucophylla* | Areoles-Veins > Veins-Peristome | 0.03 (±0.06) | 0.57 | 1.0000 | |
| *S.* x *leucophylla* | Areoles-Veins > Veins-Tube | 1.01 (±0.06) | 18.33 | <0.001 | *** |
| *S.* x *leucophylla* | Areoles-Veins > Peristome-Tube | -0.27 (±0.07) | -3.94 | 0.0012 | ** |

| | | | | | |
|---|---|---|---|---|---|
| *S.* x *leucophylla* | Areoles-Peristome > Areoles-Tube | 1.44 (±0.06) | 22.40 | <0.001 | *** |
| *S.* x *leucophylla* | Areoles-Peristome > Veins-Peristome | 1.12 (±0.06) | 17.34 | <0.001 | *** |
| *S.* x *leucophylla* | Areoles-Peristome > Veins-Tube | 2.1 (±0.06) | 32.97 | <0.001 | *** |
| *S.* x *leucophylla* | Areoles-Peristome > Peristome-Tube | 0.82 (±0.08) | 10.72 | <0.001 | *** |
| *S.* x *leucophylla* | Areoles-Tube > Veins-Peristome | -0.32 (±0.06) | -5.00 | <0.001 | *** |
| *S.* x *leucophylla* | Areoles-Tube > Veins-Tube | 0.66 (±0.06) | 10.56 | <0.001 | *** |
| *S.* x *leucophylla* | Areoles-Tube > Peristome-Tube | -0.62 (±0.08) | -8.26 | <0.001 | *** |
| *S.* x *leucophylla* | Veins-Peristome > Veins-Tube | 0.98 (±0.06) | 15.53 | <0.001 | *** |
| *S.* x *leucophylla* | Veins-Peristome > Peristome-Tube | -0.3 (±0.08) | -4.02 | <0.001 | *** |
| *S.* x *leucophylla* | Veins-Tube > Peristome-Tube | -1.28 (±0.07) | -17.15 | <0.001 | *** |
| *S.* x Juthatip soper | Areoles-Veins > Areoles-Peristome | -0.61 (±0.05) | -11.43 | <0.001 | *** |
| *S.* x Juthatip soper | Areoles-Veins > Areoles-Tube | 0.24 (±0.05) | 4.48 | <0.001 | *** |
| *S.* x Juthatip soper | Areoles-Veins > Veins-Peristome | -0.08 (±0.05) | -1.48 | 1.0000 | |
| *S.* x Juthatip soper | Areoles-Veins > Veins-Tube | 0.37 (±0.05) | 6.95 | <0.001 | *** |
| *S.* x Juthatip soper | Areoles-Veins > Peristome-Tube | -0.37 (±0.07) | -5.27 | <0.001 | *** |
| *S.* x Juthatip soper | Areoles-Peristome > Areoles-Tube | 0.86 (±0.06) | 13.70 | <0.001 | *** |
| *S.* x Juthatip soper | Areoles-Peristome > Veins-Peristome | 0.54 (±0.06) | 8.67 | <0.001 | *** |
| *S.* x Juthatip soper | Areoles-Peristome > Veins-Tube | 0.98 (±0.06) | 15.90 | <0.001 | *** |
| *S.* x Juthatip soper | Areoles-Peristome > Peristome-Tube | 0.25 (±0.08) | 3.25 | 0.0177 | * |
| *S.* x Juthatip soper | Areoles-Tube > Veins-Peristome | -0.32 (±0.06) | -5.17 | <0.001 | *** |
| *S.* x Juthatip soper | Areoles-Tube > Veins-Tube | 0.13 (±0.06) | 2.08 | 0.5705 | |
| *S.* x Juthatip soper | Areoles-Tube > Peristome-Tube | -0.61 (±0.08) | -7.95 | <0.001 | *** |
| *S.* x Juthatip soper | Veins-Peristome > Veins-Tube | 0.45 (±0.06) | 7.31 | <0.001 | *** |
| *S.* x Juthatip soper | Veins-Peristome > Peristome-Tube | -0.29 (±0.08) | -3.79 | 0.0023 | ** |
| *S.* x Juthatip soper | Veins-Tube > Peristome-Tube | -0.74 (±0.08) | -9.69 | <0.001 | *** |
| *S.* x *mitchelliana* | Areoles-Veins > Areoles-Peristome | -0.07 (±0.06) | -1.17 | 1.0000 | |
| *S.* x *mitchelliana* | Areoles-Veins > Areoles-Tube | 0.79 (±0.05) | 14.69 | <0.001 | *** |
| *S.* x *mitchelliana* | Areoles-Veins > Veins-Peristome | 0.26 (±0.06) | 4.71 | <0.001 | *** |
| *S.* x *mitchelliana* | Areoles-Veins > Veins-Tube | -0.1 (±0.05) | -1.82 | 1.0000 | |
| *S.* x *mitchelliana* | Areoles-Veins > Peristome-Tube | -0.16 (±0.07) | -2.23 | 0.3884 | |
| *S.* x *mitchelliana* | Areoles-Peristome > Areoles-Tube | 0.86 (±0.06) | 13.45 | <0.001 | *** |
| *S.* x *mitchelliana* | Areoles-Peristome > Veins-Peristome | 0.33 (±0.07) | 5.02 | <0.001 | *** |
| *S.* x *mitchelliana* | Areoles-Peristome > Veins-Tube | -0.03 (±0.06) | -0.52 | 1.0000 | |
| *S.* x *mitchelliana* | Areoles-Peristome > Peristome-Tube | -0.1 (±0.08) | -1.20 | 1.0000 | |
| *S.* x *mitchelliana* | Areoles-Tube > Veins-Peristome | -0.53 (±0.06) | -8.31 | <0.001 | *** |
| *S.* x *mitchelliana* | Areoles-Tube > Veins-Tube | -0.89 (±0.06) | -14.28 | <0.001 | *** |
| *S.* x *mitchelliana* | Areoles-Tube > Peristome-Tube | -0.96 (±0.08) | -12.11 | <0.001 | *** |
| *S.* x *mitchelliana* | Veins-Peristome > Veins-Tube | -0.36 (±0.06) | -5.65 | <0.001 | *** |
| *S.* x *mitchelliana* | Veins-Peristome > Peristome-Tube | -0.42 (±0.08) | -5.30 | <0.001 | *** |
| *S.* x *mitchelliana* | Veins-Tube > Peristome-Tube | -0.06 (±0.08) | -0.80 | 1.0000 | |
| S. *purpurea* | Areoles-Veins > Areoles-Peristome | -0.57 (±0.05) | -10.78 | <0.001 | *** |
| S. *purpurea* | Areoles-Veins > Areoles-Tube | 0.55 (±0.05) | 10.40 | <0.001 | *** |
| S. *purpurea* | Areoles-Veins > Veins-Peristome | 0.11 (±0.05) | 2.13 | 0.4970 | |
| S. *purpurea* | Areoles-Veins > Veins-Tube | 0.29 (±0.05) | 5.42 | <0.001 | *** |
| S. *purpurea* | Areoles-Veins > Peristome-Tube | -0.28 (±0.07) | -4.09 | <0.001 | *** |

| | | | | | |
|---|---|---|---|---|---|
| S. *purpurea* | Areoles-Peristome > Areoles-Tube | 1.12 (±0.06) | 18.23 | <0.001 | *** |
| S. *purpurea* | Areoles-Peristome > Veins-Peristome | 0.68 (±0.06) | 11.11 | <0.001 | *** |
| S. *purpurea* | Areoles-Peristome > Veins-Tube | 0.86 (±0.06) | 13.94 | <0.001 | *** |
| S. *purpurea* | Areoles-Peristome > Peristome-Tube | 0.29 (±0.08) | 3.76 | 0.0025 | ** |
| S. *purpurea* | Areoles-Tube > Veins-Peristome | -0.44 (±0.06) | -7.12 | <0.001 | *** |
| S. *purpurea* | Areoles-Tube > Veins-Tube | -0.26 (±0.06) | -4.29 | <0.001 | *** |
| S. *purpurea* | Areoles-Tube > Peristome-Tube | -0.83 (±0.08) | -10.95 | <0.001 | *** |
| S. *purpurea* | Veins-Peristome > Veins-Tube | 0.17 (±0.06) | 2.83 | 0.0704 | |
| S. *purpurea* | Veins-Peristome > Peristome-Tube | -0.4 (±0.08) | -5.21 | <0.001 | *** |
| S. *purpurea* | Veins-Tube > Peristome-Tube | -0.57 (±0.08) | -7.49 | <0.001 | *** |
| Stage 1 | Areoles-Veins > Areoles-Peristome | -0.64 (±0.07) | -9.59 | <0.001 | *** |
| Stage 1 | Areoles-Veins > Areoles-Tube | 0.26 (±0.07) | 3.93 | 0.0013 | ** |
| Stage 1 | Areoles-Veins > Veins-Peristome | -0.32 (±0.07) | -4.84 | <0.001 | *** |
| Stage 1 | Areoles-Veins > Veins-Tube | 0.39 (±0.06) | 6.03 | <0.001 | *** |
| Stage 1 | Areoles-Veins > Peristome-Tube | -0.25 (±0.08) | -3.00 | 0.0400 | * |
| Stage 1 | Areoles-Peristome > Areoles-Tube | 0.9 (±0.08) | 11.74 | <0.001 | *** |
| Stage 1 | Areoles-Peristome > Veins-Peristome | 0.32 (±0.08) | 4.22 | <0.001 | *** |
| Stage 1 | Areoles-Peristome > Veins-Tube | 1.03 (±0.08) | 13.63 | <0.001 | *** |
| Stage 1 | Areoles-Peristome > Peristome-Tube | 0.39 (±0.09) | 4.19 | <0.001 | *** |
| Stage 1 | Areoles-Tube > Veins-Peristome | -0.58 (±0.08) | -7.62 | <0.001 | *** |
| Stage 1 | Areoles-Tube > Veins-Tube | 0.13 (±0.08) | 1.77 | 1.0000 | |
| Stage 1 | Areoles-Tube > Peristome-Tube | -0.51 (±0.09) | -5.55 | <0.001 | *** |
| Stage 1 | Veins-Peristome > Veins-Tube | 0.71 (±0.07) | 9.49 | <0.001 | *** |
| Stage 1 | Veins-Peristome > Peristome-Tube | 0.07 (±0.09) | 0.71 | 1.0000 | |
| Stage 1 | Veins-Tube > Peristome-Tube | -0.65 (±0.09) | -7.05 | <0.001 | *** |
| Stage 2 | Areoles-Veins > Areoles-Peristome | -0.18 (±0.04) | -4.75 | <0.001 | *** |
| Stage 2 | Areoles-Veins > Areoles-Tube | 1 (±0.04) | 26.01 | <0.001 | *** |
| Stage 2 | Areoles-Veins > Veins-Peristome | 0.56 (±0.04) | 14.63 | <0.001 | *** |
| Stage 2 | Areoles-Veins > Veins-Tube | 0.58 (±0.04) | 15.10 | <0.001 | *** |
| Stage 2 | Areoles-Veins > Peristome-Tube | 0.39 (±0.05) | 7.92 | <0.001 | *** |
| Stage 2 | Areoles-Peristome > Areoles-Tube | 1.18 (±0.04) | 26.64 | <0.001 | *** |
| Stage 2 | Areoles-Peristome > Veins-Peristome | 0.74 (±0.04) | 16.79 | <0.001 | *** |
| Stage 2 | Areoles-Peristome > Veins-Tube | 0.76 (±0.04) | 17.19 | <0.001 | *** |
| Stage 2 | Areoles-Peristome > Peristome-Tube | 0.57 (±0.05) | 10.59 | <0.001 | *** |
| Stage 2 | Areoles-Tube > Veins-Peristome | -0.44 (±0.04) | -9.85 | <0.001 | *** |
| Stage 2 | Areoles-Tube > Veins-Tube | -0.42 (±0.04) | -9.45 | <0.001 | *** |
| Stage 2 | Areoles-Tube > Peristome-Tube | -0.61 (±0.05) | -11.17 | <0.001 | *** |
| Stage 2 | Veins-Peristome > Veins-Tube | 0.02 (±0.04) | 0.40 | 1.0000 | |
| Stage 2 | Veins-Peristome > Peristome-Tube | -0.17 (±0.05) | -3.12 | 0.0270 | * |
| Stage 2 | Veins-Tube > Peristome-Tube | -0.19 (±0.05) | -3.45 | 0.0084 | ** |
| Stage 3 | Areoles-Veins > Areoles-Peristome | -0.92 (±0.04) | -24.21 | <0.001 | *** |
| Stage 3 | Areoles-Veins > Areoles-Tube | 0.38 (±0.04) | 10.03 | <0.001 | *** |
| Stage 3 | Areoles-Veins > Veins-Peristome | -0.13 (±0.04) | -3.48 | 0.0076 | ** |
| Stage 3 | Areoles-Veins > Veins-Tube | 0.3 (±0.04) | 7.71 | <0.001 | *** |
| Stage 3 | Areoles-Veins > Peristome-Tube | -0.68 (±0.05) | -13.87 | <0.001 | *** |

| | | | | | |
|---|---|---|---|---|---|
| Stage 3 | Areoles-Peristome > Areoles-Tube | 1.31 (±0.04) | 29.70 | <0.001 | *** |
| Stage 3 | Areoles-Peristome > Veins-Peristome | 0.79 (±0.04) | 17.99 | <0.001 | *** |
| Stage 3 | Areoles-Peristome > Veins-Tube | 1.22 (±0.04) | 27.65 | <0.001 | *** |
| Stage 3 | Areoles-Peristome > Peristome-Tube | 0.24 (±0.05) | 4.51 | <0.001 | *** |
| Stage 3 | Areoles-Tube > Veins-Peristome | -0.52 (±0.04) | -11.73 | <0.001 | *** |
| Stage 3 | Areoles-Tube > Veins-Tube | -0.09 (±0.04) | -1.99 | 0.7044 | |
| Stage 3 | Areoles-Tube > Peristome-Tube | -1.07 (±0.05) | -19.79 | <0.001 | *** |
| Stage 3 | Veins-Peristome > Veins-Tube | 0.43 (±0.04) | 9.71 | <0.001 | *** |
| Stage 3 | Veins-Peristome > Peristome-Tube | -0.55 (±0.05) | -10.20 | <0.001 | *** |
| Stage 3 | Veins-Tube > Peristome-Tube | -0.98 (±0.05) | -18.13 | <0.001 | *** |
| Stage 4 | Areoles-Veins > Areoles-Peristome | -0.59 (±0.07) | -8.67 | <0.001 | *** |
| Stage 4 | Areoles-Veins > Areoles-Tube | 0.3 (±0.07) | 4.43 | <0.001 | *** |
| Stage 4 | Areoles-Veins > Veins-Peristome | 0.22 (±0.07) | 3.24 | 0.0178 | * |
| Stage 4 | Areoles-Veins > Veins-Tube | 0.3 (±0.07) | 4.53 | <0.001 | *** |
| Stage 4 | Areoles-Veins > Peristome-Tube | -0.54 (±0.09) | -6.15 | <0.001 | *** |
| Stage 4 | Areoles-Peristome > Areoles-Tube | 0.88 (±0.08) | 11.34 | <0.001 | *** |
| Stage 4 | Areoles-Peristome > Veins-Peristome | 0.81 (±0.08) | 10.23 | <0.001 | *** |
| Stage 4 | Areoles-Peristome > Veins-Tube | 0.89 (±0.08) | 11.42 | <0.001 | *** |
| Stage 4 | Areoles-Peristome > Peristome-Tube | 0.05 (±0.1) | 0.51 | 1.0000 | |
| Stage 4 | Areoles-Tube > Veins-Peristome | -0.07 (±0.08) | -0.96 | 1.0000 | |
| Stage 4 | Areoles-Tube > Veins-Tube | 0.01 (±0.08) | 0.08 | 1.0000 | |
| Stage 4 | Areoles-Tube > Peristome-Tube | -0.84 (±0.1) | -8.72 | <0.001 | *** |
| Stage 4 | Veins-Peristome > Veins-Tube | 0.08 (±0.08) | 1.04 | 1.0000 | |
| Stage 4 | Veins-Peristome > Peristome-Tube | -0.76 (±0.1) | -7.86 | <0.001 | *** |
| Stage 4 | Veins-Tube > Peristome-Tube | -0.84 (±0.1) | -8.78 | <0.001 | *** |
| Stage 1 | *S.* x *leucophylla* > *S.* x Juthatip soper | 0.43 (±0.39) | 1.08 | 1.0000 | |
| Stage 1 | *S.* x *leucophylla* > *S.* x *mitchelliana* | 0.92 (±0.39) | 2.32 | 0.2077 | |
| Stage 1 | *S.* x *leucophylla* > *S. purpurea* | 0.87 (±0.39) | 2.22 | 0.2539 | |
| Stage 1 | *S.* x Juthatip soper > *S.* x *mitchelliana* | 0.49 (±0.39) | 1.24 | 1.0000 | |
| Stage 1 | *S.* x Juthatip soper > *S. purpurea* | 0.45 (±0.39) | 1.14 | 1.0000 | |
| Stage 1 | *S.* x *mitchelliana* > *S. purpurea* | -0.04 (±0.39) | -0.10 | 1.0000 | |
| Stage 2 | *S.* x *leucophylla* > *S.* x Juthatip soper | 0.49 (±0.23) | 2.15 | 0.2875 | |
| Stage 2 | *S.* x *leucophylla* > *S.* x *mitchelliana* | 0.38 (±0.23) | 1.68 | 0.6793 | |
| Stage 2 | *S.* x *leucophylla* > *S. purpurea* | 0.03 (±0.23) | 0.13 | 1.0000 | |
| Stage 2 | *S.* x Juthatip soper > *S.* x *mitchelliana* | -0.11 (±0.23) | -0.47 | 1.0000 | |
| Stage 2 | *S.* x Juthatip soper > *S. purpurea* | -0.46 (±0.23) | -2.02 | 0.3661 | |
| Stage 2 | *S.* x *mitchelliana* > *S. purpurea* | -0.35 (±0.23) | -1.55 | 0.8467 | |
| Stage 3 | *S.* x *leucophylla* > *S.* x Juthatip soper | 0.33 (±0.23) | 1.45 | 1.0000 | |
| Stage 3 | *S.* x *leucophylla* > *S.* x *mitchelliana* | -0.23 (±0.23) | -1.00 | 1.0000 | |
| Stage 3 | *S.* x *leucophylla* > *S. purpurea* | 0.4 (±0.23) | 1.74 | 0.6168 | |
| Stage 3 | *S.* x Juthatip soper > *S.* x *mitchelliana* | -0.56 (±0.23) | -2.45 | 0.1632 | |
| Stage 3 | *S.* x Juthatip soper > *S. purpurea* | 0.07 (±0.23) | 0.29 | 1.0000 | |
| Stage 3 | *S.* x *mitchelliana* > *S. purpurea* | 0.62 (±0.23) | 2.73 | 0.0920 | |
| Stage 4 | *S.* x *leucophylla* > *S.* x Juthatip soper | 0.81 (±0.39) | 2.06 | 0.3403 | |
| Stage 4 | *S.* x *leucophylla* > *S.* x *mitchelliana* | 0.74 (±0.4) | 1.87 | 0.4885 | |

| | | | | | |
|---|---|---|---|---|---|
| Stage 4 | *S.* x *leucophylla* > *S. purpurea* | 0.89 (±0.39) | 2.25 | 0.2391 | |
| Stage 4 | *S.* x Juthatip soper > *S.* x *mitchelliana* | -0.08 (±0.4) | -0.19 | 1.0000 | |
| Stage 4 | *S.* x Juthatip soper > *S. purpurea* | 0.07 (±0.39) | 0.19 | 1.0000 | |
| Stage 4 | *S.* x *mitchelliana* > *S. purpurea* | 0.15 (±0.4) | 0.38 | 1.0000 | |
| Areoles-Veins | Stage 1 > (2, 3, 4) | -0.57 (±0.16) | -3.53 | 0.0106 | * |
| Areoles-Veins | Stage 2 > (1, 3, 4) | 0.47 (±0.16) | 2.95 | 0.0378 | * |
| Areoles-Veins | Stage 3 > (1, 2, 4) | -0.02 (±0.12) | -0.19 | 1.0000 | |
| Areoles-Veins | Stage 4 > (1, 2, 3) | 0.12 (±0.16) | 0.75 | 1.0000 | |
| Areoles-Peristome | Stage 1 > (2, 3, 4) | -0.49 (±0.17) | -2.95 | 0.0332 | * |
| Areoles-Peristome | Stage 2 > (1, 3, 4) | -0.07 (±0.16) | -0.43 | 1.0000 | |
| Areoles-Peristome | Stage 3 > (1, 2, 4) | 0.43 (±0.12) | 3.59 | 0.0080 | ** |
| Areoles-Peristome | Stage 4 > (1, 2, 3) | 0.13 (±0.17) | 0.76 | 1.0000 | |
| Areoles-Tube | Stage 1 > (2, 3, 4) | -0.27 (±0.17) | -1.61 | 0.4971 | |
| Areoles-Tube | Stage 2 > (1, 3, 4) | -0.22 (±0.16) | -1.34 | 0.7884 | |
| Areoles-Tube | Stage 3 > (1, 2, 4) | 0.11 (±0.12) | 0.92 | 1.0000 | |
| Areoles-Tube | Stage 4 > (1, 2, 3) | 0.37 (±0.17) | 2.25 | 0.1474 | |
| Veins-Peristome | Stage 1 > (2, 3, 4) | -0.03 (±0.17) | -0.19 | 1.0000 | |
| Veins-Peristome | Stage 2 > (1, 3, 4) | -0.17 (±0.16) | -1.06 | 1.0000 | |
| Veins-Peristome | Stage 3 > (1, 2, 4) | 0.27 (±0.12) | 2.21 | 0.1590 | |
| Veins-Peristome | Stage 4 > (1, 2, 3) | -0.06 (±0.17) | -0.39 | 1.0000 | |
| Veins-Tube | Stage 1 > (2, 3, 4) | -0.57 (±0.16) | -3.43 | 0.0115 | * |
| Veins-Tube | Stage 2 > (1, 3, 4) | 0.22 (±0.16) | 1.36 | 0.7661 | |
| Veins-Tube | Stage 3 > (1, 2, 4) | 0.11 (±0.12) | 0.88 | 1.0000 | |
| Veins-Tube | Stage 4 > (1, 2, 3) | 0.24 (±0.17) | 1.46 | 0.6456 | |
| Peristome-Tube | Stage 1 > (2, 3, 4) | -0.59 (±0.18) | -3.37 | 0.0104 | * |
| Peristome-Tube | Stage 2 > (1, 3, 4) | -0.42 (±0.17) | -2.50 | 0.0861 | |
| Peristome-Tube | Stage 3 > (1, 2, 4) | 0.53 (±0.13) | 4.15 | 0.0015 | ** |
| Peristome-Tube | Stage 4 > (1, 2, 3) | 0.48 (±0.18) | 2.71 | 0.0481 | * |
| *S.* x *leucophylla* | Stage 1 > (2, 3, 4) | -0.18 (±0.31) | -0.60 | 1.0000 | |
| *S.* x *leucophylla* | Stage 2 > (1, 3, 4) | -0.23 (±0.24) | -0.97 | 1.0000 | |
| *S.* x *leucophylla* | Stage 3 > (1, 2, 4) | -0.1 (±0.22) | -0.47 | 1.0000 | |
| *S.* x *leucophylla* | Stage 4 > (1, 2, 3) | 0.52 (±0.31) | 1.70 | 0.4383 | |
| *S.* x Juthatip soper | Stage 1 > (2, 3, 4) | -0.06 (±0.31) | -0.21 | 1.0000 | |
| *S.* x Juthatip soper | Stage 2 > (1, 3, 4) | -0.2 (±0.24) | -0.83 | 1.0000 | |
| *S.* x Juthatip soper | Stage 3 > (1, 2, 4) | 0.14 (±0.22) | 0.66 | 1.0000 | |
| *S.* x Juthatip soper | Stage 4 > (1, 2, 3) | 0.12 (±0.31) | 0.40 | 1.0000 | |
| *S.* x *mitchelliana* | Stage 1 > (2, 3, 4) | -0.8 (±0.31) | -2.62 | 0.0771 | |
| *S.* x *mitchelliana* | Stage 2 > (1, 3, 4) | -0.14 (±0.24) | -0.59 | 1.0000 | |
| *S.* x *mitchelliana* | Stage 3 > (1, 2, 4) | 0.8 (±0.22) | 3.68 | 0.0088 | ** |
| *S.* x *mitchelliana* | Stage 4 > (1, 2, 3) | 0.14 (±0.31) | 0.46 | 1.0000 | |
| S. *purpurea* | Stage 1 > (2, 3, 4) | -0.62 (±0.31) | -2.03 | 0.2419 | |
| S. *purpurea* | Stage 2 > (1, 3, 4) | 0.46 (±0.24) | 1.87 | 0.3220 | |
| S. *purpurea* | Stage 3 > (1, 2, 4) | 0.1 (±0.22) | 0.46 | 1.0000 | |
| S. *purpurea* | Stage 4 > (1, 2, 3) | 0.07 (±0.31) | 0.22 | 1.0000 | |

**Table S5.** Factors of variation in the number of prey individuals, flying Hymenoptera individuals and crawling Hymenoptera individuals. Only the best models are presented. "Chisq" values refer to type III Wald chi-square tests. Symbols describe various levels of p-values *: p<0.05, **: p<0.01, and ***: p<0.001. Contrasts are detailed in Table 3.

| Dependent variable | Explanatory variables | Chisq | Df | p-value |
|---|---|---|---|---|
| Number of prey individuals | Intercept | 33.58 | 1 | <0.001 *** |
| | Plant taxon | 40.80 | 3 | <0.001 *** |
| | Pitcher stage | 86.04 | 3 | <0.001 *** |
| | Aperture width | 9.55 | 1 | 0.0020 ** |
| | Pitcher length | 45.67 | 1 | <0.001 *** |
| Number of flying Hymenoptera individuals | Season | 32.67 | 1 | <0.001 *** |
| | Pitcher length | 39.14 | 1 | <0.001 *** |
| | Areoles-Tube colour contrast | 16.77 | 1 | <0.001 *** |
| | Areoles brightness contrast | 7.83 | 1 | 0.0051 ** |
| | Areoles-Peristome colour contrast | 36.87 | 1 | <0.001 *** |
| Number of crawling Hymenoptera individuals | Intercept | 1.70 | 1 | 0.1925 |
| | Plant taxon | 80.69 | 3 | <0.001 *** |
| | Pitcher stage | 15.12 | 3 | 0.0017 ** |
